# Moderate High Temperature is Beneficial or Detrimental Depending on Carbon Availability in the Green Alga *Chlamydomonas reinhardtii*

**DOI:** 10.1101/2022.12.04.519034

**Authors:** Ningning Zhang, Benedikt Venn, Catherine E. Bailey, Ming Xia, Erin M. Mattoon, Timo Mühlhaus, Ru Zhang

## Abstract

High temperatures impair plant and algal growth and reduce food and biofuel production, but the underlying mechanisms remain elusive. The unicellular green alga *Chlamydomonas reinhardtii* is a superior model to study heat responses in photosynthetic cells due to its fast growth rate, many similarities in cellular processes to land plants, simple and sequenced genome, and ample genetic and genomics resources. Chlamydomonas grows in light by photosynthesis and/or with the externally supplied organic carbon source, acetate. Most of the published research about Chlamydomonas heat responses used acetate-containing medium. Understanding how organic carbon sources affect heat responses is important for the algal industry but understudied. We cultivated Chlamydomonas wild-type cultures under highly controlled conditions in photobioreactors at control of 25°C, moderate high temperature of 35°C, or acute high temperature of 40°C with and without constant acetate supply for 1- or 4-days. Our results showed that 35°C increased algal growth with constant acetate supply but reduced algal growth without sufficient acetate. The overlooked and dynamic effects of 35°C could be explained by induced carbon metabolism, including acetate uptake and assimilation, glyoxylate cycle, gluconeogenesis pathways, and glycolysis. Acute high temperature at 40°C for more than 2 days was lethal to algal cultures with and without constant acetate supply. Our research provides insights to understand algal heat responses and help improve thermotolerance in photosynthetic cells.

**Highlight:** We revealed the overlooked, dynamic effects of moderate high temperature in algae depending on carbon availability and demonstrated the importance of carbon metabolism in thermotolerance of photosynthetic cells.

## Introduction

Many biological processes in photosynthetic cells are sensitive to high temperatures (Mittler *et al*., 2012; Schroda *et al*., 2015; Mishra *et al*., 2021; Guihur *et al*., 2022). Global warming increases the intensity, duration, and frequency of high temperatures in the field, reducing plant growth and food production (Janni *et al*., 2020). A recent model revealed high temperature as the primary climatic driver for yield loss in crops from 1981-2017 in the U.S. (Ortiz-Bobea *et al*., 2019). Considering the increasing global temperature and human population, it is imperative to understand how photosynthetic cells respond to high temperatures and improve plant thermotolerance.

High temperatures in the field have different intensities and durations. For many land plants and algae grown in temperate regions, moderate high temperatures refer to heat slightly above the optimal temperature for growth (at or around 35°C), while acute high temperatures refer to heat at or above 40°C, which significantly reduces or completely halts growth (Zhang *et al*., 2022*a*). Moderate high temperatures are often long-lasting and frequent in nature with mild effects on photosynthetic cells, while acute high temperatures are often short-term but severely damaging. Most previous heat-stress experiments in plants and algae used acute high temperatures at or above 40°C (Hemme *et al*., 2014; Rütgers *et al*., 2017; Balfagón *et al*., 2019; Kim *et al*., 2020; Ji *et al*., 2021), likely due to the rapid onset and easily quantifiable phenotypes as compared to moderate high temperatures. Although the impact of moderate high temperatures in photosynthetic cells can be difficult to investigate due to comparatively mild phenotypes, moderate high temperatures are physiologically relevant stresses in field conditions. Additionally, global warming can further increase the frequency and duration of moderate high temperatures in nature. The frequent and long-lasting features of moderate high temperatures could reduce agricultural yield significantly (Delorge *et al*., 2014; Anderson *et al*., 2021). Arabidopsis seedlings treated with 5-day heat at 35°C had reduced growth and viability (Song *et al*., 2021). Furthermore, long-term (4-6 days) moderate high temperatures resulted in coral bleaching by causing cell death in endosymbiotic algae and their coral hosts (Dunn *et al*., 2004) and destabilizing the symbiotic nutrient cycling relationship between these two organisms (Rädecker *et al*., 2021). However, the effects of moderate high temperatures in photosynthetic cells and the interaction of moderate high temperatures with nutrient status especially carbon availability are underexplored. Algae have great potential for production of biofuels and bioproducts (Scranton *et al*., 2015; Mathimani and Pugazhendhi, 2019). However, how algal cells respond to high temperatures is under-investigated as compared to land plants (Schroda *et al*., 2015). To study the effects of carbon status, heat intensities and duration on heat responses in photosynthetic cells, we utilized the unicellular green alga *Chlamydomonas reinhardtii* (Chlamydomonas throughout).

Chlamydomonas is a superior model to study heat responses in photosynthetic cells (Schroda *et al*., 2015). Chlamydomonas can grow in light under photoautotrophic conditions using photosynthesis (Minagawa and Tokutsu, 2015). It can also grow with supplied organic carbon source (acetate) in light (mixotrophic) or dark (heterotrophic) conditions (Sasso *et al*., 2018), providing a platform to study heat responses under different light and carbon conditions. Additionally, it has a haploid, sequenced, annotated, small genome (111 Mb, 17,741 protein-encoding genes) with simpler gene families and fewer gene duplications than land plants (Merchant *et al*., 2007; Karpowicz *et al*., 2011). Furthermore, several well-established gene editing, cloning, high-throughput, and functional genomic tools are available in Chlamydomonas (Shimogawara *et al*., 1998; Li *et al*., 2016, 2019; Greiner *et al*., 2017; Crozet *et al*., 2018; Wang *et al*., 2019; Dhokane *et al*., 2020; Emrich-Mills *et al*., 2021).

Outdoor algal ponds frequently experience moderate high temperatures around 35°C during the summer (Mata *et al*., 2010; Krishnan *et al*., 2021), but the effects of moderate high temperatures on algal growth have been overlooked. Previously published algal heat treatments were often conducted in flasks in pre-warmed water baths at or above 42°C with sharp temperature switches (Hemme *et al*., 2014; Rütgers *et al*., 2017). While previous research was highly valuable to understanding algal heat responses, high temperatures in the field or outdoor ponds often increase gradually. Rapid increases to high temperatures largely reduce algal viability (Zhang *et al*., 2022*a*); similar effects have also been reported in land plants (Mittler *et al*., 2012). Acute high temperatures at or above 40°C inhibit algal cell division (Mühlhaus *et al*., 2011*b*; Hemme *et al*., 2014; Zachleder *et al*., 2019; Ivanov *et al*., 2021), while moderate high temperature at 35°C only transiently inhibits cell division during the first 4-8 hour (h) heat (Zhang *et al*., 2022*a*). Thus, conducting long-term experiments at moderate high temperatures in flasks can result in overgrown cultures, nutrient depletion, and light limitation, complicating data interpretation.

Consequently, it is advantageous to investigate algal heat responses under highly-controlled conditions in photobioreactors (PBRs) (Zhang *et al*., 2022*a*; Mattoon *et al*., 2022*a*), which have several evident strengths: (1) precise temperature regulation with a sterile temperature probe inside each algal culture; (2) controlled heating to mimic the heating speed in nature; (3) precisely controlled cultivation conditions, including temperature, light, nutrient, and air agitation, allowing for reproducible experiments; (4) automatic recording of culture status every minute, including growth conditions and optical densities, enabling quantitative growth rate measurements; (5) the availability of turbidostatic control based on defined parameters (e.g., chlorophyll contents) to enable frequent culture dilutions with constant medium supply; (6) being able to simulate nutrient depletion by turning off the turbidostatic control to investigate how nutrient availability affects algal heat responses. Utilization of PBRs for algal cultivation and heat treatments can largely reduce compounding effects during high temperature treatments and improve our understanding of algal heat responses.

Recently, we conducted systems-wide analyses in wild-type (WT) Chlamydomonas cultures during and after 24-h heat of 35°C and 40°C in PBRs under light in acetate-containing medium (mixotrophic condition) with turbidostatic control and constant nutrient supply (Zhang *et al*., 2022*a*). We chose moderate high temperature of 35°C and acute high temperature of 40°C based on the algal growth rates under different temperatures and their relevance to physiologically high temperatures experienced in nature (Krishnan *et al*., 2021). The 24-h heat duration allowed us to balance time-course sample collection over a sufficient duration while maintaining culture viability (heat at 40°C for longer time killed algal cells); our experimental conditions enabled us to investigate algal heat responses during and after physiologically relevant high temperatures (Zhang *et al*., 2022*a*). Our results showed that 40°C inhibited algal growth while 35°C increased algal growth with constant acetate supply. The growth inhibition at 40°C could be explained by reduced photosynthesis, impaired respiration, and arrested cell cycle, while these cell parameters had minor changes in algal cultures treated by 35°C. Our proteomics data indicated that several proteins involved in acetate uptake and assimilation, glyoxylate cycle, and gluconeogenesis were up-regulated during 35°C (Zhang *et al*., 2022*a*). Chlamydomonas uptakes acetate and feeds it into the glyoxylate cycle and gluconeogenesis for starch biosynthesis (Johnson and Alric, 2012, 2013). The main function of glyoxylate cycle is assimilation of 2-carbon compounds (e.g., acetate) into organic acids that can be metabolized by the cell; the glyoxylate cycle is a shunt of the tricarboxylic acid (TCA) cycle in mitochondria without releasing CO_2_, which allows for the anabolism of simple carbon compounds in gluconeogenesis, a process to make sugars, e.g., glucose (Johnson and Alric, 2012; Chew *et al*., 2019; Walker *et al*., 2021).

We hypothesized that the increased growth in Chlamydomonas at 35°C with constant acetate supply was attributable to up-regulated acetate metabolism. In our previous papers, we found out that 35°C increased the growth rates of Chlamydomonas cultures with constant acetate supply (Zhang *et al*., 2022*a*), but not in cultures without acetate (Mattoon *et al*., 2022*b*), as compared to the control at 25°C with the same medium type. Therefore, we hypothesized that the effects of moderate high temperature at 35°C was dynamic depending on the availability of acetate, e.g., constant-acetate, acetate-depleting, or no-acetate conditions. The majority of published high temperature research in Chlamydomonas has been conducted in acetate-containing medium (Voß *et al*., 2010; Mühlhaus *et al*., 2011*a*; Hemme *et al*., 2014; Rütgers *et al*., 2017; Zhang *et al*., 2022*a*), but the interface between acetate supply and Chlamydomonas heat responses is understudied. To validate our hypothesis and address these unknown questions, we cultivated the same Chlamydomonas WT strain in PBRs as before but heated the cultures at 35°C or 40°C with and without constant acetate supply for 1 or 4 days. Our previous work was conducted at 35°C and 40°C for 24 h with constant acetate supply (Zhang *et al*., 2022*a*). In this paper, to investigate the effects of carbon availability on algal thermotolerance, we included 3 new different heat treatments at 35°C and 40°C: (1) acetate-depleting condition for 24 h, (2) constant-acetate condition for 4 days, and (3) no-acetate condition for 4 days. The last two conditions utilized turbidostatic control with constant nutrient supply. Our new results confirmed our hypothesis about the role of acetate metabolism in algal thermotolerance and revealed the overlooked effects of moderate high temperature of 35°C on algal growth. Heat of 35°C was beneficial or detrimental to Chlamydomonas cultures depending on acetate availability. Acute high temperature of 40°C was lethal to Chlamydomonas cultures, with and without constant acetate supply. Our research helps fill the knowledge gaps about how external carbon supply affects algal heat responses, provides new insights to understand algal heat responses, has potential applications in production of algal biofuel and bioproducts, and contributes to improved thermotolerance in photosynthetic cells.

## Materials and methods

### Algal cultivation

The *Chlamydomonas reinhardtii* wild-type strain CC-1690 (also called *21gr*, mating type plus) (Sager, 1955; Pröschold *et al*., 2005; Zhang *et al*., 2022*b*) was purchased from the Chlamydomonas resource center and used in all experiments. Algal cultures were grown in Tris-acetate-phosphate (TAP, with acetate) or Tris-phosphate (TP, without acetate) medium with revised trace elements (Kropat *et al*., 2011) in 400 mL photobioreactors (PBRs) (Photon System Instruments, FMT 150/400-RB) as described previously (Zhang *et al*., 2022*a*). Each medium type was used in individual PBRs for independent experiments. Cultures were illuminated with constant 100 µmol photons m^−2^ s^−1^ light (red: blue, 1:1 ratio), and agitated by bubbling with filtered air at a flow rate of 1 L min^−1^. Algal cultures with targeted cell density around 1-2 ×10^6^ cells mL^−1^ (∼4-6 µg mL^−1^ chlorophyll content) were maintained at 25°C for 4 days with constant nutrient supply through turbidostatic control before different temperature treatments. The turbidostatic mode was controlled by OD_680_ (optical density at 680 nm), which is proportional to chlorophyll contents (µg mL^−1^) and was monitored once per min automatically. When OD_680_ increased to a maximum value slightly above the target value (for target cell density) due to algal growth, OD_680_ signaled the control computer to turn on a turbidostatic pump to add fresh medium to dilute the culture until a minimum OD_680_ slightly below the target value was reached, then the turbidostatic pump was turned off automatically. Because of the small OD range we used, the PBR cultures had exponential growth between dilution events through the turbidostatic control with constant nutrient supply (Fig. 1A). The OD_680_ data during exponential growth phases in between dilution events was log_2_ transformed, and the relative growth rate was calculated using the slope of log_2_(OD_680_). The relative growth rate is the inverse of the algal culture doubling time. For the treatments without constant nutrient supply (acetate-depleting condition), the turbidostatic pumps were turned off after PBR cultures reached steady growth and at the start of different temperature treatments (Fig. 1B).

**Fig. 1.**
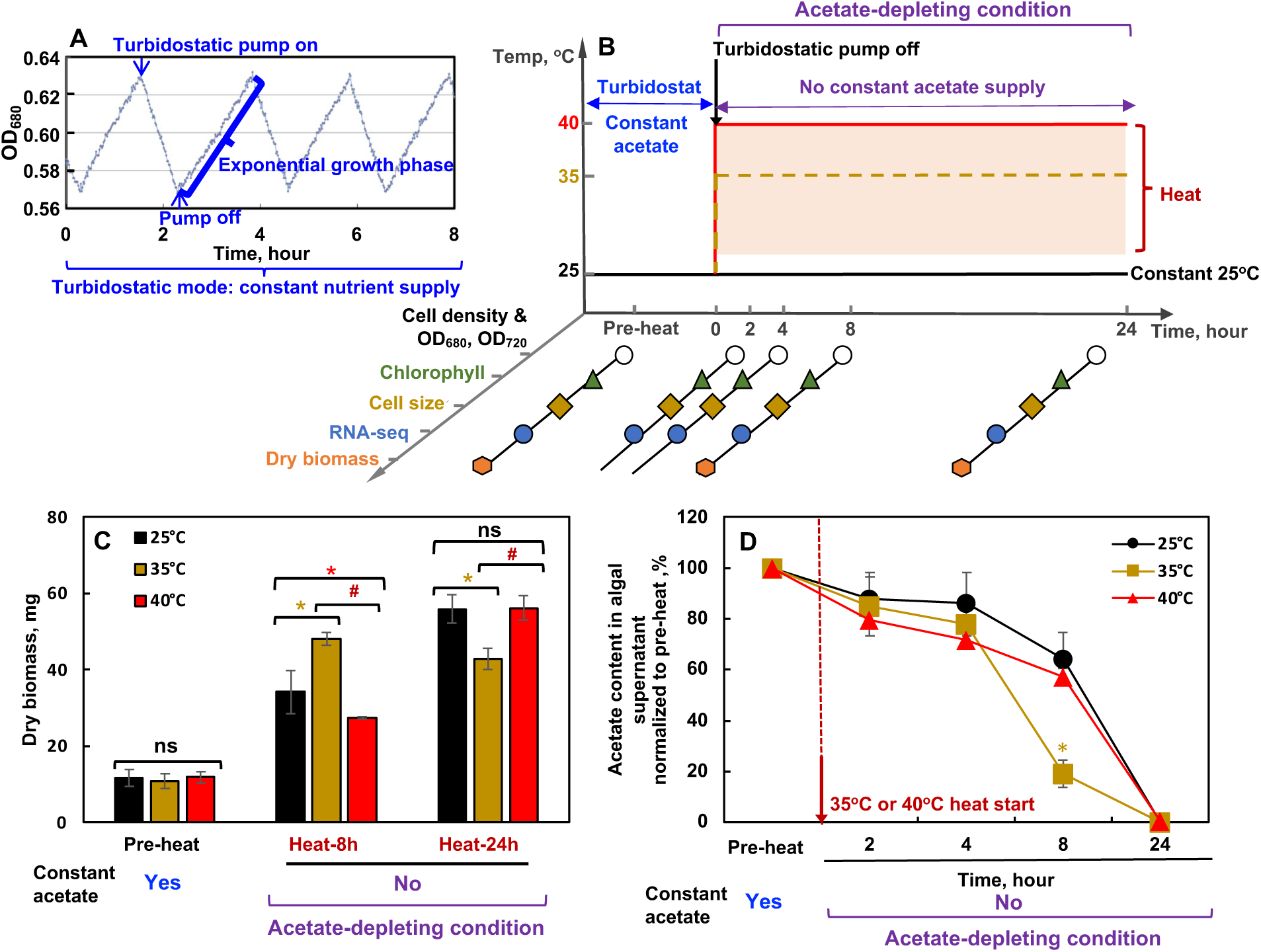
Under the acetate-depleting condition, moderate high temperature of 35°C transiently increased but then decreased Chlamydomonas biomass accumulation. **(A)** Before heat treatments, Chlamydomonas cells (CC-1690, also called *21gr*, wild-type) were grown in photobioreactors (PBRs) in Tris-acetate-phosphate (TAP) medium (acetate as an organic carbon source) with a light intensity of 100 μmol photons m^−2^ s^−1^ and constant bubbling of air. Algal cultures were maintained turbidostatically with frequent medium dilution using a small range of OD_680_, which monitors chlorophyll content. Figure cited and modified from a supplemental figure of our previous paper (Zhang *et al*., 2022*a*). **(B)** Experimental outline and sampling time points for the acetate-depleting condition. **(C)** Under the acetate-depleting condition, algal dry biomass (in 90 mL cultures) increased with 8-h heat but decreased with 24-h heat of 35°C as compared to the constant 25°C. Mean ± SE, *n*=3-8 biological replicates. **(D)** Heat of 35°C accelerated acetate consumption. Acetate content in algal supernatant was quantified using the Acetate Colorimetric Assay Kit. The red dashed line marks the start of heat or the corresponding time point at constant 25°C. **(C, D)** Statistical analyses were performed using two-tailed t-test assuming unequal variance; *, p<0.05, the colors of the asterisks match the heated condition as compared to 25°C at the same time points; #, p<0.05, for the comparisons between 35°C and 40°C at the same time points. Not significant, ns.

### High temperature treatments in PBRs

After algal cultures in PBRs reached steady growth with turbidostatic control at 25°C for 4 days, PBR temperatures were increased to moderate (35°C) or acute (40°C) high temperatures in different PBRs for the indicated duration with indicated nutrient conditions. PBR temperatures were increased from 25°C to 35°C or 40°C gradually over the course of 30 min with controlled heating speeds. PBR cultures grown under constant 25°C with the same nutrient status and treatment duration served as controls. Algal cultures were harvested from PBRs at different time points during different treatments for various measurements.

### Algal biomass quantification

Algal cultures (90 mL) were harvested from PBRs, centrifuged to remove supernatant, flash frozen in liquid nitrogen, then stored in a −80°C freezer until analysis. Algal cell pellets were freeze dried for 24 h over liquid nitrogen. The remaining algal dry biomass were weighed and quantified.

### Acetate quantification assay

Algal cultures were harvested from PBRs (2 mL) and centrifuged to collect 500 μL top clear supernatant to a new tube. The supernatant was stored in a −80°C freezer until analysis. The acetate content in the supernatant was quantified using the Acetate Colorimetric Assay Kit (Sigma, Cat No. MAK086) according to manufacture instructions.

### Cell imaging using light microscopy

Chlamydomonas cultures harvested at select time points with different temperature treatments were fixed with 0.2% glutaraldehyde (VWR, Cat No. 76177-346). Algal cells were imaged with a Leica DMI6000 B microscope and a 63x (NA1.4) oil-immersion objective.

### Chlorophyll quantification

Chlorophyll contents in algal cells were quantified as previously described (Zhang *et al*., 2022*a*). PBR cultures of 1 mL were harvested in 1.5-mL tubes with 2.5 μL 2% Tween20 (Sigma, P9416-100ML, to help cell pelleting), and centrifuged at 18,407 g at 4°C. After removing supernatant, cell pellets were stored in a −80°C freezer until quantification. Later, cell pellets were thawed, resuspended in 1 mL of HPLC grade methanol (100%, Sigma, 34860-4L-R), vortexed for 1 min, incubated in the dark at 4°C for 5 min, and centrifuged at 15,000 g at 4°C for 5 min. Supernatant was pipetted out to a two-sided disposable plastic cuvettes (VWR, 97000-586) for chlorophyll (Chl) quantification at 652 nm and 665 nm in a spectrophotometer (IMPLEN Nonophotometer P300) using the following equations: Chl a + Chl b = 22.12*A_652_ + 2.71*A_665_ (in μg mL^−1^ algal cultures) (Zhang *et al*., 2022*a*). Chlorophyll concentrations were also normalized to cell densities (Chl pg cell^−1^) or mean cell volume (Chl pg μm^−3^). Cell density and mean cell volume were measured using a Coulter Counter (Multisizer 3, Beckman Counter, Brea, CA). For 35°C with photoautotrophic medium, algal cells formed palmelloids, clumps of non-motile cells, thus the size of palmelloids instead of cell size was measured by the Coulter Counter.

### RNA extraction

PBR cultures of 2 mL were pelleted with Tween-20 (0.005%, v/v) by centrifugation at 11,363 x g and 4°C for 2 min, followed by supernatant removal. The cell pellet was flash frozen in liquid nitrogen and then stored in a −80°C freezer until processing. RNA extraction and RT-qPCR analysis were performed as previously described with minor modifications (Zhang *et al*., 2022*a*). Total RNA was extracted with TRIzol reagent (Thermo Fisher Scientific, Cat No. 15596026), digested on-column with RNase-free DNase (Qiagen, Cat No. 79256), purified by RNeasy mini-column (Qiagen, Cat No. 74106), and quantified with Qubit™ RNA BR Assay Kit, (Life technology, Cat No. Q10210).

### RT-qPCR analysis

Total 0.4 μg RNA was reverse transcribed with oligo dT primers using SuperScript® III First-Strand Synthesis System (Life technology, Cat No. 18080-051) following manufacturer instructions. Quantitative real-time PCR (RT-qPCR) analysis was performed using a CFX384 Real-Time System (C 1000 Touch Thermal Cycler, Bio-Rad, Hercules, California) using SensiFAST SYBR No-ROS kit (Bioline, BIO-98020) following this program: (1) 95°C (2 min); (2) 40 cycles of 95°C (5 s), 60°C (10 s), and 72°C (15 s); (3) final melt curve at 60°C (5 s), followed by continuous temperature ramping from 60°C to 99°C at a rate of 0.5°C s^−1^. Melting curves and qPCR products were tested to ensure there were no primer dimers or unspecific PCR products. All RT-qPCR products were sequenced for confirmation. Primers and gene IDs for RT-qPCR can be found in Supplementary Table 1. *CBLP* (*β-subunit-like polypeptide*, Cre06.g278222) (Schloss, 1990; Xie *et al*., 2013) and *EIF1A* (*Eukaryotic translation initiation factor 1A,* Cre02.g103550) (Strenkert *et al*., 2019) had stable expression among all time points, and were used as reference genes for RT-qPCR normalization. The relative gene expressions were calculated relative to the pre-heat time point by using the 2−^ΔΔCT^ method (Livak and Schmittgen, 2001; Hellemans *et al*., 2007; Remans *et al*., 2014). Three biological replicates were included for each time point and treatment. RT-qPCR was used to verify some selected transcripts and compared with the results of RNA-seq for samples harvested under the acetate-depleting conditions (Supplemental Fig. 2). RNA-seq for samples harvested under the constant-acetate conditions were verified by RT-qPCR previously (Zhang *et al*., 2022*a*).

### RNA-seq analysis

Raw sequencing data for the constant-acetate experiments at 35°C and 40°C were obtained from Zhang *et al*., 2022*a*. Raw sequencing data for the constant-acetate experiment at 25°C and acetate-depleting experiments at 25°C, 35°C, and 40°C were collected in this study. All raw sequencing data were processed using the same pipeline as described below.

RNA libraries were prepared and sequenced by the Joint Genome Institute (JGI, Community Science Program) using the NovaSeq platform generating 151-nt paired-end reads. Plate-based RNA sample preparation was performed on the PerkinElmer Sciclone NGS robotic liquid handling system using Illumina’s TruSeq Stranded mRNA HT sample prep kit utilizing poly-A selection of mRNA following the protocol outlined by Illumina in their user guide, with the following conditions: total RNA starting material was 1 μg per sample and 8 cycles of PCR was used for library amplification. The prepared libraries were quantified using KAPA Biosystems’ next-generation sequencing library qPCR kit and ran on a Roche LightCycler 480 real-time PCR instrument. Sequencing of the flowcell was performed on the Illumina NovaSeq sequencer using NovaSeq XP V1.5 reagent kits, S4 flowcell, following a 2×151 indexed run recipe.

Samples were quality control filtered using the JGI BBDuk and BBMap pipelines (BBDuk, 2022). Samples were quality assessed using FastQC (Andrews) and mapped to the *Chlamydomonas reinhardtii* v5.6 genome (Merchant *et al*., 2007) using HISAT2 version 2.2.0 (Kim *et al*., 2015). Reads per feature were counted via featureCounts (Liao *et al*., 2014). The count matrix was combined with existing data (Zhang *et al*., 2022*a*), resulting in a 16,403×101 count matrix (Supplemental Dataset 4, 5). The normalization of the count matrix was conducted using the median of ratios method (Liao *et al*., 2014). Prior to imputation of two missing time points in the constant-acetate-25°C condition (see details below), transcripts were filtered to have nonzero counts in at least 90% of the samples.

Under constant 25°C, acetate, and light with turbidostatic control (constant-acetate-25°C), the algal growth rate and cell physiology stays constant (Fig. 5). We had RNA-seq data for 8 time points (pre-heat, 0, 0.5, 1, 2, 24, 26, and 48 h) under constant-acetate-25°C and these samples were highly correlated with each other (with Pearson correlation ≥ 0.993) (Supplemental Fig. 1). RNA samples for the time points of 4 and 8 h under the constant-acetate-25°C condition were unavailable; thus RNA-seq data for the 4 and 8 h under the constant-acetate-25°C condition were imputed from other extra time points (0, 0.5, 1, 26, 48 h) of the same condition. For the time points of 4 and 8 h under the constant-acetate-25°C condition, each transcript triplicates were sampled randomly using the 5 extra time points under the same condition mentioned above. It was ensured that both missing time points were not imputed with the same count data. The final count matrix consists of 14,893 transcripts in 90 samples (5 time points measured at 6 conditions as triplicates). The natural logarithm was determined for all normalized counts before averaging triplicates. Pearson correlation coefficients were calculated for all sample pairs.

The count data were tested for differential expression using DESeq2 v1.38.3 (Love *et al*., 2014). DESeq2 applies a Benjamini Hochberg correction for multiple testing. The design matrix was constructed to test for global kinetic differences between the three different temperatures and both growth media over time. The following tests were performed (Supplemental Dataset 1, 3, 5, Supplemental Fig. 5): (A) acetate-depleting 25°C vs acetate-depleting 35°C; (B) acetate-depleting 25°C vs acetate-depleting 40°C; (C) constant-acetate 25°C vs constant-acetate 35°C; (D) constant-acetate 25°C vs constant-acetate 40°C; (E) temperature effect: (constant-acetate 35°C and acetate-depleting 35°C) vs (constant-acetate 40°C and acetate-depleting 40°C); (F) medium effect: (constant-acetate 35°C and constant-acetate 40°C) vs (acetate-depleting 35°C and acetate-depleting 40°C); (G) acetate-depleting 25°C vs constant-acetate 25°C.

Functional descriptions were determined for each transcript (Merchant *et al*., 2007; Usadel *et al*., 2009; Venn and Mühlhaus, 2022). Transcripts were grouped according to their functional description. Time courses of normalized transcript counts were visualized as log_2_(fold change). An ANOVA was performed for each transcript at each treatment to elucidate whether a transcript underwent a relevant change during its time course (Venn et al., 2022a). Heatmaps (Supplemental Dataset 1) and functional sets figures (Supplemental Dataset 2) were created using Plotly.NETv4.0.0 (Schneider *et al*., 2022).

Ontology enrichment was performed using extended MapMan annotations (for example “PS.lightreaction.LHC” became “PS.Lightreaction.LHC”, “PS.Lightreaction” and “PS”) (Supplemental Dataset 3). The measured and filtered transcripts served as background. If a transcript showed significant differential expression in the respective comparison (FDR < 0.05), it was considered as significant for its annotation. Enrichment p values were determined using hypergeometric tests. Multiple testing correction was performed using the Benjamini-Hochberg method (Benjamini and Hochberg, 1995; Venn *et al*., 2022). Additional ontology enrichment was performed using ChlamyCyc annotations (v2023-01-04). This pathway ontology combines KEGG, MapMan, and JGI pathway information (May *et al*., 2009). The enrichment procedure and statistical analysis were the same as above.

## Results

To investigate how the availability of an organic carbon source, acetate, affected algal heat responses, we first cultivated WT Chlamydomonas cells (CC-1690, *21gr*) in PBRs in Tris-acetate-phosphate (TAP, acetate as an organic carbon source) medium at 25°C with constant nutrient supply through turbidostatic control (providing frequent fresh medium for culture dilution) (Fig. 1A). After algal cultures reached steady growth rates in PBRs, the turbidostatic control was turned off and the temperatures of algal cultures were increased to 35°C or 40°C or stayed at 25°C for 24 h without constant acetate supply, defined as the acetate-depleting condition. This experiment allowed us to quantify acetate consumption under different temperatures, which could not be performed in cultures with turbidostatic control and frequent medium dilution. It also enabled us to investigate algal heat responses under the acetate-depleting condition, like shaker cultures, which were used by most previous algal heat experiments in published literatures (Hemme *et al*., 2014; Rütgers *et al*., 2017). Under the acetate-depleting condition, algal cultures were harvested at different time points to analyze cell physiology, biomass, and transcriptome (Fig. 1B). Algal dry biomass quantification showed that cultures treated with 35°C had increased biomass at 8-h heat but decreased biomass at 24-h heat as compared to 25°C (Fig. 1C). Acetate quantification in the supernatant of algal cultures showed that 35°C-treated cultures had increased acetate consumption and depleted acetate faster than 25°C or 40°C (Fig. 1D).

We hypothesized the transiently increased and then decreased algal biomass in 35°C-treated cultures was due to increased acetate uptake and usage which led to acetate starvation at 24 h of the acetate-depleting condition (Fig. 1). Chlamydomonas uptakes acetate and feeds it into the glyoxylate and gluconeogenesis cycles to produce starch, which can be used in glycolysis to generate ATP (Johnson and Alric, 2012, 2013) (Fig. 2A). To verify this hypothesis, we performed RNA-seq analysis and investigated key transcripts involved in acetate uptake and assimilation, glyoxylate, gluconeogenesis, and glycolysis cycles under the acetate-depleting condition by comparing with the constant-acetate condition (Fig. 2B-P). Under the acetate-depleting condition, almost all these transcripts related to acetate uptake and metabolism were up-regulated under 35°C as compared to 25°C, especially the early heat time points of 2 and 4 h at 35°C. Similar 35°C effects on these transcripts were held under the constant-acetate condition, although the up-regulation of these transcripts were often stronger and lasted longer at 35°C under the constant-acetate condition than the acetate-depleting condition. In contrast, these transcripts related to acetate uptake and metabolism were down-regulated under 40°C as compared to 25°C under both the acetate-depleting and constant-acetate conditions, especially the 2-h heat time point at 40°C. Our RT-qPCR results of selected transcripts showed similar trends as our RNA-seq data (Supplemental Fig. 2). These results showed that 35°C promoted acetate metabolism, from acetate uptake and assimilation to glyoxylate, gluconeogenesis, and glycolysis cycles; however, these acetate metabolism pathways were reduced under 40°C.

**Fig. 2.**
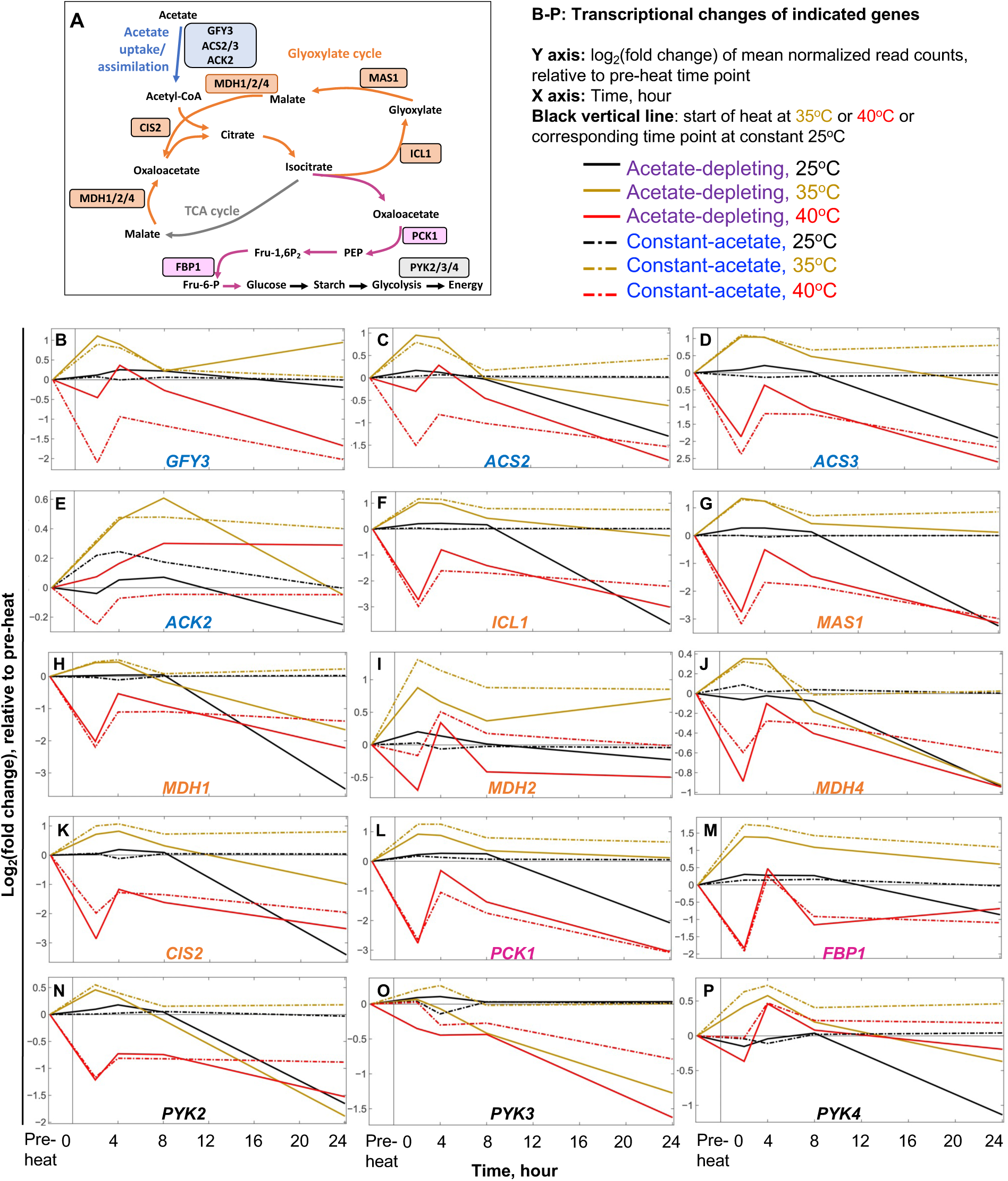
Moderate high temperature of 35°C up-regulated transcripts related to acetate uptake and assimilation, glyoxylate cycle, gluconeogenesis, and glycolysis pathways. **(A)** Simplified pathways of acetate metabolism in Chlamydomonas based on these papers (Johnson and Alric 2013; Durante et al. 2019). Key enzymes are in boxes and were analyzed via RNA-seq in panels B-P. Acetate uptake/assimilation enzymes: GFY3, putative acetate transporter; ACS2/3, acetyl-CoA synthases; ACK2, acetate kinase. Glyoxylate cycle key enzymes: ICL1, isocitrate lyase; MAS1, malate synthase; CIS2, Citrate synthase; MDH1/2/4, NAD-dependent malate dehydrogenases. Gluconeogenesis key enzymes: PCK1, phosphoenolpyruvate carboxykinase; FBP1, fructose-1,6-bisphosphatase. Glycolysis key enzymes: PYK2/3/4, pyruvate kinase 2/3/4. See all gene IDs and annotations in Supplemental Table 1. PEP, phosphoenolpyruvate. Fru-6-P, fructose-6-phosphate. Fru-1,6-P_2_, fructose-1,6-bisphosphate. **(B-P)** Transcriptional change of the indicated genes under six different conditions, presented as log_2_(fold change) of mean normalized read counts relative to the pre-heat time point. In each panel, each line represents the transcriptional time course of the indicated gene under one condition as indicated by the legend. The black vertical lines mark the start of heat or the corresponding time point at constant 25°C. Solid lines for the acetate-depleting condition and dashed lines for the constant-acetate condition. Black, brown, red lines for 25°C, 35°C, and 40°C temperature treatments, respectively. See Supplemental Dataset 1 for bigger pictures and more information.

In addition to the acetate metabolism pathways mentioned above, we investigated transcripts related to other processes that are often affected by high temperatures (Supplemental Fig. 3, 4). Chlamydomonas has two heat shock transcription factors, HSF1 and HSF2; HSF1 is the master heat regulator but the function of HSF2 is unclear (Schulz-Raffelt *et al*., 2007). The transcripts of both *HSF*s were induced by heat, with higher induction at 40°C than 35°C under the same acetate condition and higher induction under the acetate-depleting than constant-acetate condition under the same high temperature (Supplemental Fig. 3A, B). Heat shock proteins (HSPs) are the hallmarks of heat responses and function as chaperones to help refold denatured proteins, fold nascent polypeptides, and prevent denatured protein aggregates (Altschuler and Mascarenhas, 1982; Schroda *et al*., 2015; Haq *et al*., 2019). Based on molecular mass, HSPs are grouped into large HSPs (HSP100/90/70/60, >50 kDa) and small HSPs (10-43 kDa) (Jee, 2016; Bourgine and Guihur, 2021). Under the acetate-depleting condition, many transcripts related to large HSPs, *HSP60/70/90s*, were down-regulated at 24 h of 25°C and 35°C; such down-regulation of *HSP60/70/90s* was absent at 24 h of 25°C and 35°C under the constant-acetate condition or 24 h of 40°C under both acetate conditions (Supplemental Fig. 4A-C). Additionally, transcripts of *HSP100* and most of the small *HSPs* did not have such down-regulation at 24 h of 25°C and 35°C under the acetate-depleting condition (Supplemental Fig. 4D, E). Almost all transcripts related to the *HSPs* we examined, including *HSP60/70/90/100s* and small *HSPs*, were up-regulated at 2 h of 40°C, with similar induction fold change under either the acetate-depleting or constant-acetate condition (Supplemental Fig. 4A-E). This suggested the early induction of these *HSPs* under 40°C was temperature-dependent (present at 40°C while much less so under 35°C) but acetate-availability-independent. Alternatively, the acetate-availability might not differ significantly at 2 h of the acetate-depleting and constant-acetate condition (Fig. 1D). However, the absence of down-regulated *HSP* transcripts at 24 h of 40°C under the acetate-depleting condition supported that the induced *HSP* transcripts at 40°C was independent of acetate.

The global transcriptome changes related to light reaction and Calvin-Benson cycle of photosynthesis suggested more disturbance of these transcripts under the acetate-depleting than the constant-acetate conditions at the same temperatures, and more disturbance of these transcripts at 40°C than 35°C under the same acetate conditions (Supplemental Fig. 4F-M). These photosynthesis-related transcripts behaved similarly at 40°C under both the acetate-depleting and constant-acetate conditions, suggesting 40°C heat effects were more dominant than the changes of acetate availability. Chlamydomonas employs a cellular structure, pyrenoid, to concentrate CO_2_ for improved carbon fixation efficiency, called carbon concentrating mechanisms (CCM) (Meyer *et al*., 2017; Hennacy and Jonikas, 2020). The CCM is especially important under high temperature conditions, where the solubility of CO_2_ decreases in water by approximately 30% from 25°C to 40°C (Dodds *et al*., 1956). Under the acetate-depleting conditions, several transcripts related to CCM were up-regulated at 24 h under 25°C and 35°C, but not under the same temperature of the constant-acetate condition (Supplemental Fig. 4N). This suggested that as acetate became depleted in the medium and cell density increased, cells relied more on the photosynthetic carbon fixation by up-regulating CCM. Such induction of the CCM transcripts was much less at 40°C under the acetate-depleting condition, which may correlate with the compromised pyrenoid structures with decreased appearance of thylakoid tubules at 40°C in our previous publication (Zhang *et al*., 2022*a*). Thylakoid tubules deliver concentrated CO_2_ to the pyrenoid and also serve as the diffusion path of Calvin-Benson Cycle metabolites (Hennacy and Jonikas, 2020). Transcripts related to photoprotection had larger induction at 24 h under all three temperatures of the acetate-depleting than the constant-acetate conditions at the same temperature, indicating compromised photosynthesis and the need for photoprotection when acetate was depleted (Supplemental Fig. 4O).

The compromised photosynthesis under high temperatures lead to increased accumulation of reactive oxygen species (ROS) (Pospíšil, 2016; Janni *et al*., 2020; Niemeyer *et al*., 2021). Some transcripts related to ROS scavenging were down-regulated at 24 h of 25°C and 35°C under the acetate-depleting condition but much less so under the constant-acetate condition at the same temperatures (Supplemental Fig. 4R-V). This may be related to the increased cell density and reduced light and oxidative stress at 24 h of 25°C and 35°C under the acetate-depleting condition. Such down-regulation of transcripts related to ROS scavenging was not present at 24 h of 40°C under the acetate-depleting condition. This may suggest relatively increased ROS accumulation and ROS scavenging activity at 40°C than 25°C and 35°C under the acetate-depleting conditions. Additionally, transcripts related to ROS scavenging had very similar transcriptional changes at 40°C under either the acetate-depleting or constant-acetate condition, suggesting 40°C had more dominant effects on the ROS accumulation and ROS scavenging activities than acetate.

Transcripts related to cell cycle and cell wall had similar transcriptional patterns under each of the six conditions (Supplemental Fig. 4W, X). Under diurnal cycles, cell wall genes were up-regulated following the up-regulation of cell cycle genes and cell walls form around daughter cells during cell division (Zones *et al*., 2015). These transcripts were down-regulated at 24 h of 25°C under the acetate-depleting condition, suggesting the transition to reduced cell division due to the acetate-depletion. The reduction of these transcripts was larger and more prolonged under 40°C than 35°C, consistent with complete and transient inhibition of cell cycle under 40°C and 35°C, respectively (Hemme *et al*., 2014; Zhang *et al*., 2022*a*). At 24 h of 40°C and 35°C, these transcripts were more reduced under the acetate-depleting condition than the constant-acetate condition. It is worthwhile to note that the changes during 24 h of the acetate-depletion condition were dynamic and here we just mentioned the changes at 24 h in the end when acetate was depleted.

Almost all transcripts mentioned above had little changes at 25°C during all time points of the constant-acetate condition, validating the steady growth and constant cell status under the turbidostatic control with constant nutrient and light but without heat treatments in our algal cultures grown in PBRs (Supplemental Fig. 4).

We performed statistical analysis to investigate global transcript kinetic differences between different conditions (Supplemental Dataset 1, 3). We then applied ontology enrichments analysis for transcripts that were significantly expressed in a defined comparison to identify differentially regulated biological processes (Supplemental Fig. 5, Supplemental Dataset 3). The top enriched function group that differed at 25°C and 35°C under both acetate-depleting and constant-acetate condition was the nuclear-encoded ribosomal proteins that are involved in chloroplast protein synthesis (Supplemental Fig. 5A, C). More than half of the ribosomal related transcripts were down-regulated at 4 h heat of 35°C as compared to 25°C under both acetate conditions (Supplemental Fig. 4Y, Z), suggesting 35°C may inhibit chloroplast protein synthesis. The down-regulation of ribosomal related transcripts at 35°C could recover partially under the constant-acetate condition, but much less so under the acetate-depleting condition. The top enriched function group that differs between 25°C and 40°C under the acetate-depleting condition was the mitochondrial electron transport (Supplemental Fig. 5B). Many transcripts related to mitochondrial electron transport were down-regulated at 2 h heat of 40°C as compared to 25°C under the acetate-depleting condition, especially these related to ATP synthesis (Supplemental Fig. 4AA, AB). When comparing 35°C and 40°C conditions independent of the acetate availability, the top enriched function group was also the mitochondrial electron transport (Supplemental Fig. 5E). The results suggested that mitochondrial electron transport was more sensitive to 40°C than 25°C and 35°C. Under the constant-acetate condition, 40°C affected transcripts related to RNA processing the most as compared to 25°C (Supplemental Fig. 5D, 4AC, AD). When comparing the acetate-depleting and constant-acetate conditions independent of temperatures, the top two enriched groups were phosphate transport and mitogen-activated protein kinase (MAP kinase) cascades signaling (Supplemental Fig. 5F). MAP kinase signaling pathways have important roles in plant growth and stress responses and involve protein phosphorylation and dephosphorylation (Bigeard and Hirt, 2018; Jagodzik *et al*., 2018; Zhang and Zhang, 2022). The biggest differences in transcripts related to the MAP kinase signaling pathways between the acetate-depleting and constant-acetate conditions were at the 24 h time point at 25°C and 35°C when the acetate was depleted (Supplemental Fig. 4AF), suggesting potential roles of the MAP kinase signaling pathways in nutrient sensing and growth regulation. When comparing the acetate-depleting and constant-acetate conditions at 25°C, the top enriched functional groups are dominantly related to cell motility (Supplemental Fig. 5G). More than half of the transcripts related to cell motility at 25°C were down-regulated at 24 h under the acetate-depleting condition but not the constant-acetate condition (Supplemental Fig. 4AG, AH). Microscopic images showed that cells had cilia at 24 h of 25°C under the acetate-depleting condition (Supplemental Fig. 6). Cell motility needs energy and ciliary beating frequency corelates with the cytosolic ATP concentration in Chlamydomonas (Takano *et al*., 2021). Thus, the down-regulation of transcripts related to cell motility may suggest the reduced cellular ATP level and the transition from motile to non-motile phase when the acetate was depleted at 24 h of the acetate-depleting condition.

Besides MapMan ontology enrichment, we performed ChlamyCyc pathway enrichment analysis of differentially regulated transcripts compared between two conditions (Supplemental Dataset 6). The ChlamyCyc annotations (v2023-01-04) and pathway ontology combine KEGG, MapMan, and JGI pathway information (May et al. 2009). The KEGG pathway annotation for Chlamydomonas is sparse, therefore it was difficult to determine significances. As comparison, for the MapMan functional ontology we got 23,500 annotations in the RNA-seq data, for KEGG pathways we only got 803. Both analyses showed the mitochondrial activity was the top enrichment for the comparisons of 25°C versus 40°C under the acetate-depleting condition and 35°C versus 40°C under either acetate condition (temperature effects) (Supplemental Fig. 5B, E, Supplemental Dataset 6).

To further understand our transcriptome data and the effects of acetate, we compared cell physiology at different temperatures between the acetate-depleting and constant-acetate conditions (Fig. 3). Under the acetate-depleting condition without medium dilution, cell density increased over time at 25°C and 35°C but no changes at 40°C because of cell cycle arrest at 40°C (Fig. 3A). Cell density was lower at 35°C than 25°C at 8 h and 24 h time points, suggesting reduced growth rates at 35°C at the end of the acetate-depleting condition. Algal cells treated at 35°C had transiently increased then decreased cell size while 40°C-treated algal cells had steadily increased cell size due to heat-inhibited cell division (Fig. 3B). Under the acetate-depleting condition, chlorophyll content per culture volume (mL) had no significant differences between 25°C and 40°C but was significantly higher at 8-h of 35°C than 25°C (Fig. 3C), consistent with the increased biomass at the 8-h of 35°C (Fig. 1C). Chlorophyll content normalized to cell density revealed constant chlorophyll per cell during 25°C, but transiently and constantly increased cellular chlorophyll during 35°C and 40°C, respectively (Fig. 3D). Chlorophyll content normalized to cell volume showed the increased cellular chlorophyll during 40°C and 35°C could not be completely explained by increased cell volume (Fig. 3E).

**Fig. 3.**
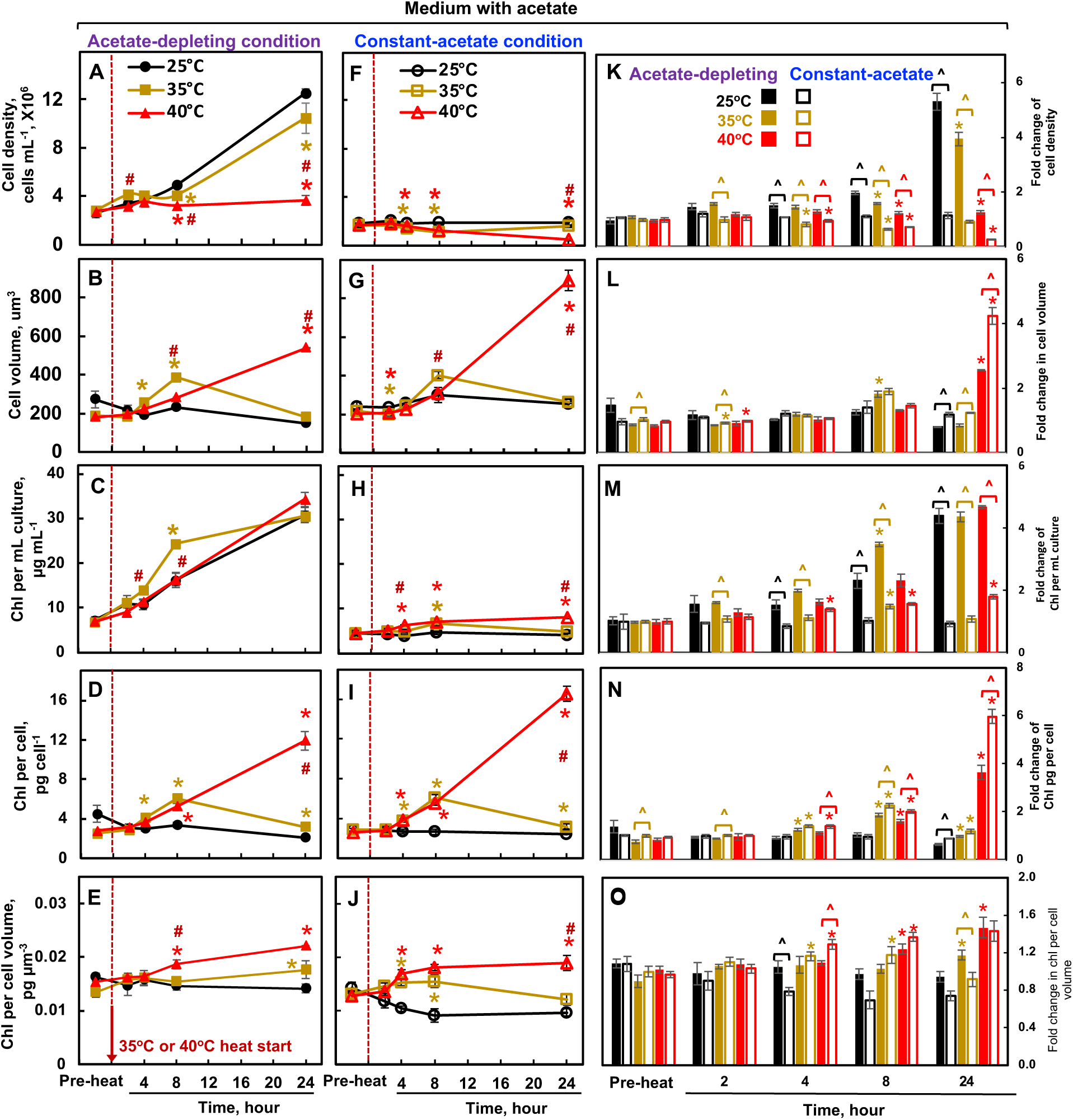
Algal cells subjected to different temperature treatments under the acetate-depleting condition were compared at the physiological level with those under the constant-acetate condition. **(A-E)** Cell parameters from algal cultures under the acetate-depleting condition. The algal cultivation and heat treatments were the same as in Fig. 1B. **(F-J)** Cell parameters from algal cultures under the constant-acetate condition, with turbidostatic control, data plotted based on the results from Zhang *et al*., 2022*a*. (**A-J**) Two panels on the same row share the same y axis. The red dashed lines mark the start of heat or the corresponding time point at constant 25°C. **(K-O)** Fold change of the indicated cell parameters from algal cultures under the acetate-depleting condition (filled bars) and under the constant-acetate condition (empty bars). (**A-O**) Mean ± SE, *n* = 3-7 biological replicates. Statistical analyses were performed with two-tailed t-test assuming unequal variance; *, p<0.05, for the comparisons between 35°C or 40°C with 25°C at the same time point under the same acetate condition, the colors of the asterisks match the heat condition; #, p<0.05, for the comparisons between 35°C and 40°C at the same time point under the same acetate condition; ^, p<0.05, for the comparisons between the acetate-depleting and constant-acetate conditions at the same time point and under the same temperature. **(D, E, I, J)** Cell parameters at different time points during constant 25°C had little change, not significantly different from the pre-heat time points (p>0.05).

Based on our previously published data, we summarized cell parameters in algal cultures grown in the same PBRs but under the 24-h constant-acetate condition through turbidostatic control (Zhang *et al*., 2022*a*) (Fig. 3F-J). Cell parameters at different time points during 25°C had little or no changes as compared to the pre-heat time points, demonstrating the steady algal growth and effectiveness of the turbidostatic control with constant acetate supply. Under the constant acetate condition, the smaller changes of cell density and chlorophyll per mL culture as compared to the acetate-depleting conditions were due to the turbidostatic control and frequent dilution of the culture (Fig. 3F and 3H). The changes of cell volume and chlorophyll content per cell during 35°C and 40°C had similar trends under both the acetate-depleting and constant-acetate condition, but the increase of these parameters was larger under 40°C in cultures with the constant-acetate than the acetate-depleting condition (Fig. 3B, D, G, I). This was supported by the fold changes of cell parameters by comparing the data under the acetate-depleting and constant-acetate conditions (Fig. 3K-O).

In addition to the cellular parameters mentioned above using time-course harvesting, we next utilized non-disruptive and automatic methods to quantify algal growth under different temperatures. The acetate-depleting condition without turbidostatic control allowed us to have a large range of cell densities under different temperature treatments to investigate the optical density (OD) vs cell density relationship and validate automatic culture monitoring under different temperature treatments. OD_680_ (optical density at 680 nm) monitors chlorophyll content per mL culture (Chapman *et al*., 2015; Xiao *et al*., 2015; Young *et al*., 2022). OD_750_ (optical density at 750 nm) monitors light scattering and is thought to be proportional to cell density or cell volume (Chioccioli *et al*., 2014; Young *et al*., 2022). Our PBRs can measure OD_680_ and OD_720_, but not OD_750_. However, OD_720_ serves as a proxy for OD_750_ for light scattering. Thus, we investigated the dynamic changes of OD_680_ and OD_720_ relative to chlorophyll content, cell density, and cell volume before and during heat treatments under the acetate-depleting condition (Fig. 4). The change of OD_680_ and OD_720_ mimicked the change of chlorophyll content per mL algal culture for all three temperature treatments (Fig. 4A, B, 3C). But the change of OD_720_ (Fig. 4B) mimicked the change of cell density for 25°C and 35°C but not 40°C (Fig. 3A). Combining all data from different temperature treatments and time points, both OD_680_ and OD_720_ were linearly proportional to chlorophyll content per mL algal culture (Fig. 4C, D), but they were much less proportional to cell volume or cell density (Fig. 4E-H). Our results showed that OD_680_ and OD_720_ can be used to accurately estimate chlorophyll content in algal cultures with different heat treatments under a wide range of cell densities. Thus, OD_680_ and OD_720_ can be used to estimate algal relative growth rates based on chlorophyll accumulation during an exponential growth phase between two dilution events under the turbidostatic control in PBRs (Zhang *et al*., 2022*a*) (See methods for details). The relative growth rates calculated from both OD_680_ and OD_720_ yielded similar results. Because OD_680_ had larger values and higher signal/noise ratios than OD_720_, we used OD_680_ to estimate relative growth rates of PBR cultures over several days under the turbidostatic controls with and without acetate supply to investigate how acetate affected algal heat responses at 35°C and 40°C in long-term (Fig. 5-8). This research goal could not be achieved without turbidostatic control because our acetate-depleting experiment revealed that acetate was fully depleted in 24 h during all three temperature treatments of 25°C, 35°C and 40°C without constant acetate supply (Fig. 1D).

**Fig. 4.**
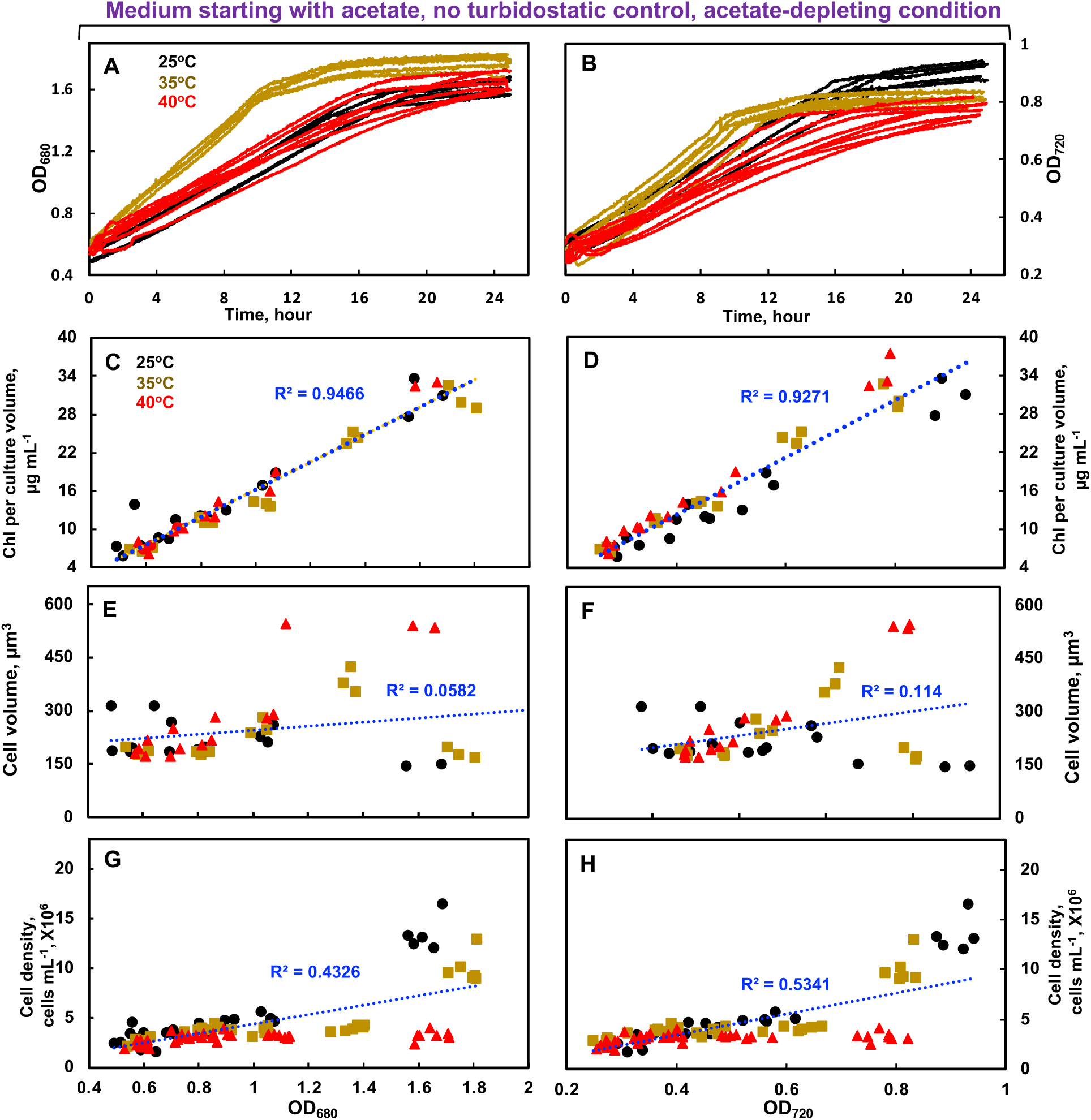
Both OD_680_ and OD_720_ were linearly proportional to chlorophyll contents under different temperatures and can be used to monitor algal growth rates automatically. Black, brown, red lines and symbols represent 25°C, 35°C, and 40°C treatments, respectively. **(A, B)** OD_680_ and OD_720_ automatically monitored algal growth every one min in algal cultures grown in photobioreactors with different temperatures under the acetate-depleting conditions. The algal cultivation and heat treatments were the same as in Fig. 1B. Multiple lines with the same colors represent biological replicates (n= 4-8) under the same condition. **(C-H)** OD_680_ and OD_720_ were linearly proportional to Chlorophyll (Chl) contents per mL cultures but not cell volume or cell density. Chl contents (**C, D**), cell volume (**E, F**), and cell densities (**G, H**) were plotted against OD_680_ or OD_720_ readings in algal cultures with different treatments. Dashed blue lines are linear trendlines of best fit and R-squared values are displayed.

**Fig. 5.**
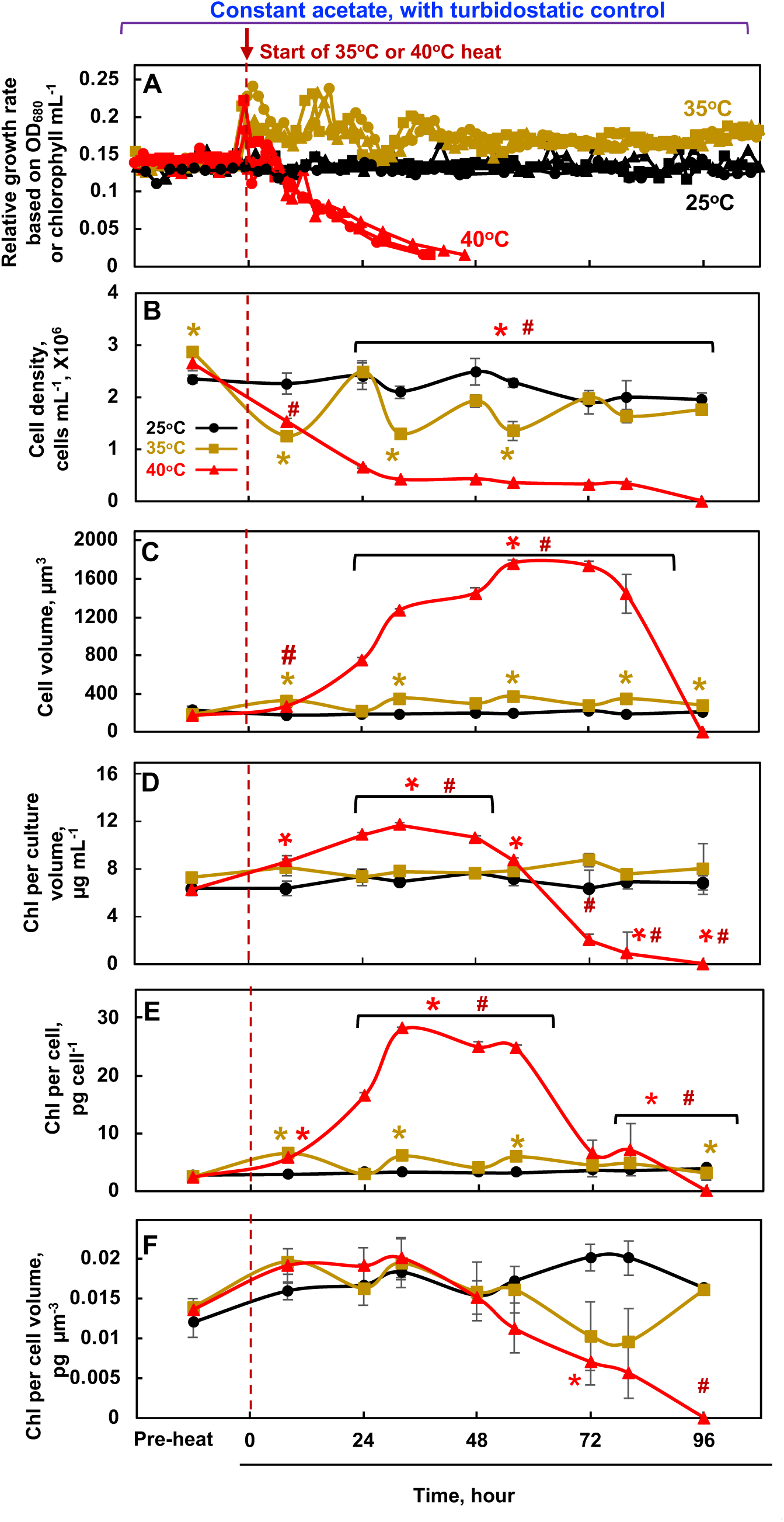
Under the constant-acetate condition with turbidostatic control, moderate (35°C) and acute high temperatures (40°C) increased and reduced algal growth during 4-day heating, respectively. Black, brown, red lines and symbols represent 25°C, 35°C, and 40°C treatments, respectively. The red dashed lines mark the start of heat or the corresponding time point at constant 25°C. **(A)** Chlamydomonas cultures were grown in photobioreactors (PBRs) with turbidostatic control at different temperatures in acetate-containing medium. Algal cultures were first acclimated at 25°C for 4 days before the temperature was switched to 35°C or 40°C or stayed at 25°C for 4 days. Relative growth rates were calculated based on the cycling of OD_680_ caused by the turbidostatic control (see Fig. 1A and method for details). Each temperature treatment had 3 biological replicates in separate PBRs. **(B-F)** Cell parameters were quantified from algal cultures harvested at different time points with different treatments. Statistical analyses were performed with two-tailed t-test assuming unequal variance by comparing 35°C or 40°C with 25°C at the same time point. *, p<0.05, the colors of the asterisks match the heat condition; #, p<0.05, for the comparisons between 35°C and 40°C at the same time point. Chl, Chlorophyll.

**Fig. 6.**
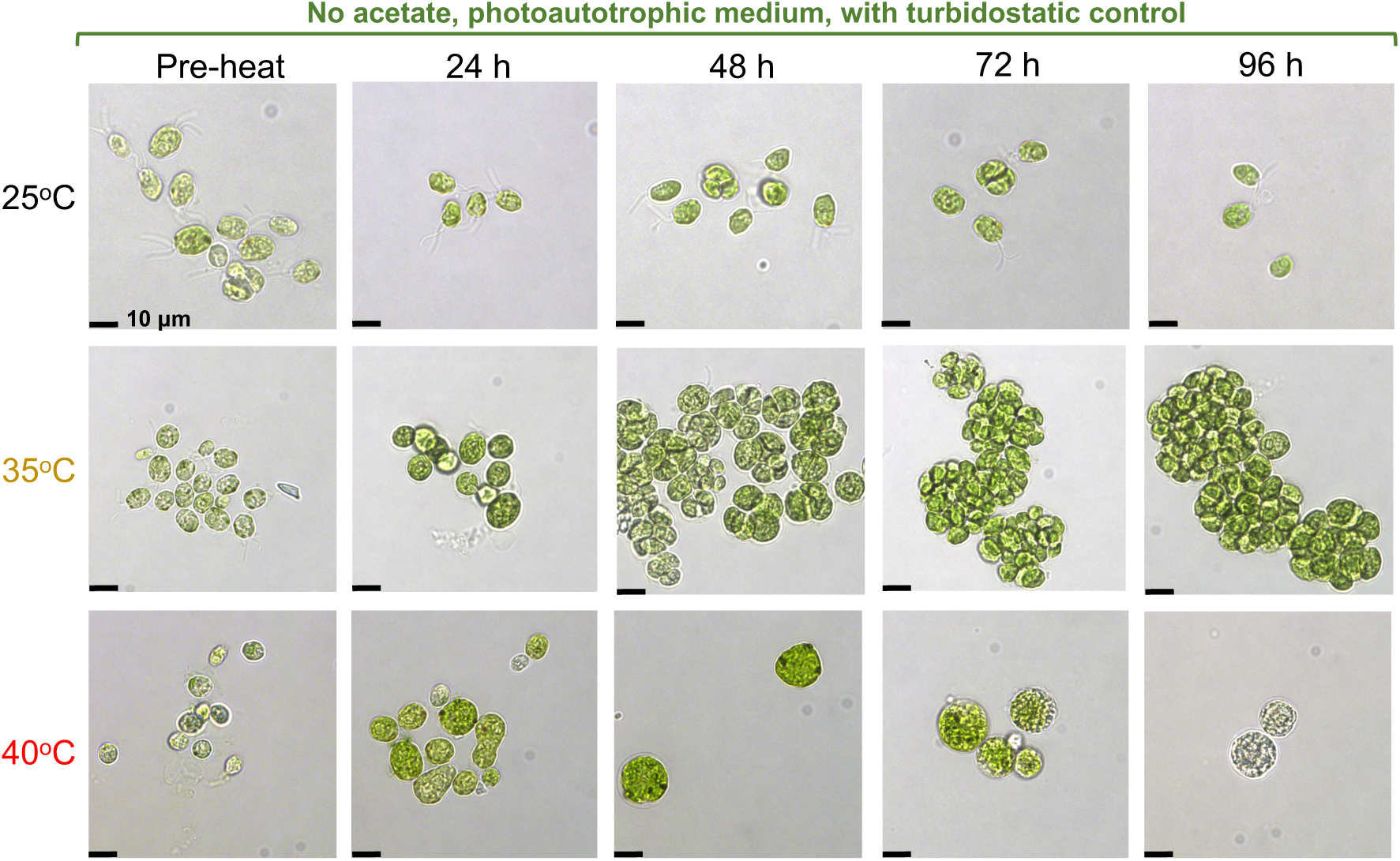
Under the no-acetate condition (photoautotrophic medium) with turbidostatic control, 35°C-treated cells formed palmelloids and 40°C-treated cells were bleached after 4-day heat treatments. Light microscopic images of Chlamydomonas cells under different conditions. Chlamydomonas cultures were grown in photobioreactors with turbidostatic control at different temperatures in photoautotrophic medium without acetate. Algal cultures were first acclimated at 25°C for 4 days before the temperature was switched to 35°C or 40°C or stayed at 25°C for 4 days. Images shown are representative results from three biological replicates. Black scale bar on each panel represents 10 µm.

**Fig. 7.**
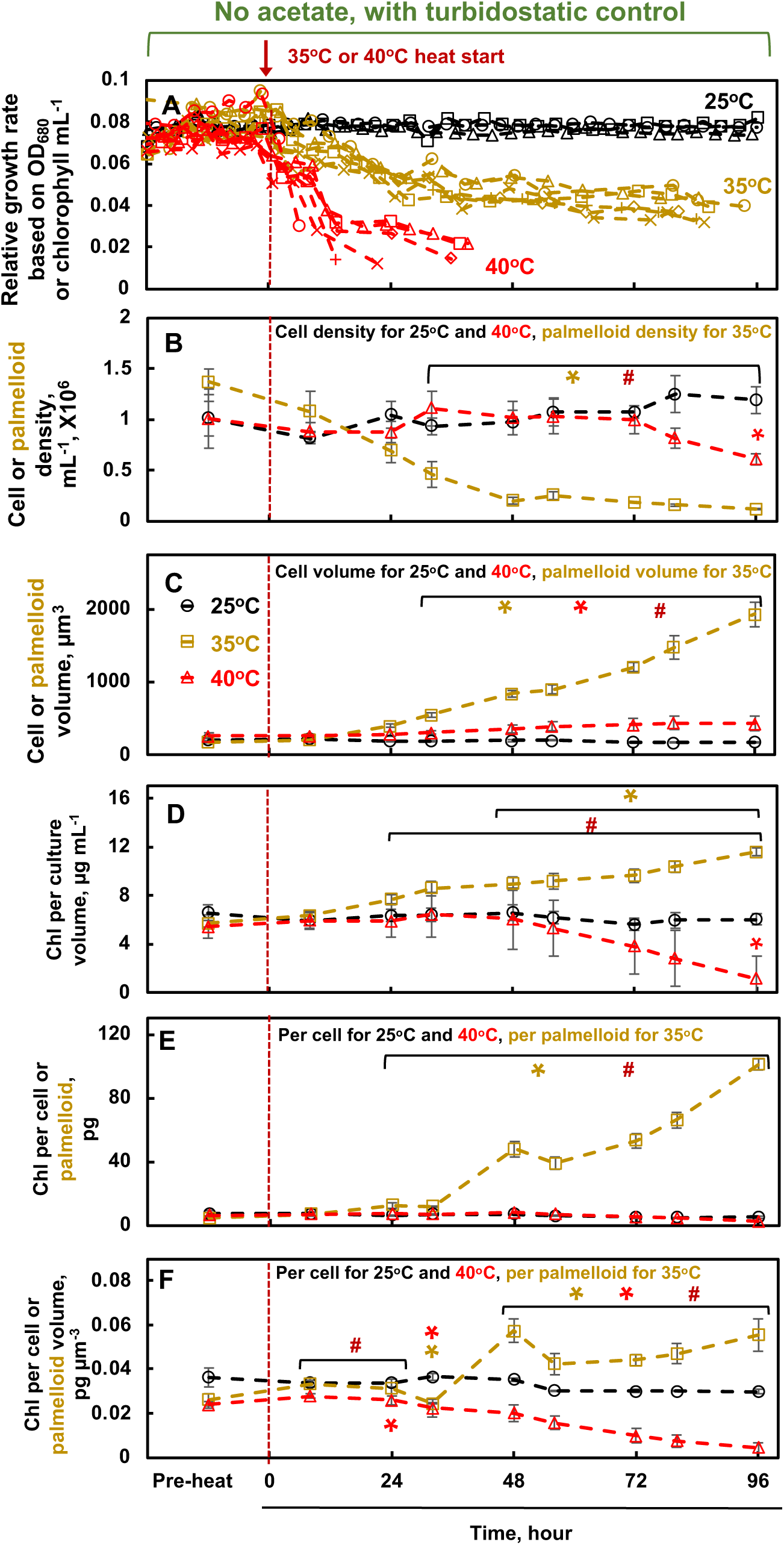
Under the no-acetate condition (photoautotrophic medium) with turbidostatic control, both moderate (35°C) and acute high temperatures (40°C) reduced algal growth during 4-day heating. Black, brown, red lines and symbols represent 25°C, 35°C, and 40°C treatments, respectively. The red dashed lines mark the start of heat or the corresponding time point at constant 25°C. **(A)** Chlamydomonas cultures were grown in photobioreactors (PBRs) with turbidostatic control at different temperatures in photoautotrophic medium without acetate. Algal cultures were first acclimated at 25°C for 4 days before the temperature was switched to 35°C or 40°C or stayed at 25°C for 4 days. Relative growth rates were calculated based on the cycling of OD_680_ caused by the turbidostatic control. Each temperature treatment had at least 3 biological replicates in separate PBRs. **(B-F)** Cell parameters were quantified from algal cultures harvested at different time points with different treatments. Because 35°C-treated cells under photoautotrophic condition formed palmelloids (see Fig 6), the term of “palmelloid” was used in place of “cell” for this condition only. Statistical analyses were performed with two-tailed t-test assuming unequal variance by comparing 35°C or 40°C with 25°C at the same time point. *, p<0.05, the colors of the asterisks match the heat condition; #, p<0.05, for the comparisons between 35°C and 40°C at the same time point. Chl, Chlorophyll.

**Fig. 8.**
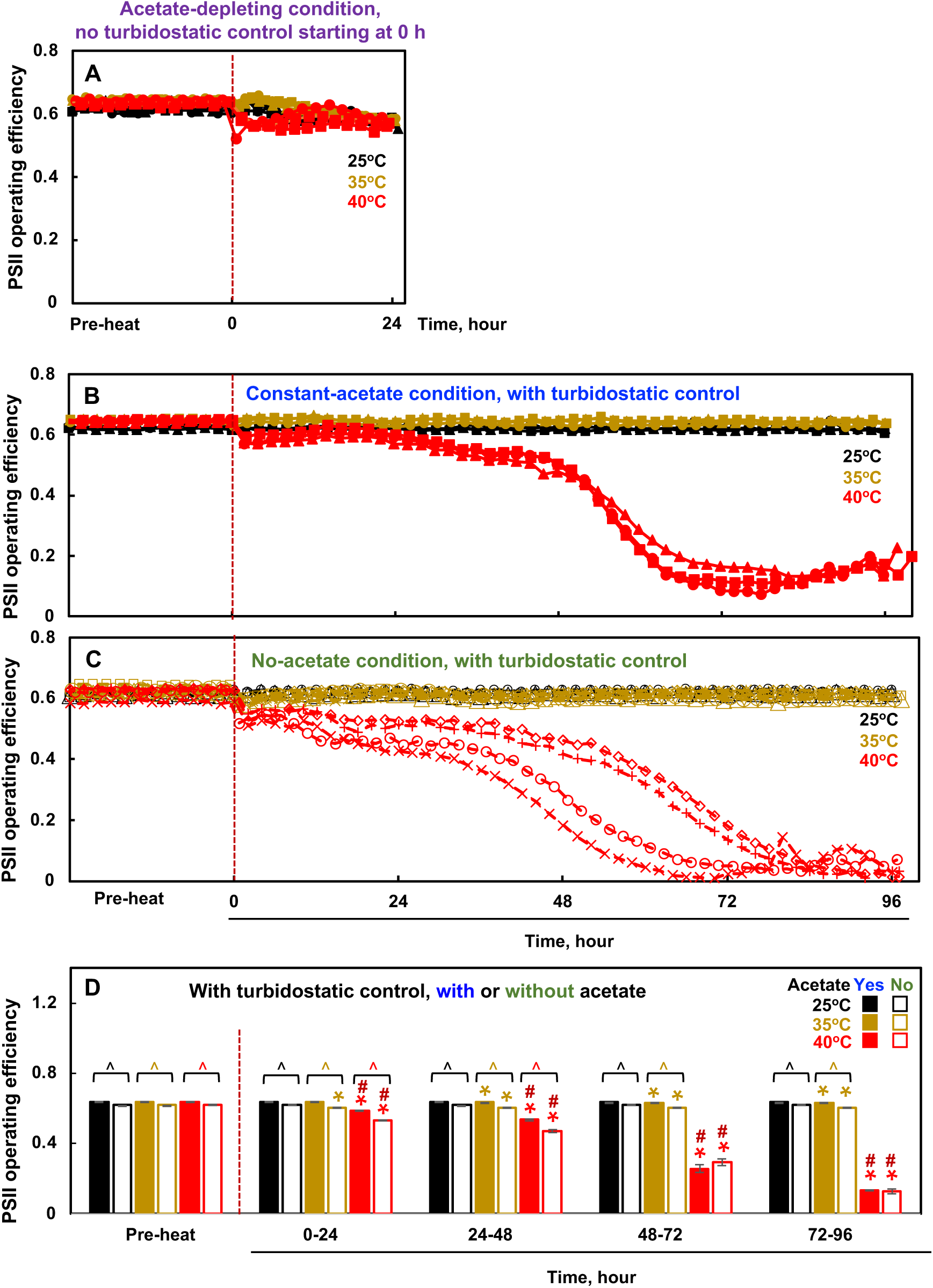
Under the turbidostatic control and no nutrient limitation, algal cultures had higher photosystem II (PSII) operating efficiency with acetate than without acetate during treatment of 25°C, 35°C, or 40°C, although the acetate effects diminished after 2-day heat of 40°C due to cell death. PSII operating efficiency was measured automatically in photobioreactors under different conditions. Black, brown, red lines and symbols represent 25°C, 35°C, and 40°C treatments, respectively. The red dashed lines mark the start of heat or the corresponding time point at constant 25°C. **(A)** Acetate-depleting condition at 25°C, 35°C, or 40°C for 24 h, in medium started with acetate but no turbidostatic control, thus no constant acetate supply. **(B)** Constant-acetate condition at 25°C, 35°C, or 40°C for 4 days, in medium with acetate and turbidostatic control, thus no nutrient limitation. **(C)** No-acetate condition at 25°C, 35°C, or 40°C for 4 days, in photoautotrophic medium without acetate but with turbidostatic control, thus no nutrient limitation. **(D)** The comparison of PSII operating efficiency in the constant-acetate (data from panel B, filled bars) and no-acetate (data from panel C, empty bars) conditions during the indicated periods at 25°C, 35°C, or 40°C for 4 days. Statistical analyses were performed with two-tailed t-test assuming unequal variance by comparing the indicated groups. *, p<0.05, comparing 35°C or 40°C with 25°C with the same medium type at the same time; #, p<0.05, for the comparisons between 35°C and 40°C with the same medium type at the same time; ^, p<0.05, for the comparisons between the same temperature treatment at the same time but with and without acetate. The colors of the significant marks match the heat condition, brown for 35°C and bright red color for 40°C related comparisons, and dark red for 35°C and 40°C comparisons.

To investigate the effects of acetate on long-term heating, after algal cultures reached steady growth at 25°C in PBRs with turbidostatic control, we conducted all three temperature treatments with continuous turbidostatic control and constant acetate supply for 4 days (Fig. 5). Without heat treatments, algal growth rates and cell parameters stayed constant at 25°C, demonstrating the effectiveness of our turbidostatic control for steady algal growth (Fig. 5A). At 35°C with constant acetate supply, the relative growth rates increased first, but the increase was smaller after 2-day heating at 35°C and stabilized thereafter. At 40°C, the relative growth rates decreased steadily, and the cultures died after 2-day heating at 40°C, suggesting algal cells could not acclimate to long-term constant heat at 40°C, even with constant acetate supply. Cells treated with 40°C had more than 4-fold increase of cell volume as compared to the pre-heat condition, followed by reduced cellular chlorophyll and cell death (Fig. 5C-F). The turbidostatic mode by OD_680_ tightly controlled the chlorophyll contents in unit of ug per mL culture in PBRs during the treatments of 25°C and 35°C (Fig. 5D), but not 40°C due to the cell cycle arrest and overwhelmingly increased chlorophyll per cell, and eventually cell death under 40°C (Fig. 5B and E).

To further test the effects of acetate on algal heat responses, we conducted experiments using algal cultures grown in PBRs under the turbidostatic control with constant nutrient supply of photoautotrophic medium (no acetate) and different temperature treatments for 4 days (Fig. 6, 7). Our microscopic images showed that 35°C-treated cells in photoautotrophic medium formed palmelloids (Fig. 6). Palmelloids are clumps of non-motile cells, which often result from a failure of hatching or release from the mother cell wall under stressful conditions (Harris, 2009). It was reported that Chlamydomonas cells aggregated to form palmelloids after 3 min acute heat shock at 50°C in acetate-containing medium (de Carpentier *et al*., 2022). The formation of palmelloids did not occur in our constant-acetate-35°C condition (Zhang *et al*., 2022*a*), suggesting 35°C was more stressful without acetate than with acetate. Without heat treatments, algal growth rates and cell parameters stayed constant at 25°C, demonstrating the effectiveness of our turbidostatic control for steady algal growth under photoautotrophic conditions (Fig. 7A). Without acetate but with constant medium supply, algal growth had gradually decreased growth rates during 35°C (Fig. 7A), unlike the increased growth rates during 35°C with constant acetate supply (Fig. 5A), strongly supporting the role of acetate in algal heat tolerance. Without acetate, algal cells with 35°C treatment had increased palmelloid volume (or size) and chlorophyll contents (Fig. 7B-F). The reduction of growth rates during 40°C was comparable with and without acetate (Fig. 5A and 7A).

Our PBRs could also monitor photosystem II (PSII) operating efficiency using chlorophyll fluorescence measurement automatically (Fig. 8). Heat at 40°C reduces PSII operating efficiency dramatically, with more reduction without acetate than with acetate during the first two days of heating (Fig. 8B, C, D). The acetate effects on PSII operating efficiency disappeared in algal cells treated with 40°C for more than 2 days (Fig. 8D). The reduction of PSII operating efficiency during 35°C was much smaller than during 40°C, with higher PSII operating efficiency in 35°C-treated algal cells with acetate than without acetate (Fig. 8D).

## Discussion

By performing algal cultivation under highly controlled conditions in PBRs with and without the constant organic carbon source, acetate, we investigated how acetate availability affected the growth of Chlamydomonas under moderate (35°C) and acute high temperatures (40°C) (Fig. 9). The moderate high temperature of 35°C is frequently experienced by outdoor, open algal ponds in summertime and enclosed outdoor algal ponds can often heat up to 40°C or above (Krishnan *et al*., 2021). Thus, both high temperatures we chose were physiologically relevant for algae in nature. Additionally, global warming increases the duration of moderate and acute high temperatures in the field conditions. By using highly controlled turbidostatic conditions in PBRs, our research not only investigated the short-term (within 24 h) but also long-term (up to 4-days) heat effects on algal growth with and without acetate supply or under the acetate-depleting conditions. Our research advanced the understanding of algal heat responses under different heat intensities, heat durations, and carbon availability.

**Fig. 9.**
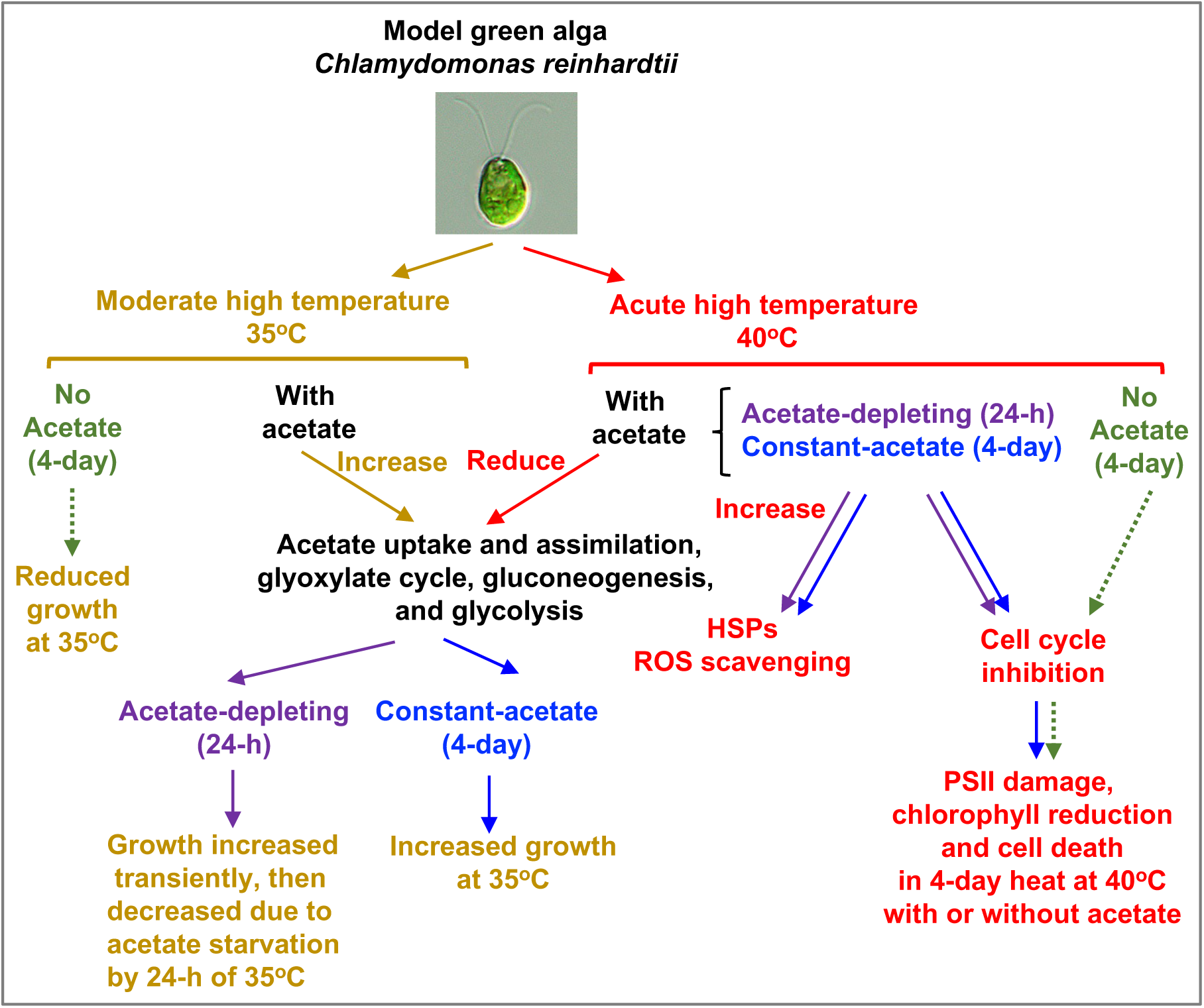
Acetate availability affected Chlamydomonas heat tolerance under the moderate high temperature (35°C) but had much less effects under the acute high temperature (40°C). In medium containing the organic carbon source, acetate, heat of 35°C accelerated acetate metabolism by upregulating transcripts related to acetate uptake and assimilation, glyoxylate cycle, gluconeogenesis, and glycolysis pathways. With constant acetate supply, 35°C increased algal growth. Under the acetate-depleting condition, 35°C increased algal growth transiently followed by decreased growth and biomass accumulation due to acetate starvation. Without acetate supply in photoautotrophic medium, 35°C reduced algal growth rates. Under either constant-acetate or depleting-acetate condition, heat of 40°C down-regulated transcripts related to acetate metabolism, but up-regulated transcripts related to heat shock proteins (HSPs) and reactive oxygen species (ROS) scavenging. In medium with or without acetate, 40°C decreased algal growth rates, inhibited cell division, damaged photosynthesis (e.g., photosystem II, PSII), reduced chlorophyll contents, and eventually led to cell death. Thus, Chlamydomonas cells could not survive continuous heat of 40°C for longer than 2 days, independent of acetate supply. Brown and red labels and lines represent the effects of 35°C and 40°C, respectively. Purple, blue, and green labels and lines represent the acetate-depleting, constant-acetate, and no-acetate conditions, respectively.

### Heat of 35°C was beneficial or detrimental depending on carbon availability

Under the turbidostatic control with frequent medium dilutions, moderate high temperature of 35°C increased algal growth rates with constant acetate supply (Fig. 5A) but decreased algal growth rates without acetate in photoautotrophic medium (Fig. 7A), confirming the important role of acetate in heat tolerance. The increased heat tolerance at 35°C with acetate is not strain-dependent because the acetate effects were similar in two different algal strains: CC-1690 (this study) and CC-5325 (Mattoon *et al*., 2022*a*). In medium starting with acetate but without constant acetate supply (the acetate-depleting condition), algal biomass increased first but then decreased at 35°C compared to 25°C, which can be explained by increased acetate uptake and usage initially followed by acetate depletion and starvation by the end of the 24-h 35°C treatments (Fig. 1C, D).

Under the acetate-depleting condition without medium dilution, besides acetate, other nutrients may also be depleted gradually during 24 h. We checked known transcription factors and regulators responding to starvation of other nutrients, including *nitrogen starvation responsive regulator 1 (NRR1)* (Boyle *et al*., 2012), *sulfur starvation responsive regulator 2.1 (SNRK2.1)* (González-Ballester *et al*., 2010), *phosphorus starvation response 1 (PSR1)* (Bajhaiya *et al*., 2016), *copper deficiency response regulator 1 (CRR1)* (Sommer *et al*., 2010), and *iron deficiency response regulator (Mitogen-Activated Protein Kinase 3, MAPK3)* (Fei *et al*., 2017). If these nutrients were depleted, the corresponding transcription factors and regulators would be up-regulated. Most of these nutrient-deficiency transcription factors and regulators had minimal or no induction at 24 h of the acetate-depleting condition (Supplemental Fig. 3C-G). One exception is the *NRR1*, which was up-regulated at 25°C or 35°C but was down-regulated under 40°C at 24 h of the acetate-depleting condition (Supplemental Fig. 3C). The results suggested minimal or no limitation of sulfur, phosphorus, copper, and iron during the acetate-depleting condition, but possibly some nitrogen deficiency at 24 h of the acetate-depleting condition at 25°C and 35°C but not 40°C. One of the responses to nitrogen deficiency is the reduction of chlorophyll (Schmollinger *et al*., 2014). During our acetate-depleting condition, most transcripts involved in chlorophyll biosynthesis were stable (Supplemental Fig. 4Q). Additionally, there was no change of the chlorophyll per cell or per cell volume at 24 h of 25°C as compared to the pre-heat time point which did not have acetate-limitation; these parameters were higher under 35°C and 40°C than 25°C at 24 h of the acetate-depleting condition (Fig. 3D, E). Collectively, the results indicated that the nitrogen limitation may have just started at 24 h of the acetate-depleting condition, thus would not majorly impact cell physiology. Finally, our results from the 4-day constant-acetate and no-acetate conditions under the turbidostatic control strongly pointed to the specific effects of acetate on algal heat tolerance but not other nutrients (Fig. 5, 7). Thus, the acetate effects are the focus of this paper.

Chlamydomonas uptakes acetate and feeds it into the glyoxylate cycle and gluconeogenesis for starch biosynthesis and starch is broken down through glycolysis to make cellular energy (Johnson and Alric, 2012, 2013) (Fig. 2A). The dynamic effects of 35°C on algal growth under different acetate conditions can be explained by the 35°C-induced acetate metabolism, supported by increased acetate consumption and up-regulated transcripts related to acetate metabolism pathway at 35°C (Fig. 1, 2). Energy produced from glycolysis can be used for energy-requiring cellular activities to increase thermotolerance (Olas *et al*., 2021), e.g., repair pathways related to photosynthesis (Murata and Nishiyama, 2018). Pyruvate kinase catalyzes the final step of glycolysis by converting phosphoenolpyruvate and one ADP to pyruvate and one ATP (Baud *et al*., 2007; Wulfert *et al*., 2020). Chlamydomonas mutants deficient in pyruvate kinase were heat-sensitive under 35°C in acetate-containing medium (Mattoon *et al*., 2022*a*), supporting the important roles of glycolysis in thermotolerance of 35°C. Although ATP is primarily produced through mitochondrial oxidative respiration, the ATP production from glycolysis can be important under stressful conditions where energy availability is limited (van Dongen *et al*., 2011). Under low oxygen conditions, plants increase activity of pyruvate kinases to produce more ATP (van Dongen *et al*., 2011). Heat-treated barley leaves utilized glycolysis as an alternative energy source for thermotolerance based on proteomics analysis (Rollins *et al*., 2013).

Additionally, acetate may protect photosynthesis from heat-induced photoinhibition. Acetate is proposed to protect PSII against photoinhibition by replacing the bicarbonate associated to the non-heme iron at the acceptor side of PSII, changing the environment of plastoquinone and affecting PSII charge recombination (Roach *et al*., 2013). Our PSII efficiency data supported this report because PSII efficiency was higher with acetate than without acetate during 25°C, 35°C or 40°C treatments (Fig. 8D). Furthermore, experimental and modeling analysis suggested that acetate promoted cyclic electron flow (CEF) around photosystem I (PSI); reducing equivalents produced during the acetate metabolism reduce plastoquinone pools and increase CEF activity (Johnson and Alric, 2012; Lucker and Kramer, 2013; Chapman *et al*., 2015). CEF cycles photosynthetic electrons around PSI, producing only ATP but no NADPH and providing extra ATP needed for photosynthesis and other cellular activities (Munekage *et al*., 2004; Baker *et al*., 2007). CEF balances the ATP/NADPH ratio, contributes to the generation of transthylakoid proton motive force, and protects both PSI and PSII from photo-oxidative damage (Johnson, 2011; Yamori and Shikanai, 2016). The increased acetate uptake under 35°C (Fig. 1D) was coupled with induced CEF activity in Chlamydomonas under the constant-acetate condition (Zhang *et al*., 2022*a*). Our transcriptome analysis also showed up-regulation of transcripts related to PSI and CEF at 2 h of 35°C treatments under either the constant-acetate or acetate-depleting condition (little acetate depletion at 2 h), consistent with heat-induced CEF in the presence of acetate (Supplemental Fig. 4J, P).

The effects of acetate on the improved heat tolerance under the moderate high temperature may also be related to the accumulation of thermoprotective metabolites (osmolytes, including amino acids, sugars, and sugar alcohols) (Guihur *et al*., 2022) or compatible solutes (Hemme *et al*., 2014; Schroda *et al*., 2015). Compatible solutes accumulate from the catabolism of large molecules upon the onset of heat to stabilize membranes and prevent protein unfolding and aggregation (Schroda *et al*., 2015). In Chlamydomonas, these compatible solutes include glycerophos-phoglycerol (GPG), trehalose, alanine, isoleucine, asparagine, glutamine, and guanosine (Hemme *et al*., 2014). The elevated acetate metabolism at 35°C with constant acetate supply may increase the accumulation of these thermoprotective compatible solutes. This is an understudied but exciting area. Future metabolite analysis may provide more information about this.

With constant acetate supply, 35°C is beneficial, increasing carbon metabolism and improving thermotolerance for enhanced growth (Fig. 5). If algal cells can be cultivated under sterile conditions with constant organic carbon supply, 35°C could be used to promote algal growth and increase biofuel and bioproduct generation under mixotrophic conditions in light. Culture temperature of 35°C in closed, outdoor algal ponds may not be difficult to reach with natural sunlight heating in the summertime of the moderate climate regions and year-round in the tropical regions. If photoautotrophic medium is used for enclosed outdoor algal cultivation, acetate could be added to improve algal thermotolerance before the occurrence of moderate high temperature around 35°C. The improved algal thermotolerance and growth may outweigh the cost of added acetate.

Without constant acetate supply, 35°C is detrimental. Under the acetate-depleting condition, the 35°C-elevated carbon metabolism without sufficient carbon input may deplete cellular carbon reserves. This was not sustainable in the long-term and eventually reduced biomass accumulation (Fig. 1C). With constant acetate supply, heat at 35°C transiently arrested the cell cycle, which recovered after 8 h at 35°C (Fig. 3G). In photoautotrophic medium without acetate, 35°C reduced algal growth rates and resulted in stressed cells aggregated to form palmelloids (Fig. 6, 7). Such effects of 35°C can particularly compromise the yields of outdoor algal ponds because they usually do not contain carbon sources but frequently experience moderate high temperatures (Mata *et al*., 2010; El-Sheekh *et al*., 2019).

It is reported that the growth rate of another Chlamydomonas WT strain (CC-125) at 35°C was about 50% of that at 28°C (0.083 vs 0.166) in photoautotrophic medium under constant light and continuous dilution but with 2% CO_2_ (Vítová *et al*., 2011). The 50% growth reduction is comparable to the reduced growth of our data at 35°C without acetate (Fig. 7A). We used air level CO_2_, which is 400 ppm or 0.04% CO_2_ to mimic the nature condition (1/50^th^ of 2% CO_2_ as in Vítová *et al*., 2011). CO_2_ provides the inorganic carbon source for algal growth. However, the equivalent 50% growth reduction at 35°C as compared to the control temperature with (Vítová *et al*., 2011) and without (this study) high CO_2_ suggests CO_2_ (inorganic carbon source) is not as effective as acetate (organic carbon source) to improve thermotolerance in algal cells. Furthermore, different algal strains with changes in cultivation conditions may have varying moderate high temperature ranges. For example, another Chlamydomonas WT strain (CC-137) had optimal growth at 35°C in photoautotrophic medium with 2% CO_2_ in day/night cycle with semi-dilution (one dilution per day at the end of dark phase) (Lien and Knutsen, 1979). Such semi-dilution may result in a large range of cell densities, which may affect the light perceived by each cell. Additionally, the day/night cycle may also alleviate photooxidative stress as photosynthesis only occurs in light. However, the author mentioned that cytokinesis and sporulation became inhibited at temperature above 36°C (Lien and Knutsen, 1979), indicating 35°C may be close to the moderate high temperature range that would reduce algal growth in this strain without acetate under their cultivation condition. Additionally, the fast-growing marine alga *Picochlorum celery*, which is tolerant to high salt, high light, and high temperature, had maximal growth rates around 35°C which was reduced at temperatures around 40°C, with photoautotrophic medium and unlimited CO_2_ supply in outdoor open ponds under semi-continuous, drain and fill mode of operation (cells were harvested and fresh medium was added every 2-3 days) (Krishnan *et al*., 2021), suggesting the moderate high temperature range is strain-specific and depending on the cultivation condition.

Furthermore, some tissues in land plants are either mixotrophic or heterotrophic, such as reproductive tissues (e.g., developing green seeds) (Koley *et al*., 2022) or sink tissues (e.g., roots). Reproductive tissues are often considered the most heat sensitive in land plants (Barnabás *et al*., 2008; Ullah *et al*., 2022) and are the main harvest of grain crops such as soybean, wheat, barley, sorghum, and maize. The up-regulated carbon metabolism under moderate high temperatures may also occur in land plants, especially in reproductive or sink tissues. Heat-induced glyoxylate cycle, gluconeogenesis, and glycolysis have been reported in plants (Rollins *et al*., 2013; Zhang *et al*., 2013; Aprile *et al*., 2013). Carbon metabolism and sugar availability were shown to be essential for heat tolerance in Arabidopsis (Olas *et al*., 2021). The thermotolerant wheat cultivar had increased carbohydrate molecules in sink tissues (Mishra *et al*., 2021). It is also reported that heat stress impaired sucrose metabolism in leaves and anthers, which were associated with heat-induced reproductive failures in chickpea (Kaushal *et al*., 2013). A better understanding of the interaction between carbon availability and heat tolerance could be leveraged to protect sink and/or reproductive tissues from heat stresses.

### Heat of 40°C was detrimental to algal cells with and without constant acetate supply

Acute high temperatures at or above 40°C inhibit the algal cell cycle (Fig. 3, 5-7) (Mühlhaus *et al*., 2011*b*; Hemme *et al*., 2014; Zachleder *et al*., 2019; Ivanov *et al*., 2021), alter thylakoid membranes, reduce photosynthesis, and damage respiration (Zhang *et al*., 2022*a*). These cellular damages took place quickly, mostly within 4 h of heat at 40°C with constant acetate supply (Zhang *et al*., 2022*a*). Although 40°C-treated cells grew bigger and had more chlorophyll per cell under the constant-acetate condition than the acetate-depleting condition in the short-term (24-h heat) (Fig. 3B, G, D, I, L, N), the constant acetate supply could not prevent ultimate cell death with 40°C-heat longer than 2 days (Fig. 5). Without acetate, Chlamydomonas cells also died after 2-day heat at 40°C (Fig 7). The death of Chlamydomonas cultures was also reported at 39°C for 33 h in photoautotrophic medium without acetate using the same WT strain *21gr* (Zachleder *et al*., 2019; Ivanov *et al*., 2021). In conclusion, constant heating at 40°C is lethal for Chlamydomonas cells independent of acetate availability.

At 24-h of the acetate-depleting condition, algal cells exposed to 40°C had very little division (only 30% increase of cell density, thus 1.3 X) but they still grew and accumulated biomass by increasing cell volume (3 X) as compared to the pre-heat condition (Fig. 3B). In comparison, algal cells exposed to 25°C at 24 h of the acetate-depleting condition had 5.5 X increased cell density but reduced cell volume (55% of the pre-heat values, due to acetate starvation). If we use fold change of cell density multiplied by fold change of cell volume to estimate biomass, it could explain the comparable amount of dry biomass in cultures treated with 40°C and 25°C by the end of 24-h treatment under the acetate-depleting condition (Fig. 1C). However, the inefficient system with reduced carbon input and little cell division at 40°C was not sustainable, eventually killing all the algal cells by the end of 2-day heat of 40°C, even with constant acetate supply (Fig. 5). Our results suggested that the increased thermotolerance by acetate did not apply to long-term heat at 40°C.

Acetate had very limited effects on the tolerance to 40°C in Chlamydomonas possibly because 40°C damaged mitochondrial activities, which are important for acetate metabolism. Most of the reducing power from acetate assimilation is used in mitochondrial respiration to produce ATP (Johnson and Alric, 2012, 2013). Our previous results showed that 40°C reduced mitochondrial activities significantly under the 24-h constant-acetate condition (Zhang *et al*., 2022*a*). When comparing 40°C vs 25°C under the acetate-depleting condition and 40°C vs 35°C independent of the acetate condition, the top enriched function group was related to mitochondrial electron transport (Supplemental Fig. 5B, E), suggesting mitochondria were particularly sensitive to 40°C. This was also supported by the down-regulation of transcripts related to mitochondrial electron transport, especially those related to F1-ATPase at 2 h of 40°C (Supplemental Fig. 4AA, AB, Supplemental Dataset 2). The compromised mitochondrial activities may restrict acetate uptake and assimilation during 40°C heat treatment. Although there were no significant differences in acetate consumption between 40°C and 25°C under the 24-h acetate-depleting conditions (Fig. 1D), the differences may be underestimated and masked by the acetate depletion. Transcripts of many genes involved in carbon metabolism, e.g., acetate uptake and assimilation, glyoxylate cycle and gluconeogenesis pathways, and glycolysis were significantly down-regulated during 40°C under either the acetate-depleting or constant-acetate condition, especially at 2 h heat (Fig. 2), consistent with the down-regulated transcripts related to mitochondrial electron transport. Heat at 40°C for longer than 1 day would eventually slow down mitochondrial activities and reduce acetate-uptake. The decreased acetate uptake and assimilation, in addition to reduced photosynthesis and arrested cell cycle, may result in the ultimate death of the algal culture with 2-day heat at 40°C (Fig. 3, 5-8).

Under the acetate-depleting condition, without frequent medium dilution, cell densities increased at 25°C and 35°C during the 24-h experiments (Fig. 3A). With the constant light source, the increased cell densities resulted in light shading. However, the amount of chlorophyll per mL culture was comparable at 24 h of 25°C, 35°C, and 40°C under the acetate-depleting condition, suggesting comparable level of light shading under all three temperatures (Fig. 3C). Thus, we could cross off the effects of light shading when comparing the results at the 24 h of 25°C, 35°C, and 40°C under the acetate-depleting condition. Additionally, the up-regulated transcripts related to photoprotection at 24 h of all three temperatures under the acetate-depleting condition when light shading was present indicated compromised or inefficient photosynthesis that was not due to high light stress (Supplemental Fig. 4O). The up-regulation of transcripts related to photoprotection at 24 h of 25°C and 35°C was only present under the acetate-depleting condition but not the constant-acetate condition, which may be due to reduced CO_2_ concentrations with increased cell density without frequent culture dilution, supported by a recent report that low CO_2_ induced photoprotection independent of light (Ruiz-Sola *et al*., 2023). The up-regulation of transcripts related to photoprotection at 24 h of 40°C was similar under either acetate condition, just with higher induction under the acetate-depleting than the constant-acetate condition, suggesting it was mainly caused by damaged photosynthesis at 40°C.

At 24-h of the acetate-depleting condition, the transcripts related to large *HSP*s, *HSP60/70/90s*, were down-regulated at 25°C and 35°C, but not under 40°C (Supplemental Fig. 4A-C). The down-regulation of large *HSP*s may be related to the increased cell density and light shading, thus reduced light stress at 24 h of 25°C and 35°C under the acetate-depleting condition (Fig. 3A). Besides heat stress, transcripts of large *HSPs* also responded to light and were also up-regulated under high light stresses (Anderson et al. 2021). Additionally, the down regulation of large *HSP*s at 25°C and 35°C but not 40°C at 24 h of the acetate-depleting condition may suggest the energy distribution between growth and heat tolerance. Most large HSPs are ATP-dependent chaperones, especially HSP90s (Al-Whaibi, 2011; Xu *et al*., 2012; Guihur *et al*., 2022). Down-regulation of these large HSPs when acetate was depleted may be helpful to reserve energy for cell growth when the temperature stress was none (25°C) or moderate (35°C). However, under the acute heat of 40°C, thermotolerance has higher priority than growth, thus the expression of these large *HSPs* were maintained while cell cycle was arrested to preserve energy (Supplemental Fig. 4A-C,W, X).

As compared to 40°C, the induction of *HSP*s at 35°C was minimal or none under either acetate condition (Supplemental Fig. 4A-C), suggesting the 35°C-induced thermotolerance by acetate may not be related to the accumulation of HSPs. This may suggest some none-overlapping heat sensing and signaling pathways between 35°C and 40°C, because HSPs have important roles in the classical heat shock responses (Guihur *et al*., 2022). Several heat sensors have been proposed (Vu *et al*., 2019), including membrane fluidity (Martinière *et al*., 2011), Ca^2+^ channels in the plasma membrane (Gao *et al*., 2012; Finka *et al*., 2012), unfolded proteins (Rütgers *et al*., 2017; Li *et al*., 2020), and phytochromes (Jung *et al*., 2016; Legris *et al*., 2016), and some circadian clock components (e.g. EARLY FLOWERING 3, ELF3) (Jung *et al*., 2020). Among these, the heat effects on phytochromes and ELF3 have been reported around the temperature ranges of 30°C-35°C (Jung *et al*., 2016, 2020; Legris *et al*., 2016), providing potential heat sensor candidates for moderate high temperatures. Future research to investigate the shared and unique heat sensors under moderate and acuate high temperatures may provide more information about how plants sense and adapt to different high temperatures.

High temperatures affect Chlamydomonas lipid metabolism (Schroda *et al*., 2015). To overcome heat-induced membrane fluidity and leakiness, Chlamydomonas reduced unsaturated membrane lipids and increased the production of saturated lipids to restore normal membrane viscosity (Schroda *et al*., 2015; Légeret *et al*., 2016). Saturated fatty acids are more stable than unsaturated fatty acids under high temperatures. It was proposed that the diacylglycerols (DAGs) acyltransferase, DGTT1, converted the chloroplast unsaturated membrane lipids monogalactosyldiacylglycerols (MGDG) to DAGs then triacylglycerols (TAGs) in Chlamydomonas with 1 h heat at 42°C (Légeret *et al*., 2016). Our RNA-seq data showed that the transcripts of *DGTT1* were quickly and largely up-regulated at 40°C independent of acetate availability (Supplemental Figure 3H), consistent with a strategy to reduce unsaturated membrane lipids during high temperatures. The transcripts of *DGTT1* were also up-regulated at 35°C, but with a smaller induction than at 40°C. Additionally, a long chain fatty acid CoA ligase, LCS2, involves in de novo synthesis of saturated fatty acids by incorporating newly synthesized fatty acids exported from the plastid into saturated TAGs in lipid droplets (Li *et al*., 2016; Li-Beisson *et al*., 2023). Our RNA-seq data showed that the transcripts of *LCS2* were quickly and significantly up-regulated at 40°C independent of acetate availability (Supplemental Figure 3I), suggesting increased de novo synthesis of saturated fatty acids at 40°C. The de novo fatty acids biosynthesis is the only option for increasing the fraction of lipids with saturated fatty acids (Hemme *et al*., 2014). Heat at 40°C reduced photosynthetic activities within 4 h, which increased chloroplast redox pressure and oxidative stress under high temperatures (Zhang *et al*., 2022*a*). Biosynthesis of lipids with saturated fatty acids via the de novo pathway requires reducing power NADPH and energy ATP (Ohlrogge and Browse, 1995), which may serve as an electron sink and alleviate heat-induced redox stresses (Hemme *et al*., 2014). The transcripts of *LCS2* were also up-regulated at 24 h of 25°C and 35°C under the acetate-depleting condition (Supplemental Figure 3I), which may be related to the protective role of the de novo lipid biosynthesis pathway as an electron sink when acetate was depleted and photosynthesis was inefficient, consistent with the induced transcripts involved in photoprotection under the same conditions (Supplemental Figure 4O). Furthermore, our MapMan ontology enrichment analysis showed that the functional group of lipid metabolism and lipid degradation was significantly enriched in differentially regulated transcripts between 25°C and 35°C under the acetate-depleting conditions but not under the constant-acetate condition, suggesting acetate depletion may make the moderate high temperature more stressful (Supplemental Figure 5A, C). Our work provides rich datasets for researchers with interest and expertise in lipid metabolism to further investigate how changes of lipids affect algal heat responses.

Most previous heat treatments in Chlamydomonas used cultures grown with acetate-containing medium in flasks without turbidostatic control (Hemme *et al*., 2014; Rütgers *et al*., 2017). Our results showed that such heat treatment could not last more than 8 h at 35°C or 24 h of 40°C due to acetate-depletion (Fig. 1D). In fact, many previous heat experiments were conducted for a short-term, e.g., 1 h (Légeret *et al*., 2016; Rütgers *et al*., 2017). Using our highly controlled PBR systems, we could not only conduct heat experiments under the 24-h acetate-depleting condition but also during 4-day heating with and without acetate under the turbidostatic control with constant nutrient supply to uniquely reveal the effects of carbon availability, heat duration and intensity on algal heat responses. Additionally, we used constant light in our experimental conditions to focus specifically on the effects of high temperatures. In nature, high temperatures are accompanied by day and night cycles. The interaction among high temperatures, carbon status, and diurnal cycles will be an exciting area in future research.

In summary, by using the model green alga Chlamydomonas and highly controlled cultivation systems in PBRs, we revealed how the availability of an organic carbon source, acetate, interacted with different intensities and durations of high temperatures in photosynthetic cells. Our research revealed the overlooked effects of moderate high temperature of 35°C, which was beneficial with constant carbon supply or detrimental with insufficient or no carbon supply. We also showed that the damaging effects of acute high temperature of 40°C was dominant and independent of carbon availability. Our research not only helped us understand heat responses in photosynthetic cells but also provided insights for high temperature effects on the production of algal biofuels and bioproducts. Our results will also be useful to study how carbon availability affects heat responses in mixotrophic and heterotrophic (reproductive or sink) tissues in land plants.

## Supplementary data

**Supplemental Table 1**: Gene IDs and primers for RT-qPCR analysis in Fig. 2 and Supplemental Fig. 2 and 3.

**Supplemental Dataset 1**. RNA-seq data and plots of each transcript detected, including mean normalized read count plots, mean normalized read count heat map, log_2_(fold change) of mean normalized read counts relative to the pre-heat time point under the same condition, and statistical analysis using False Discovery Rate (FDR).

**Supplemental Dataset 2**. Log_2_(fold change, FC) of mean normalized read counts of groups of transcripts based on MapMan function annotation (MapMan folder) or manual curation (Manual_curation folder). Manual curation was based on the gene lists from this paper Zhang *et al*., 2022*a*). Each line in the plots represent one unique transcript in that function category under one of six conditions. RNA-seq data with normalized read counts were used for this analysis. Log_2_FC is relative to the pre-heat time point under the same condition. Black vertical lines indicated the start of heat treatments under 35°C or 40°C or the equivalent time point at 25°C. ANOVA test was applied to the time course (five time points) of one transcript for each condition. If a transcript differed significantly during the time course (or underwent a global significant change) of one condition, it is colored purple (ANOVA significant); blue lines had no significant changes during the time course. Different intensities of purple and blue colors were used to distinguish different transcripts. Each file opens as a web page. Please use the interactive feature of the figures to see the gene IDs for each panel.

**Supplemental Dataset 3**. MapMan ontology enrichment. Ontology enrichment was performed for transcripts with significantly differential expression in the indicated comparisons using the extended MapMan ontology. The p values were calculated using a hypergeometric test and corrected for multiple testing using the Benjamini-Hochberg method.

**Supplemental Dataset 4**. Normalized RNA-seq read counts.

**Supplemental Dataset 5**. RNA-seq data summary, including gene IDs, MapMan function groups, annotation, adjusted False Discovery Rate (FDR) values for comparisons of different conditions, mean and standard deviation (sd) of normalized read counts.

**Supplemental Dataset 6**. ChlamyCyc Pathway enrichment analysis of differentially regulated transcripts compared between two conditions. The ChlamyCyc annotations (v2023-01-04) and pathway ontology combine KEGG, MapMan, and JGI pathway information (May *et al*., 2009). The p values were calculated using a hypergeometric test (BioFSharp.Stats) and corrected for multiple testing using the Benjamini-Hochberg method (FSharp.Stats).

## Abbreviations

PBRs: Photobioreactors
TAP medium: Tris-acetate-phosphate
TP medium: Tris-phosphate

## Acknowledgements

We would like to thank Dr. Michael Schroda for discussing the initial experimental idea under the acetate-depleting conditions, Dr. Trent Northen for recommending the acetate quantification assay, and Dr. Jaruswan Warakanont for discussion about Chlamydomonas lipid metabolism. We also want to thank the team of the Department of Energy (DOE) Joint Genome Institute (JGI) Community Science Program (Anna Lipzen, Christa Pennacchio, Kerrie Barry, Dr. Jeremy Schmutz, and others) for managing and sequencing our RNA samples.

## Author contributions

RZ designed the experiments. CB and MX operated and maintained the algal growth in photobioreactors. NZ, CB, and MX harvested algal samples at different time points and quantified biomass. NZ and MX performed acetate quantification assay. NZ performed RT-qPCR analysis and extracted RNA for RNA-seq. CB and NZ performed light microscope imaging. CB, MX, and NZ quantified cell density, cell size, and chlorophyll contents, and recorded OD_608_/OD_720_ readings. CB and MX performed 4-day heating experiments and quantified growth rates as well as cell parameters. CB analyzed PSII data. BV, TM, RZ did RNA-seq analysis. EMM did initial analysis of the acetate pathways. NZ, BV, CB, RZ, MX, and EMM prepared the figures. BV wrote the RNA-seq method and related figure legends. RZ wrote the rest of the manuscript. RZ, EMM, BV, and CB revised the manuscript.

## Conflict of interest

No conflict of interest declared.

## Funding statement

This work was supported by the start-up funding from the Donald Danforth Plant Science Center (DDPSC) and the DOE Biological & Environmental Research (BER) grant (DE-SC0020400) to RZ. EMM was supported by the William H. Danforth Fellowship in Plant Sciences and the DDPSC start-up funding (to RZ). BV and TM were supported by the Germany funding TRR 175 D02 (Project No: 270050988). The RNA-seq service support was from the Department of Energy (DOE) Joint Genome Institute (JGI) Community Science Program (CSP, 503414) to RZ. The NovaSeq for RNA-seq (proposal: 10.46936/10.25585/60001126) was conducted by the U.S. Department of Energy Joint Genome Institute (https://ror.org/04xm1d337), a DOE Office of Science User Facility, is supported by the Office of Science of the U.S. Department of Energy operated under Contract No. DE-AC02-05CH11231.

## Data availability

All data supporting the findings of this study are available within the paper and within its supplementary materials published online. Raw RNA-seq data was deposited to Sequence Read Archive (SRA) following the Data Policy of the Joint Genome Institute (JGI) with Sequencing Project ID: 1377265.

---

**Supplemental Fig. 1.**
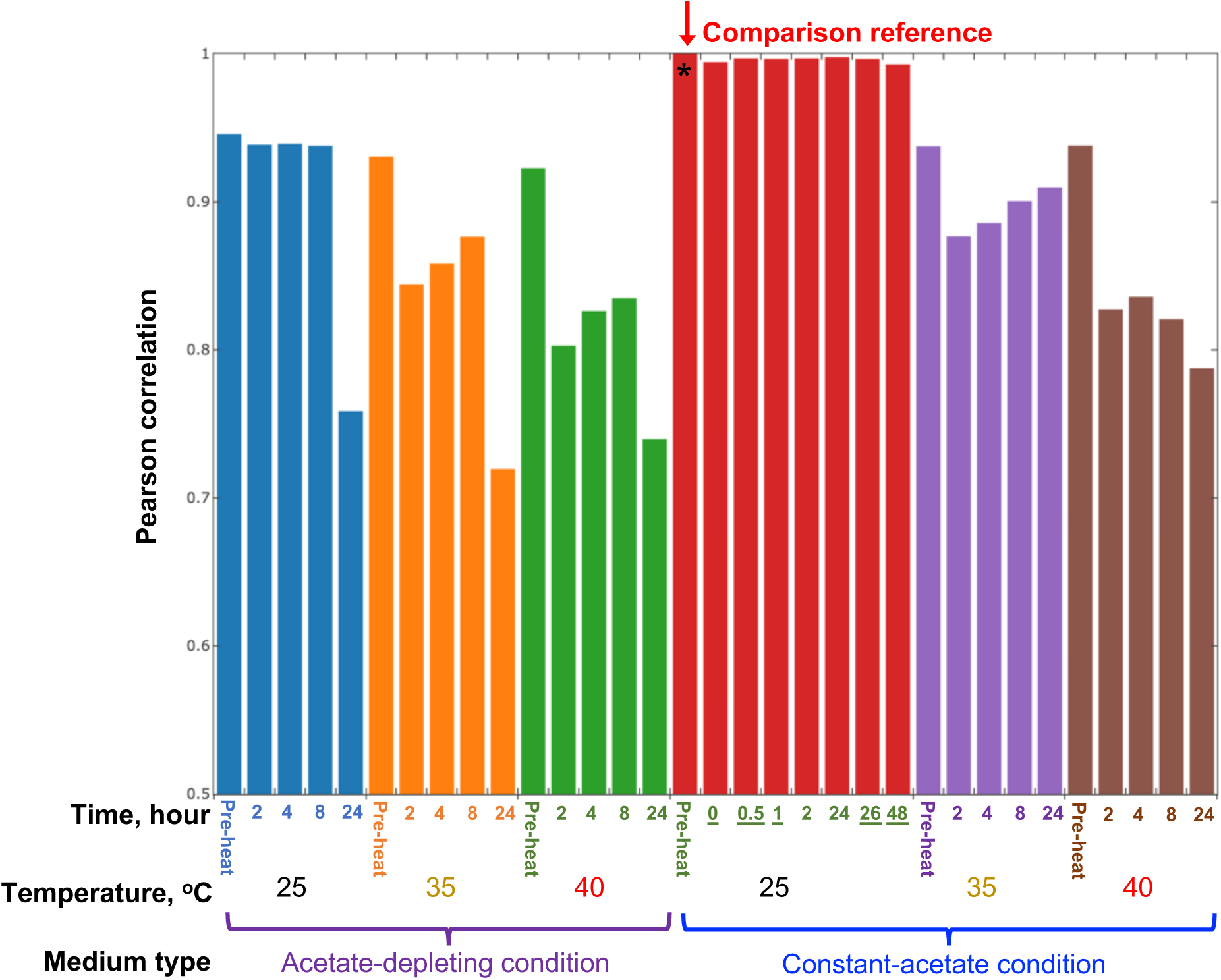
RNA-seq data from algal samples harvested from photobioreactors (PBRs) under the constant-acetate-25°C with turbidostatic control were highly correlated with each other. The Pearson correlations were generated by comparing all RNA-seq samples to the first sample (marked by *) of the constant-acetate-25°C condition. Under constant acetate, 25°C, and light (constant-acetate-25°C) with turbidostatic control in PBRs, the algal growth rates and cell physiologies stay constant (Fig. 5). We had RNA-seq data for 8 time points under the constant-acetate-25°C condition, and these samples were highly correlated with each other (with Pearson correlation ≥ 0.993). The 0 h of the constant-acetate-25°C condition corresponds to the time when PBRs reach expected high temperatures during heat treatments, which takes about 30 min from 25°C to 35°C or 40°C. All pre-heat samples were harvested from PBRs under the same constant-acetate-25°C condition with turbidostatic control and these RNA-seq samples had Pearson correlation > 0.95. Because the cell status and transcriptome stay steady under the constant-acetate-25°C, we used the five extra time points under the constant-acetate-25°C condition (with underlines) to impute the 2 h and 4 h time points under the constant-acetate-25°C condition. See method for details.

**Supplemental Fig. 2.**
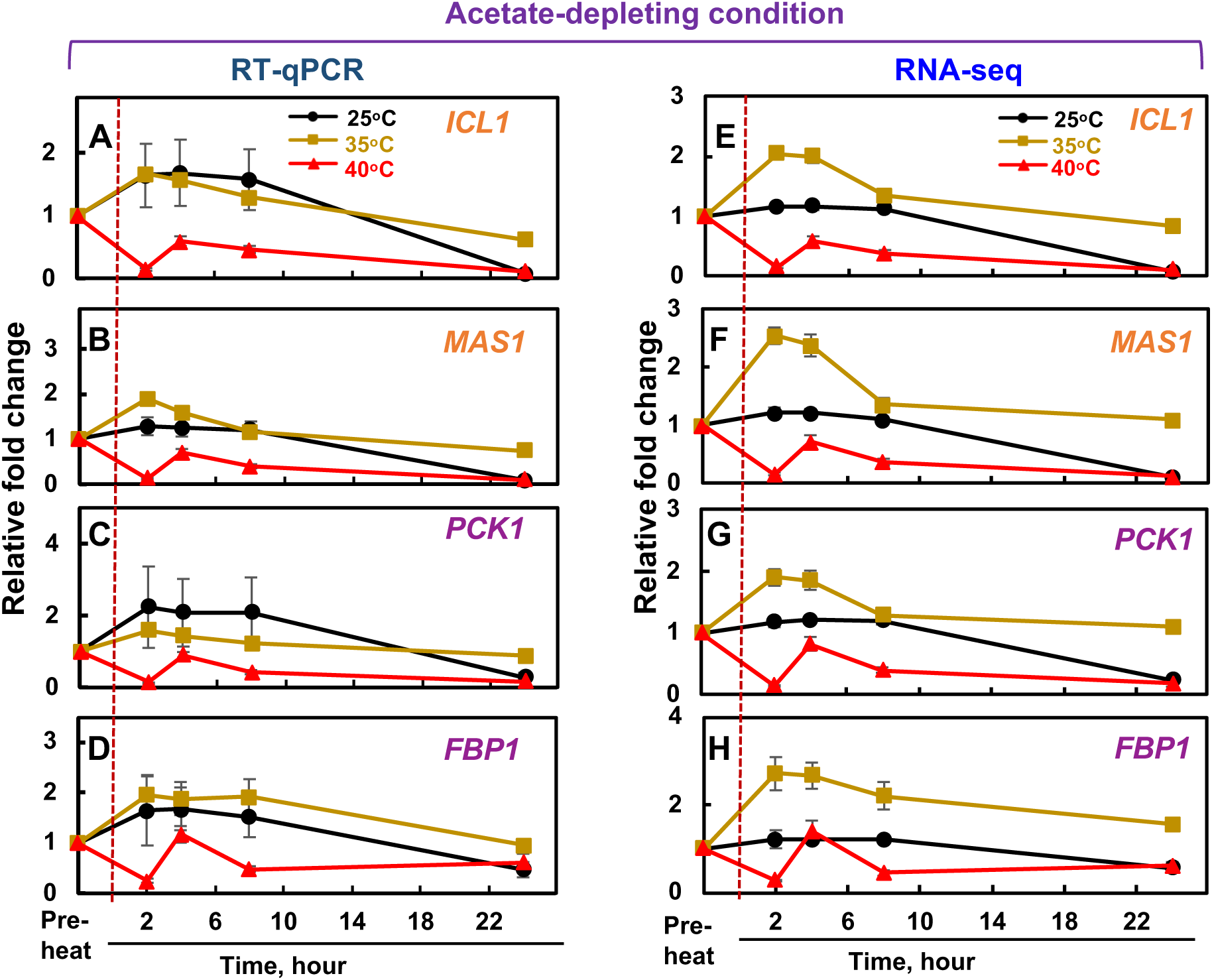
RT-qPCR analysis was consistent with RNA-seq results. Fold changes of four selected transcripts involved in the glyoxylate cycle (*ICL1* and *MAS1*) and gluconeogenesis cycle (*PCK1* and *FBP1*) as in Fig. 2A were calculated based on RT-qPCR (**A-D**) or RNA-seq (**E-H**) results. The red dashed lines mark the start of heat or the corresponding time point at constant 25°C. For RT-qPCR results, the relative fold changes were calculated by normalizing the transcript expression to the reference genes *CBLP*, *EIF1A* and the pre-heat level. For RNA-seq results, the relative fold changes were calculated based on normalized RNA-seq read counts relative to the pre-heat level. Values are mean ± SE, n = 3 biological replicates. Glyoxylate cycle key enzymes: ICL1, isocitrate lyase; MAS1, malate synthase. Gluconeogenesis key enzymes: PCK1, phosphoenolpyruvate carboxykinase; FBP1, fructose-1,6-bisphosphatase (See all gene IDs and RT-qPCR primers in Supplemental Table 1).

**Supplemental Fig. 3.**
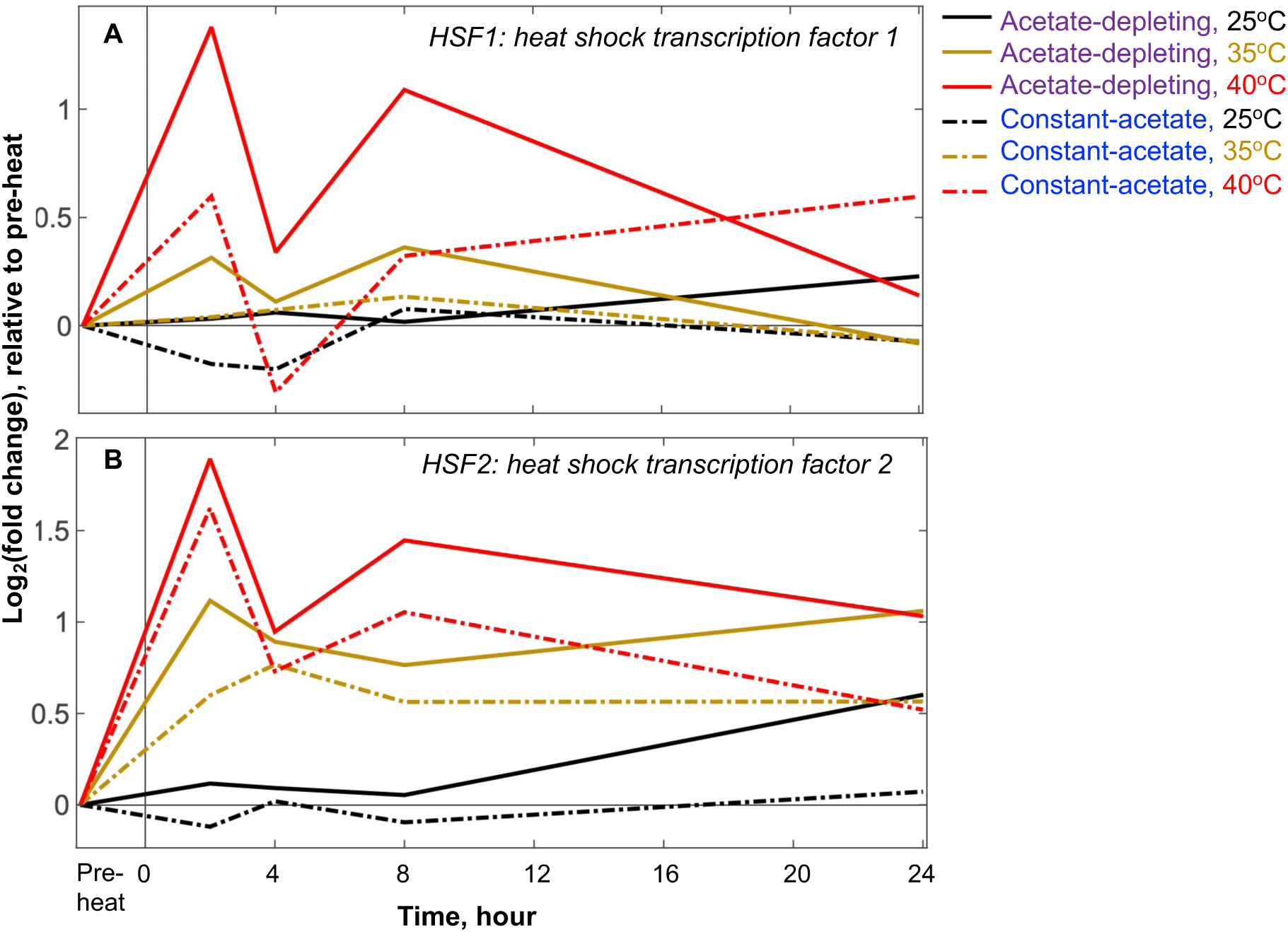
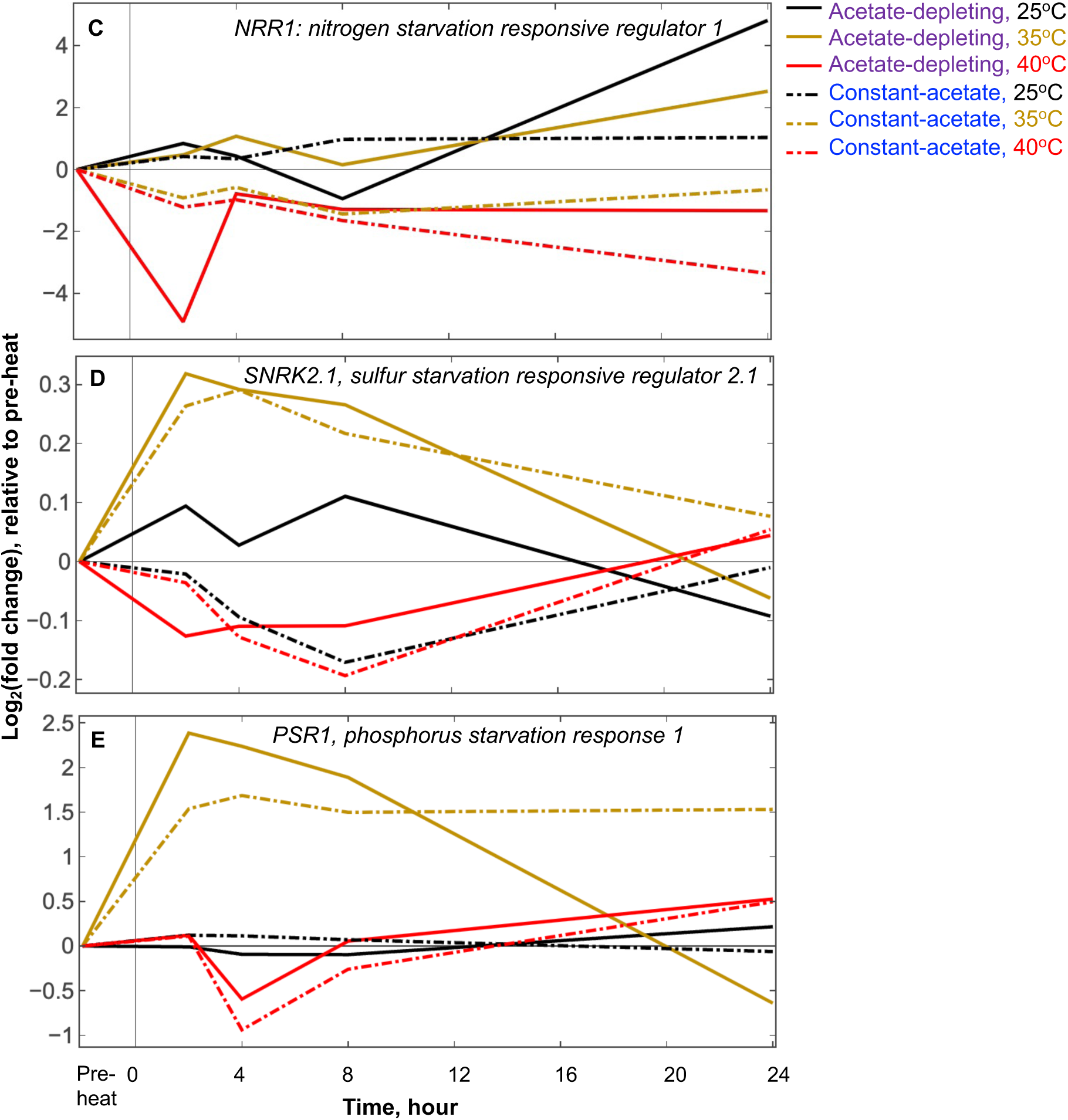
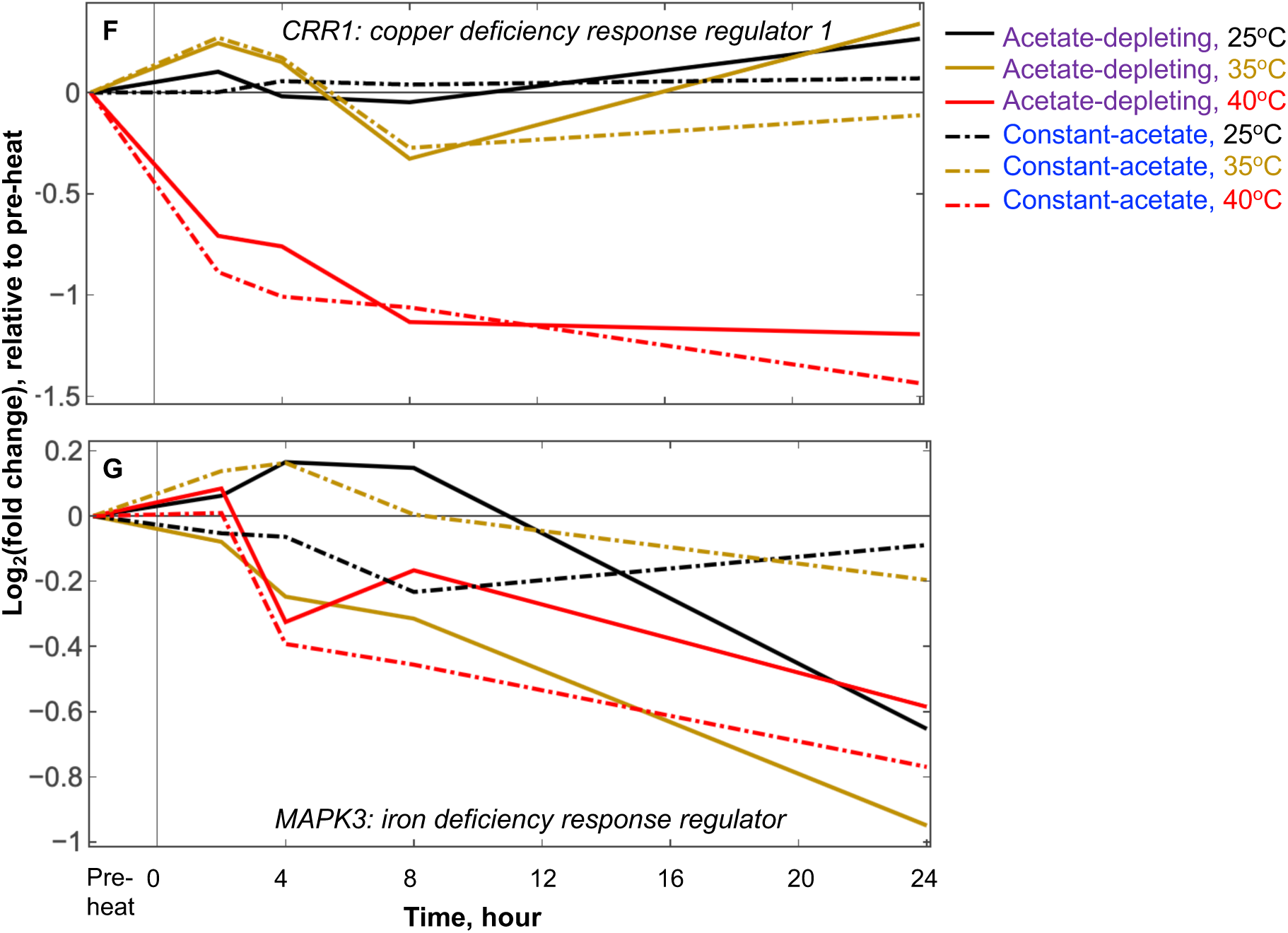
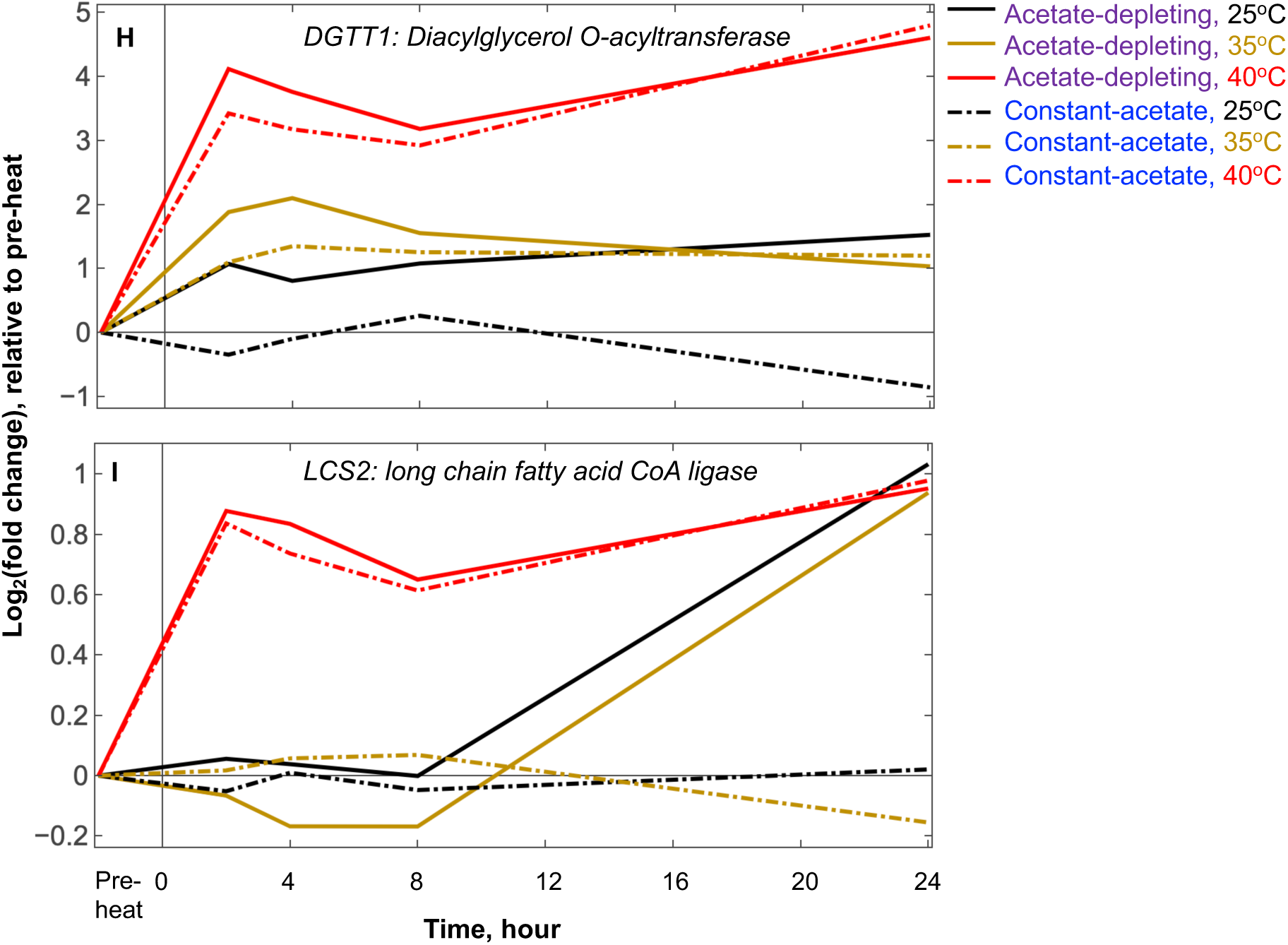
Log_2_(fold change) of the indicated transcript relative to pre-heat under different conditions. Log_2_(fold change) were determined on the mean normalized transcript abundance, relative to the pre-heat time point. Each line represents one condition as indicated by the legend. The black vertical lines mark the start of heat or the corresponding time point at constant 25°C. Solid lines for the acetate-depleting condition and dashed lines for the constant-acetate condition. Black, brown, red lines for 25°C, 35°C, and 40°C temperature treatments, respectively. See all gene IDs in Supplemental Table 1 and Supplemental Dataset 1 for bigger pictures and more information.

**Supplemental Fig. 4.**
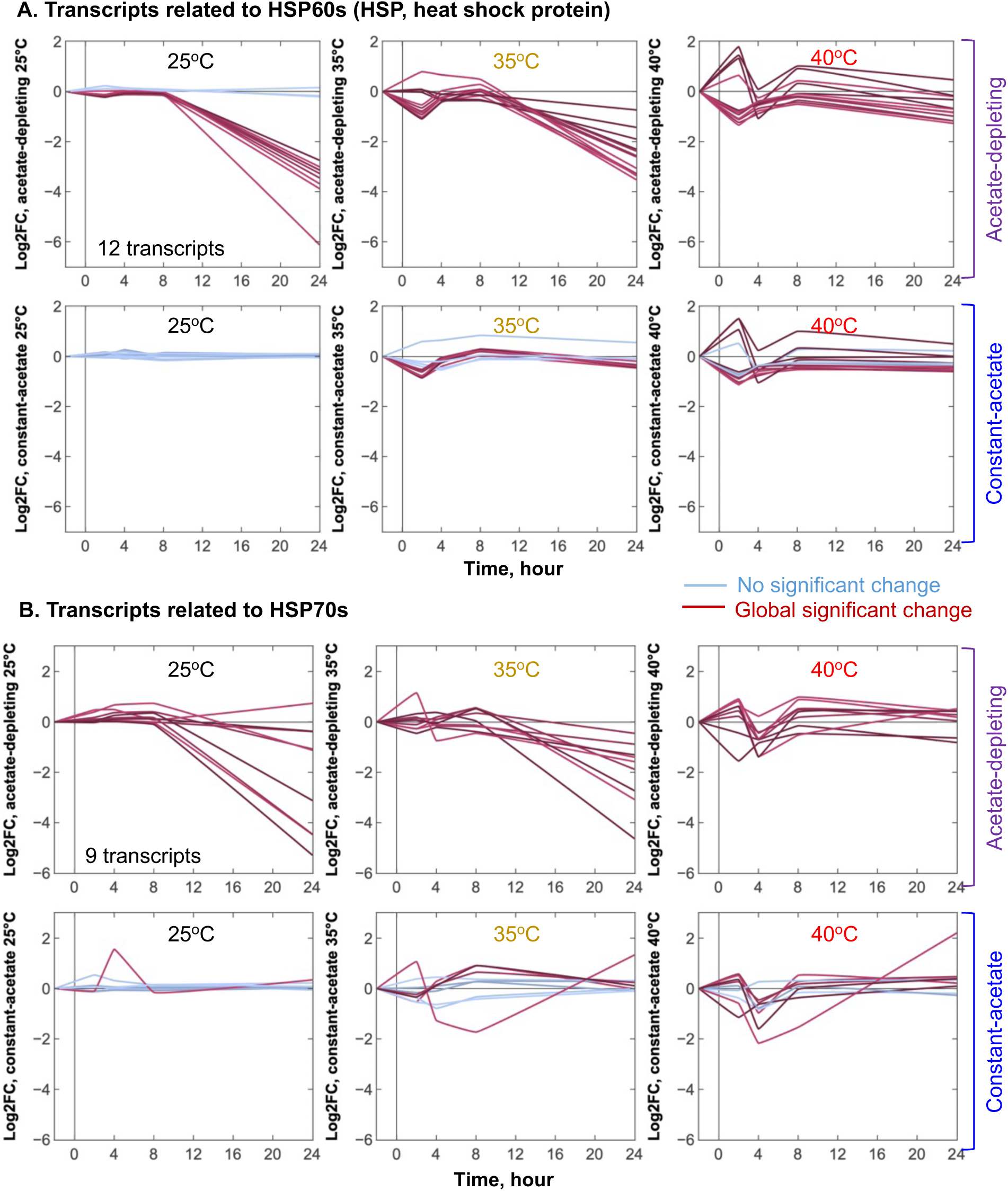
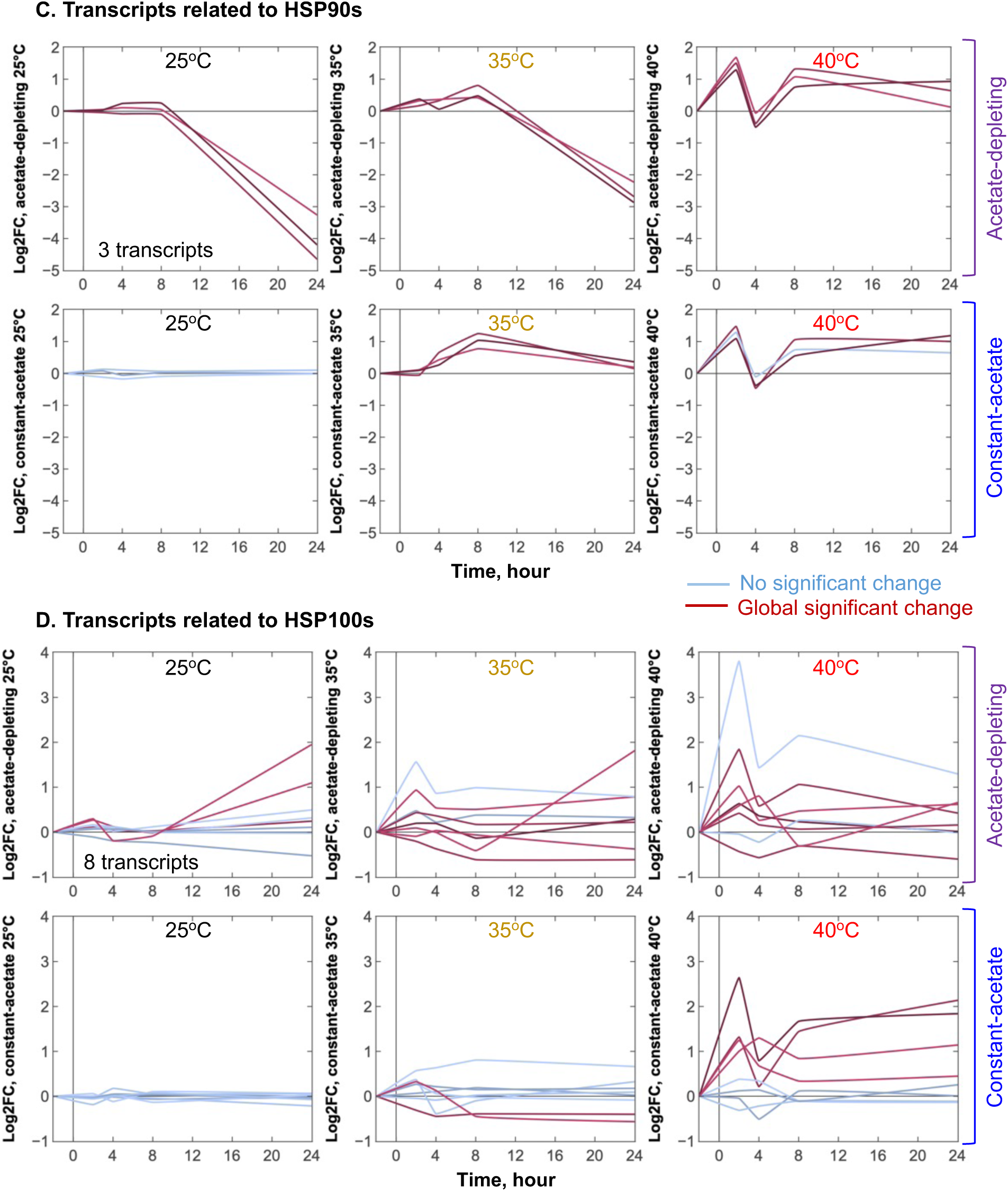
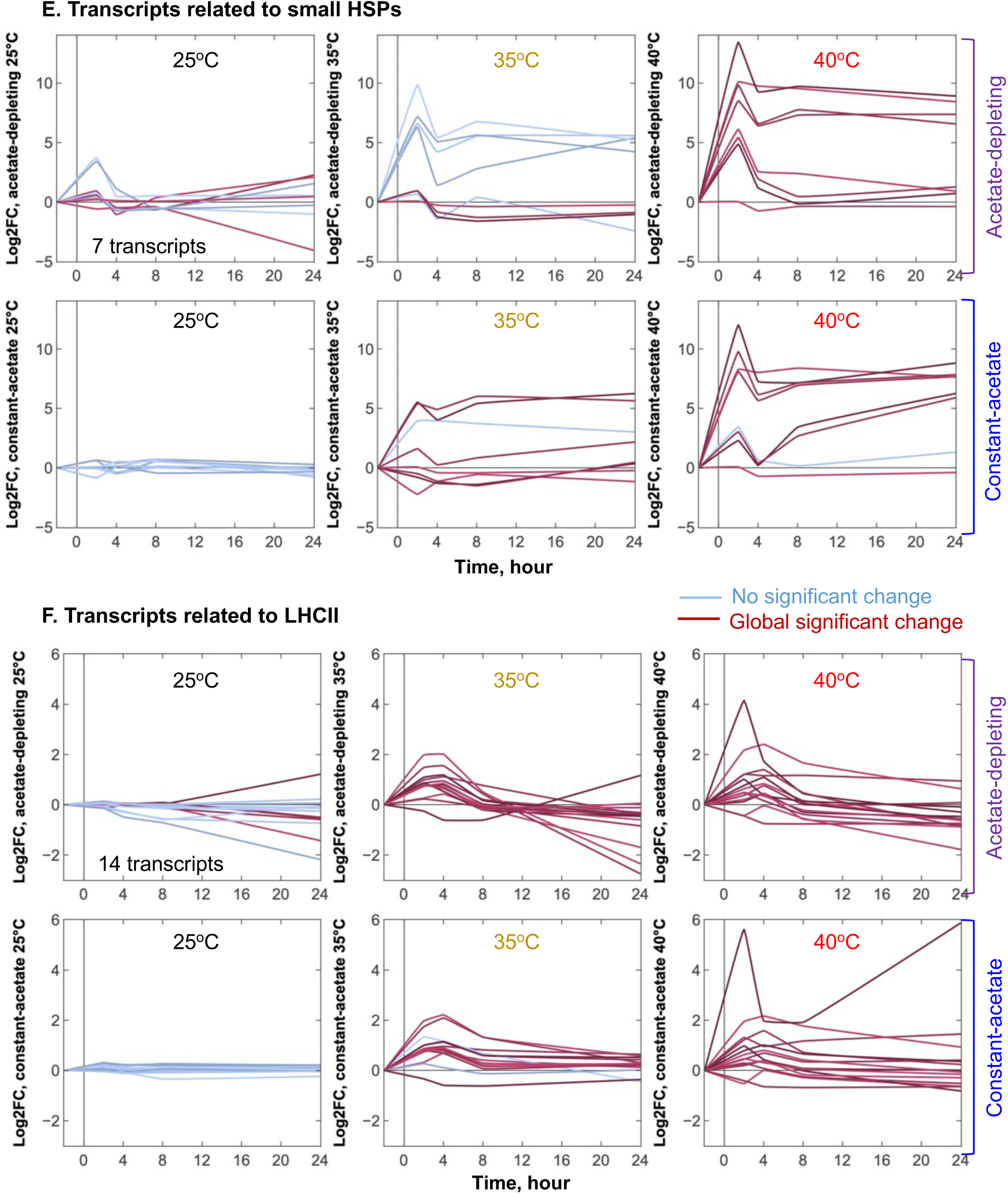
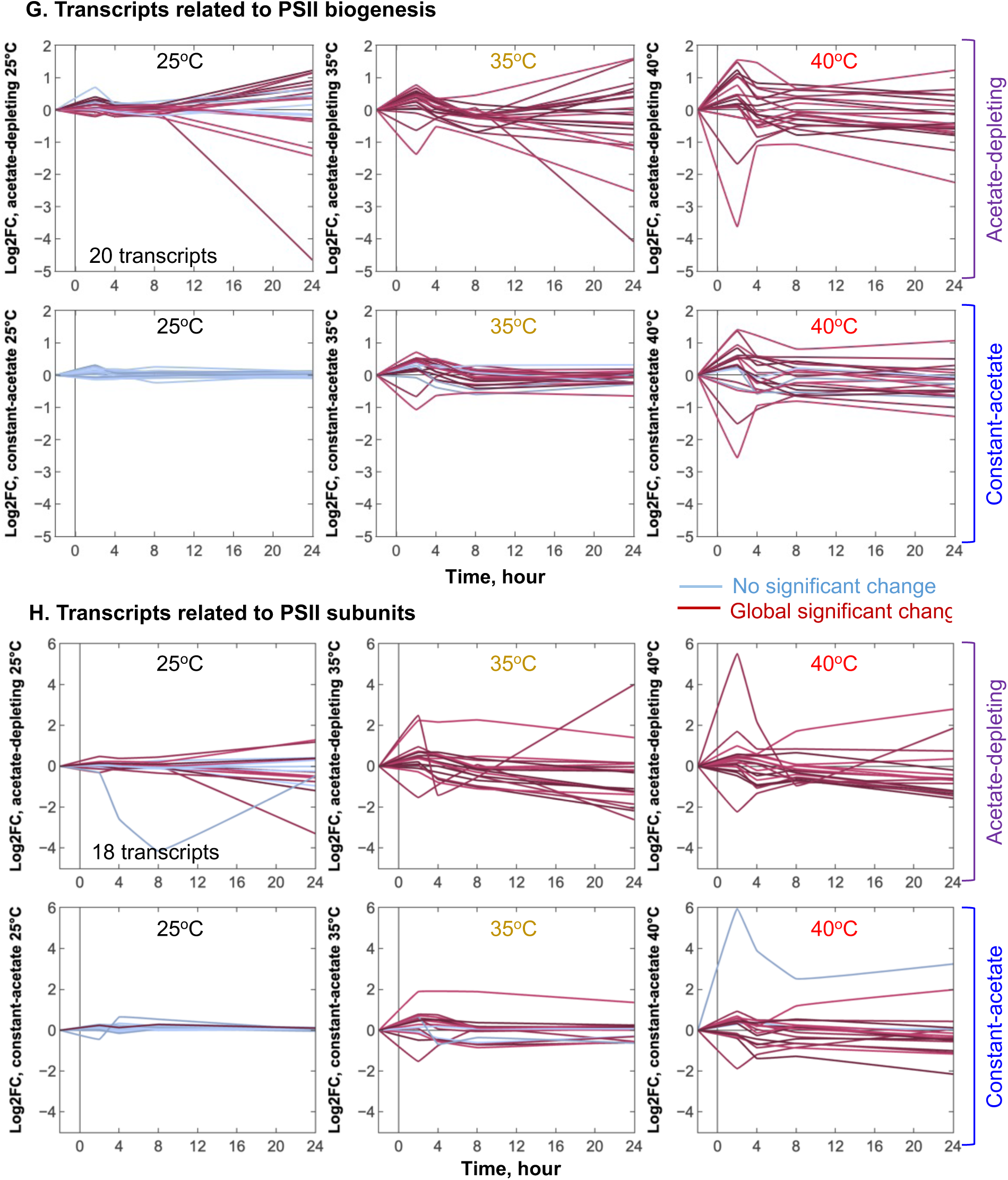
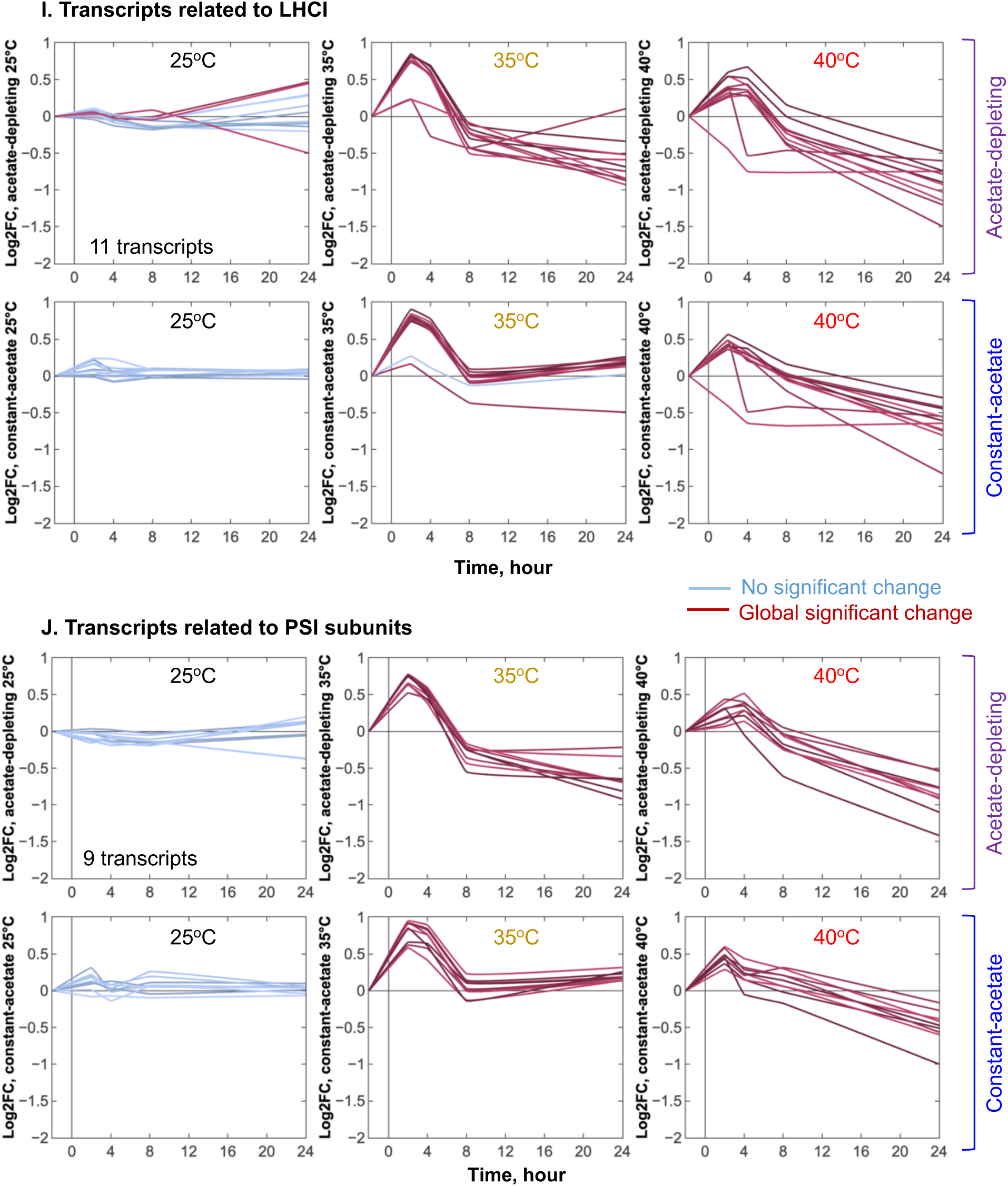
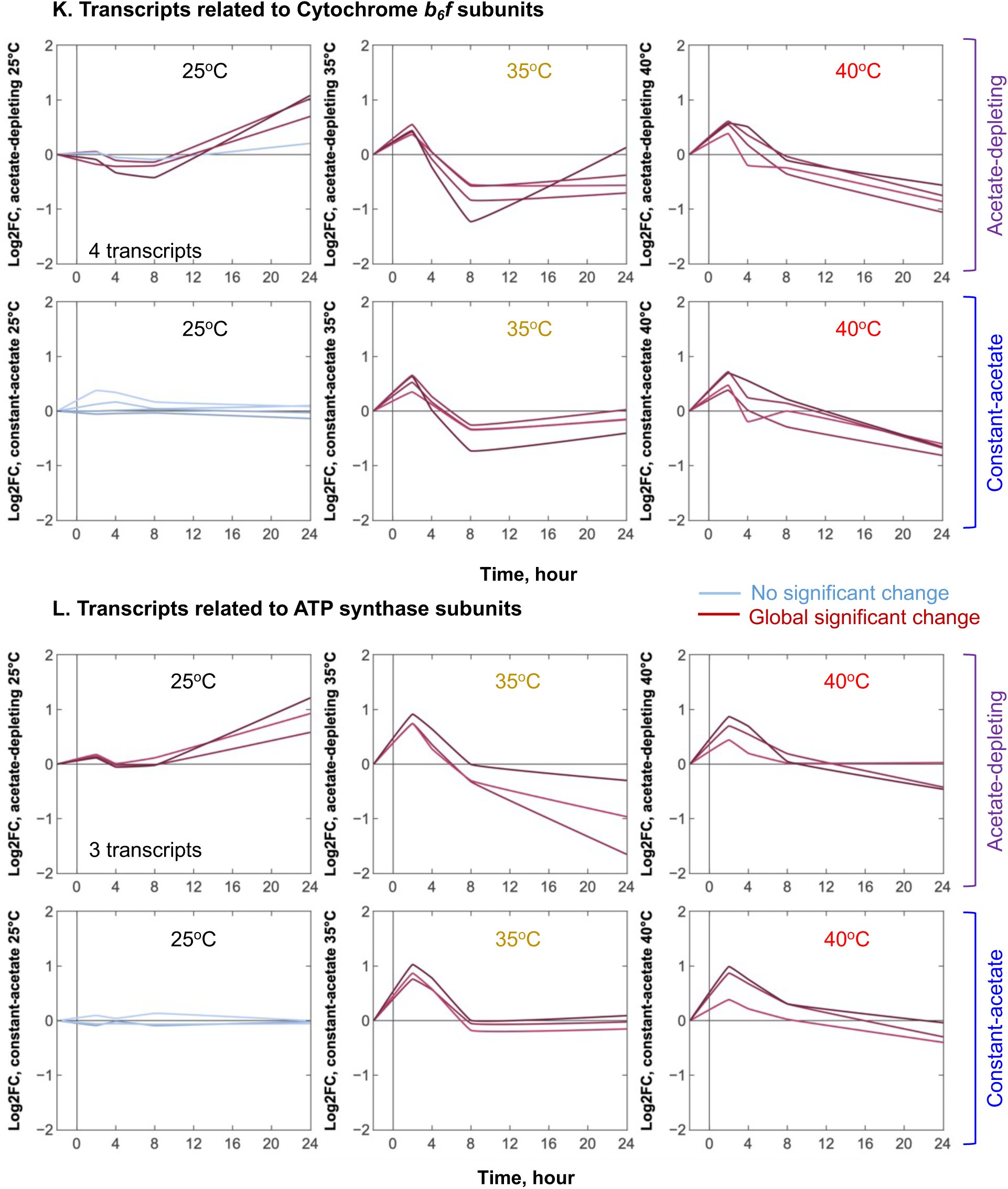
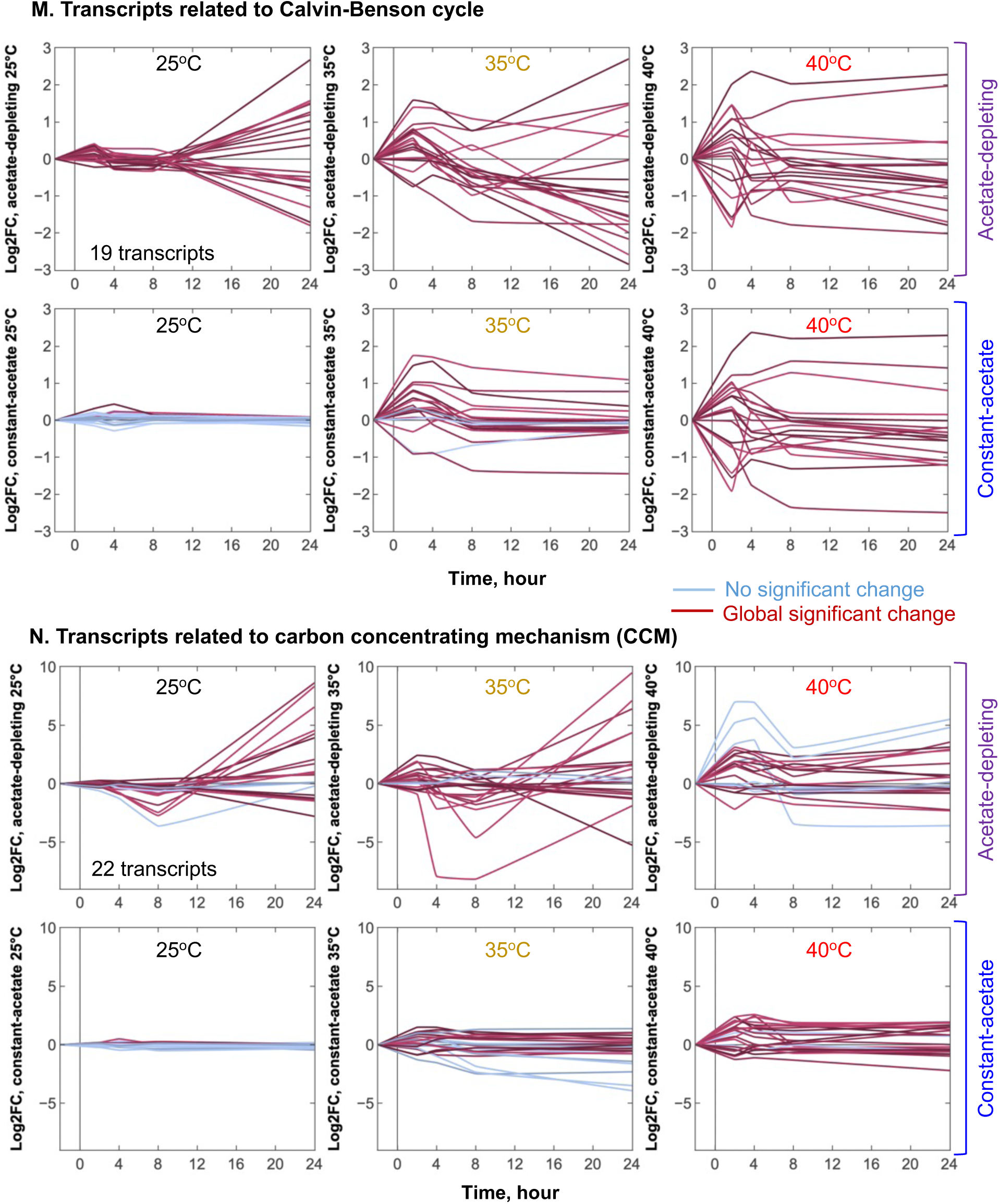
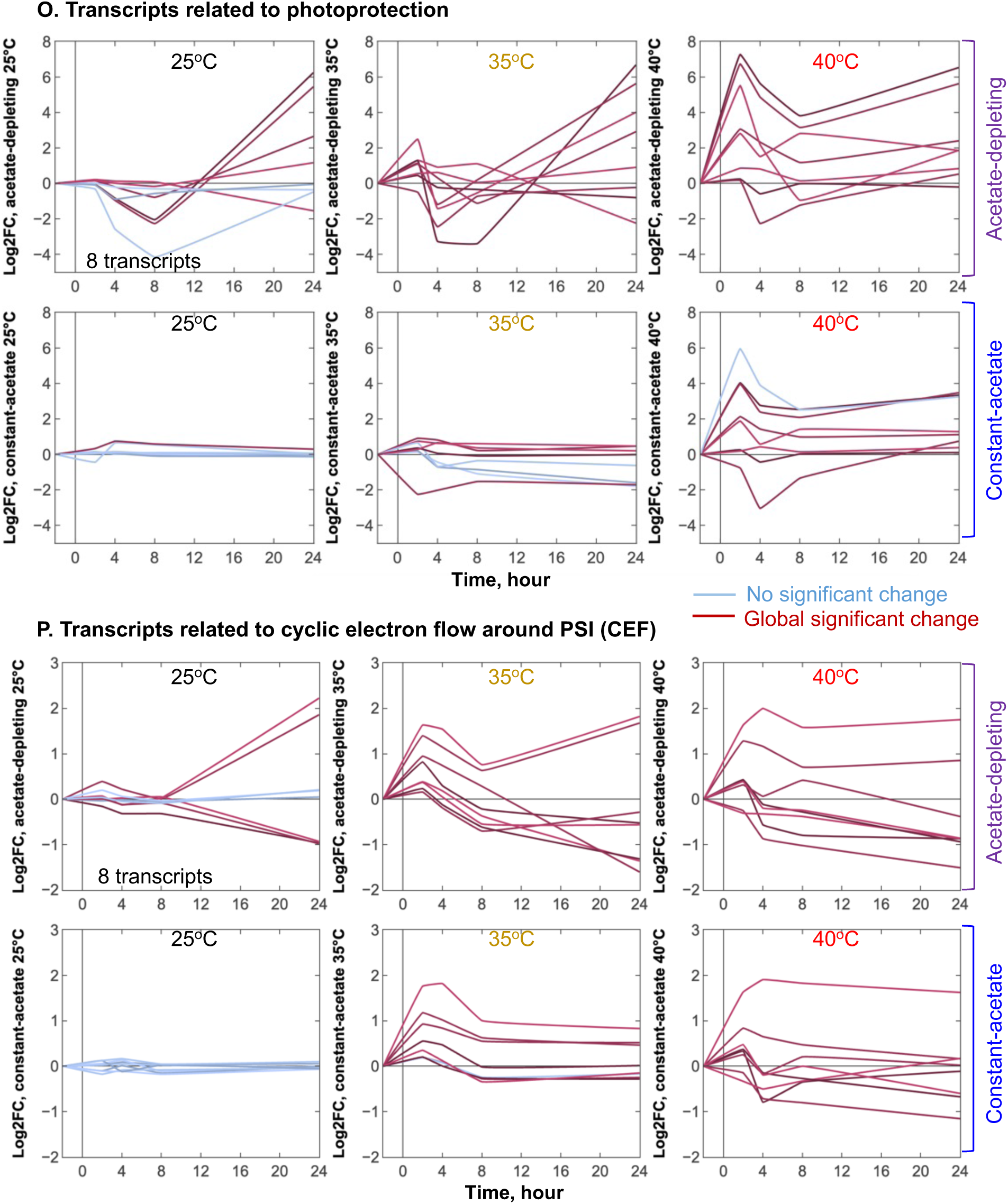
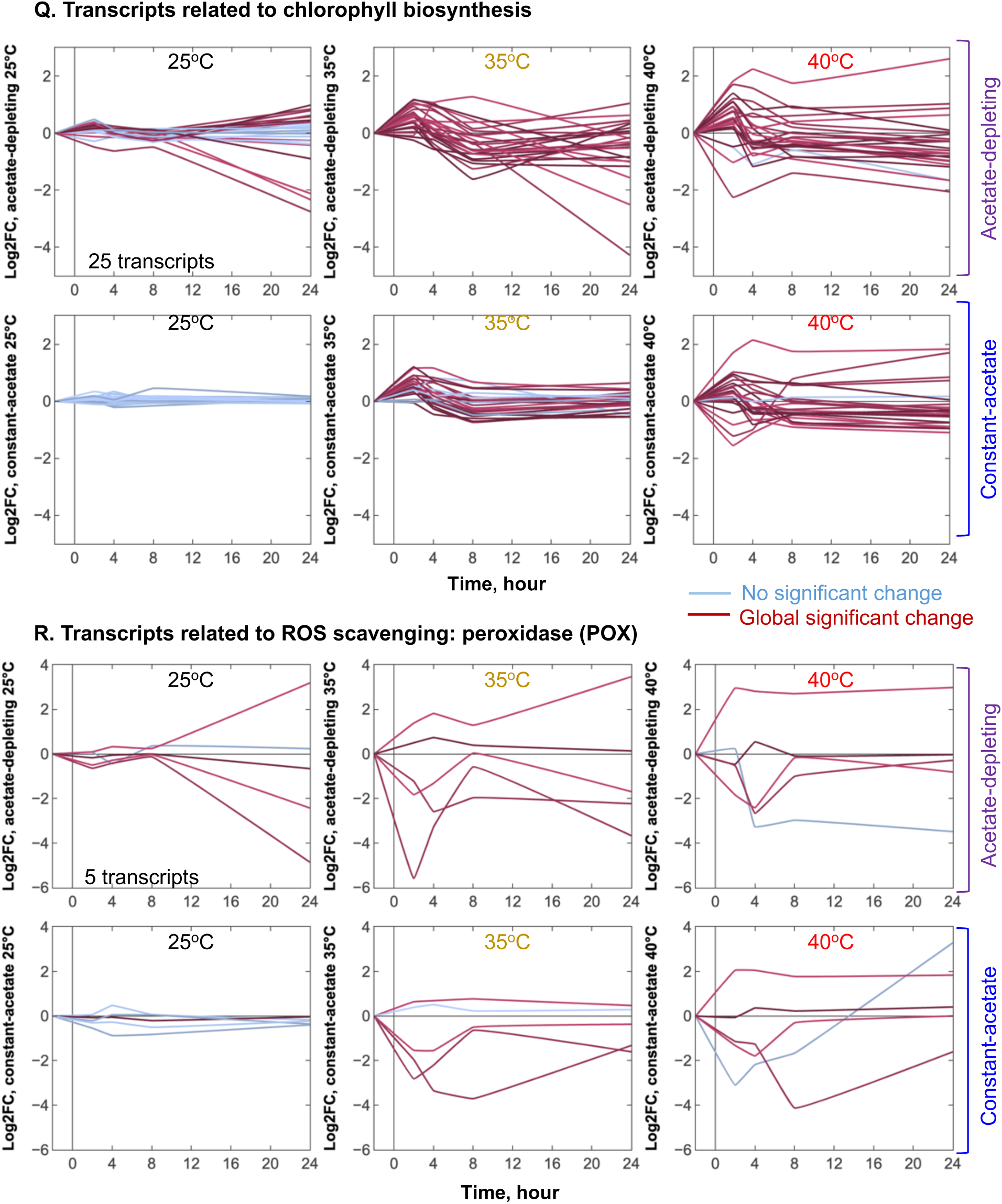
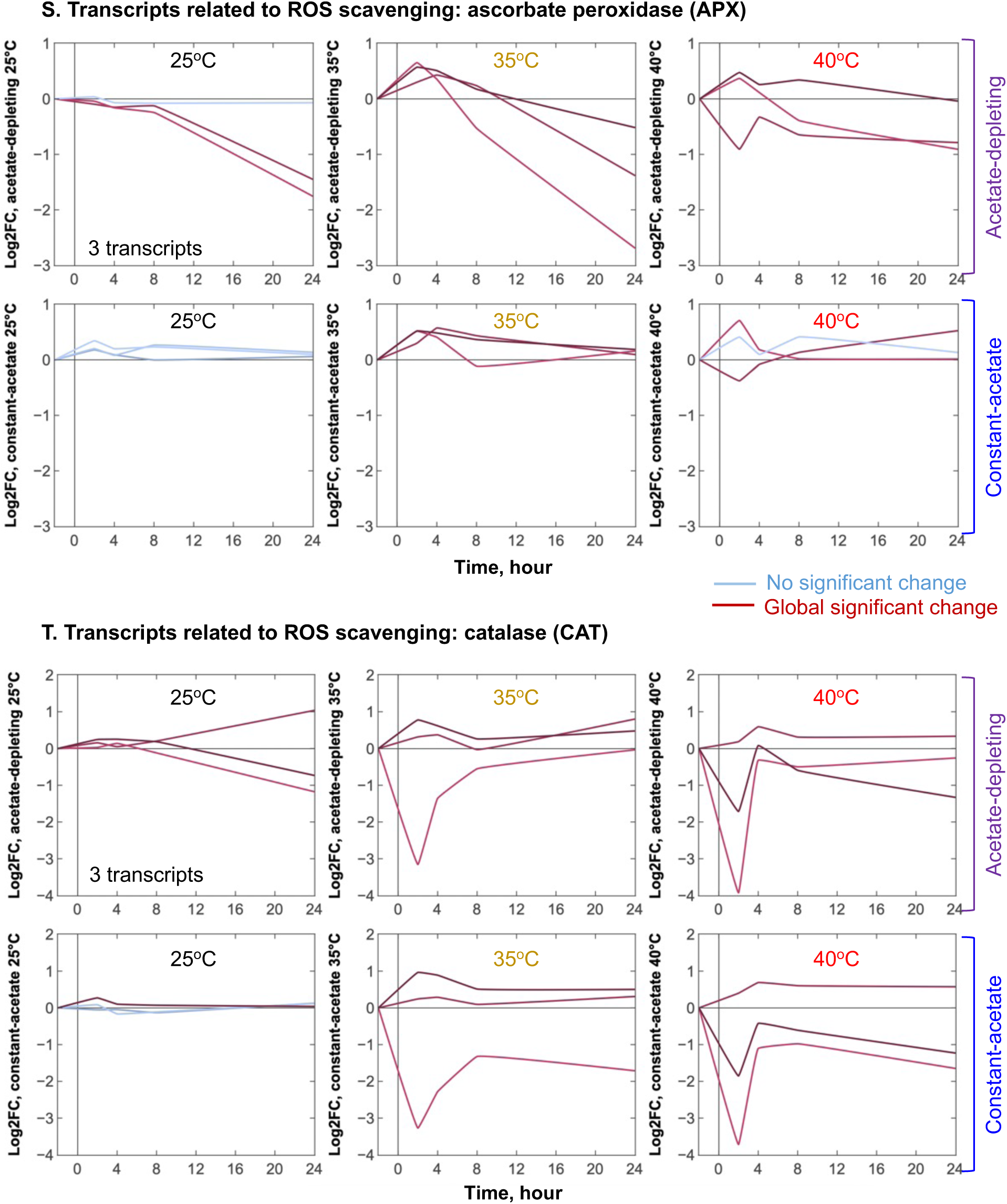
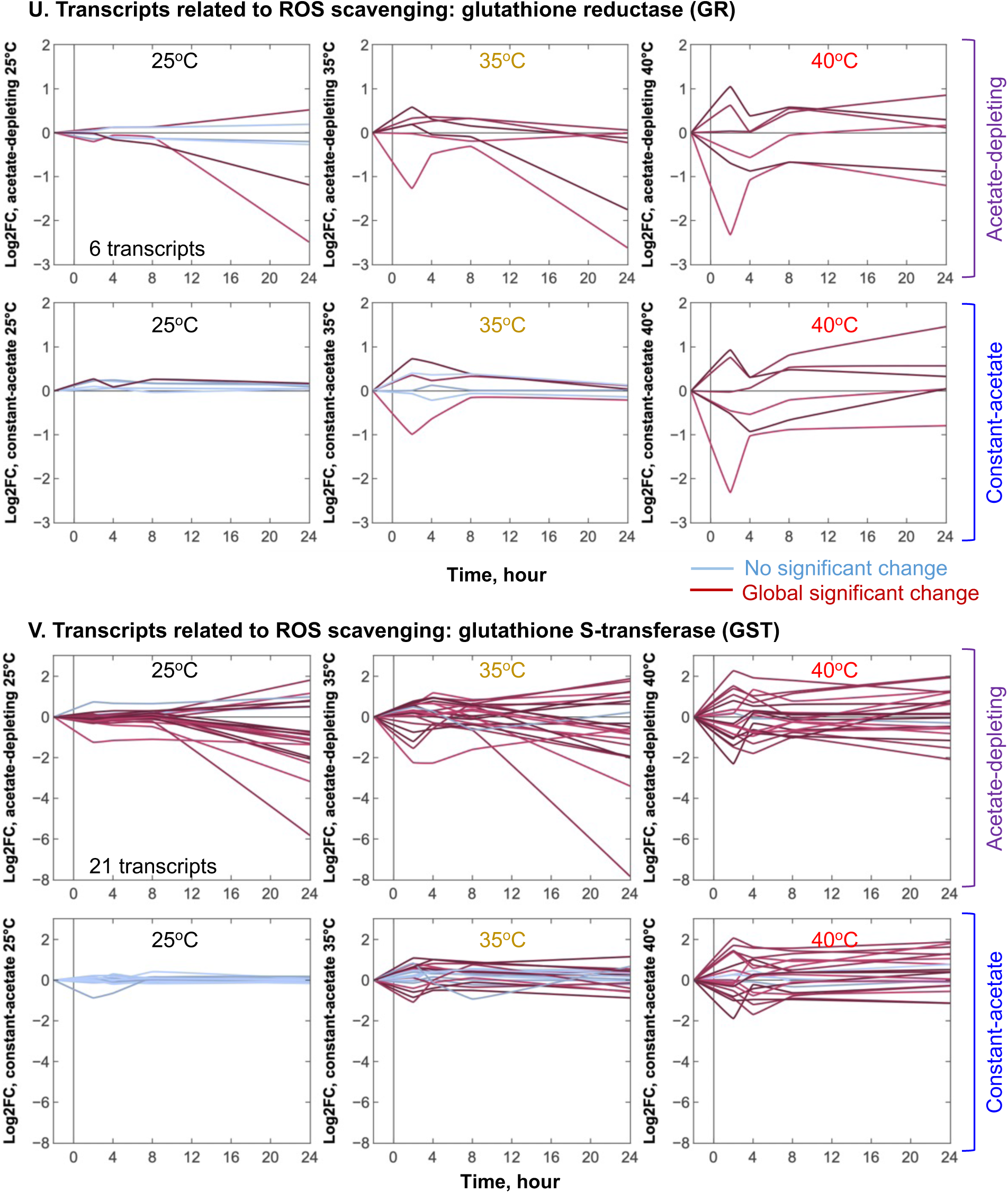
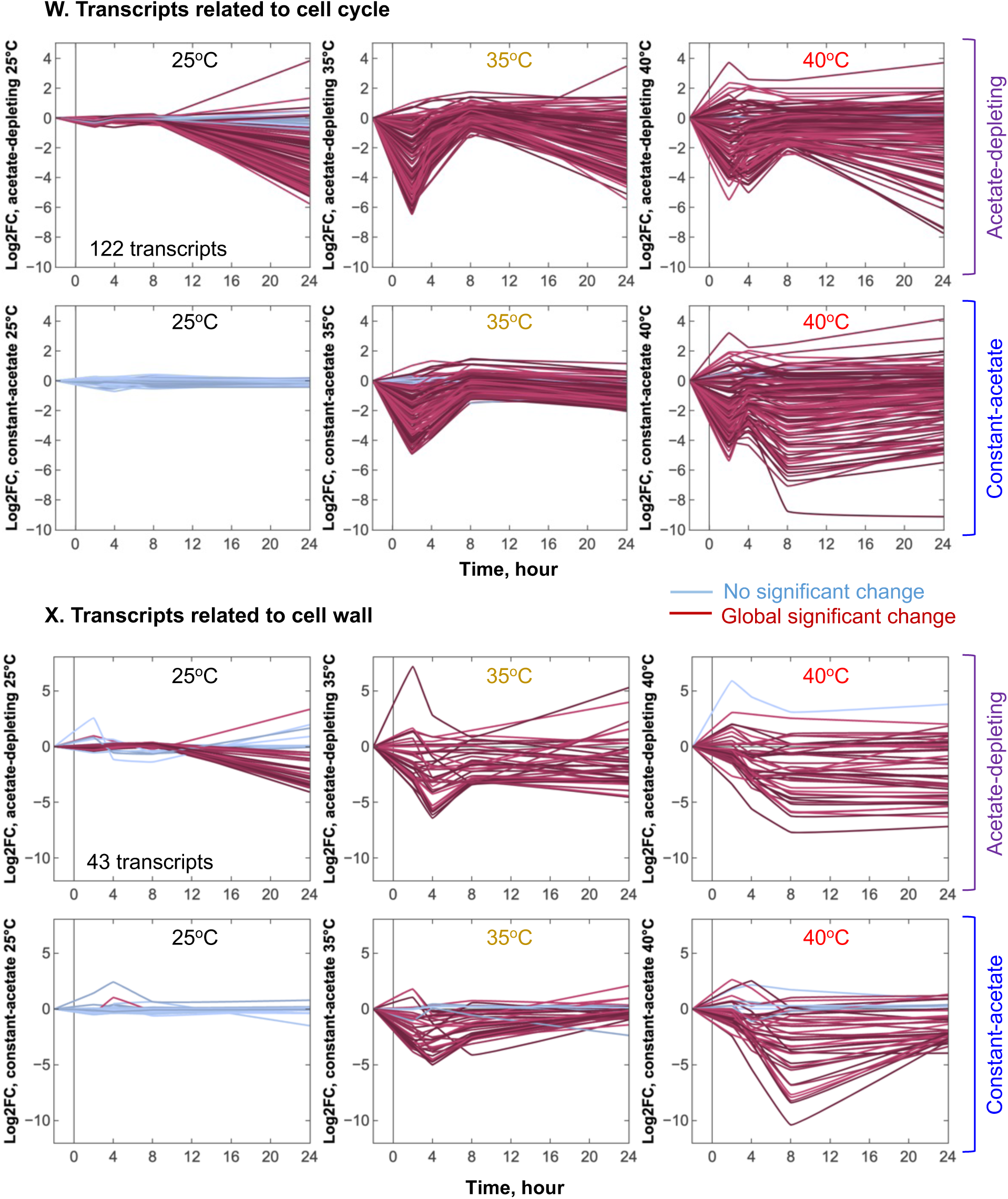
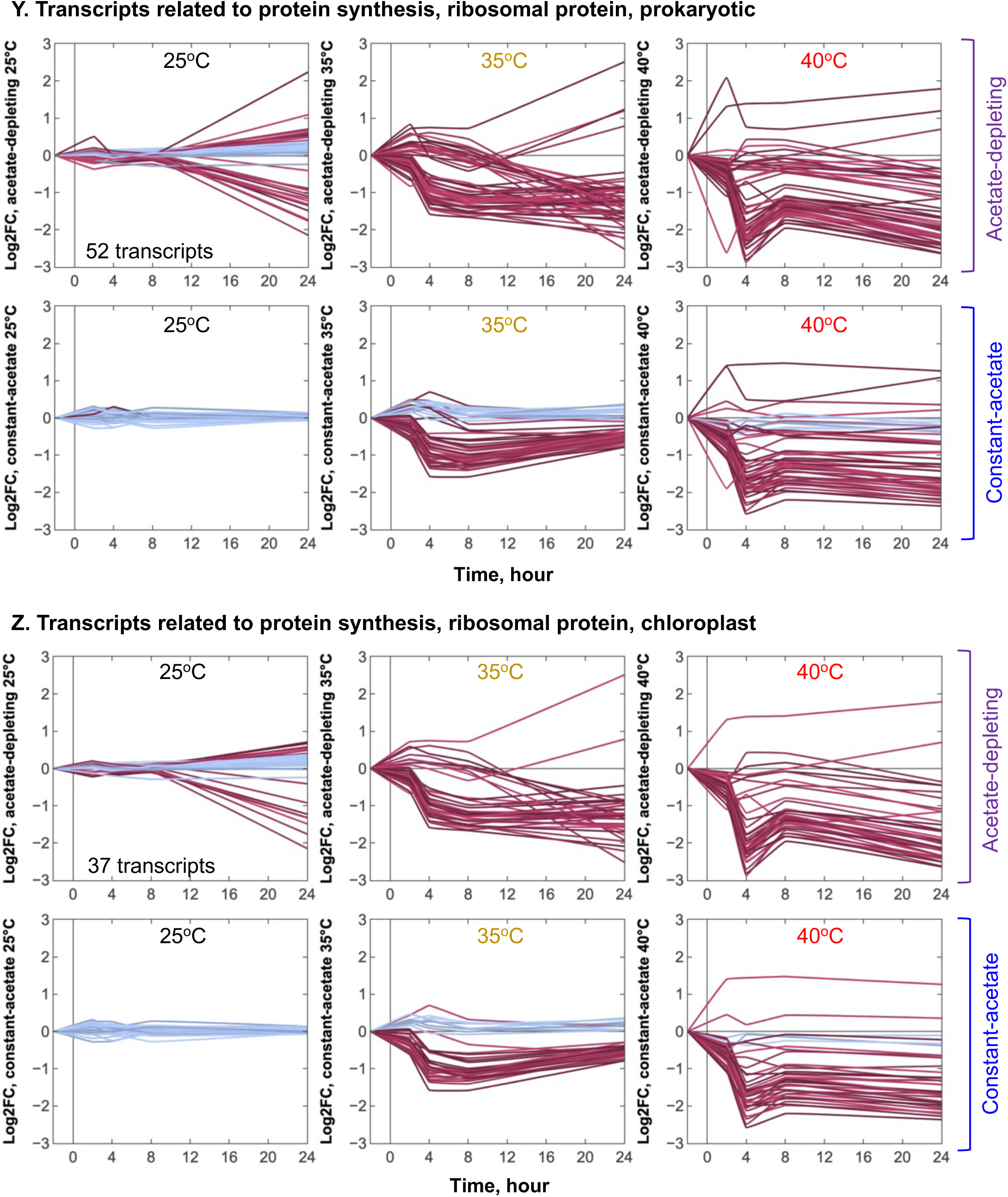
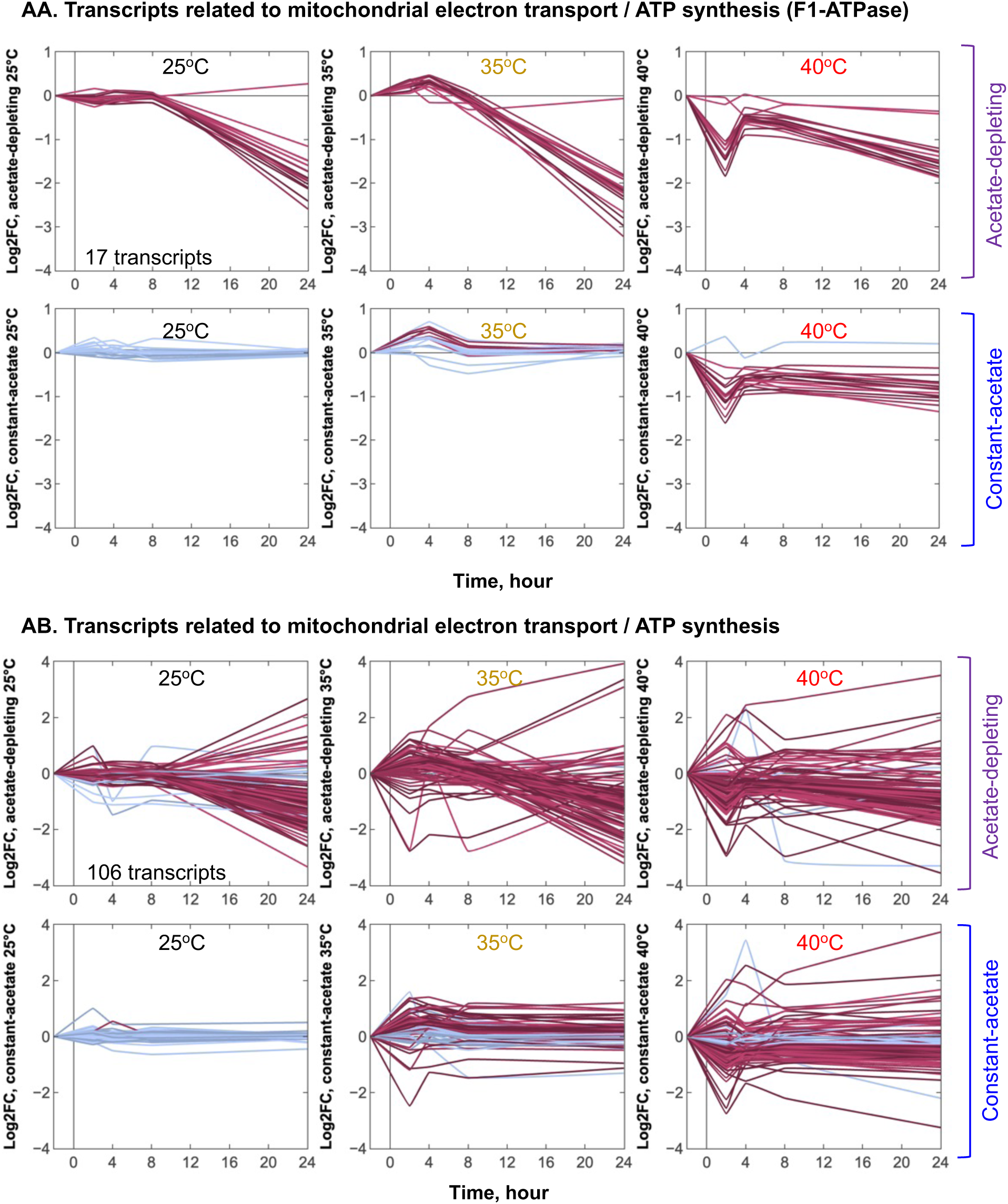
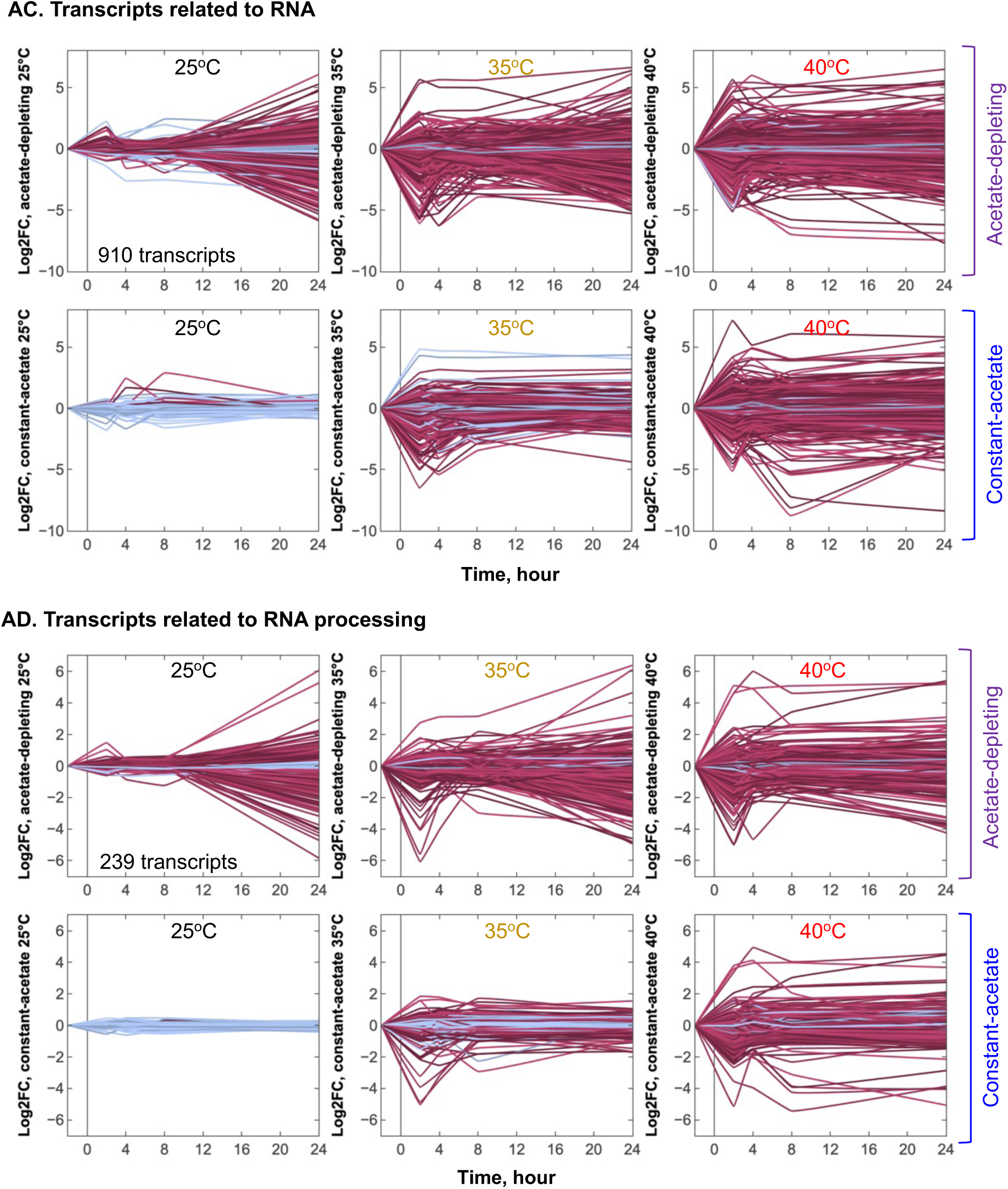
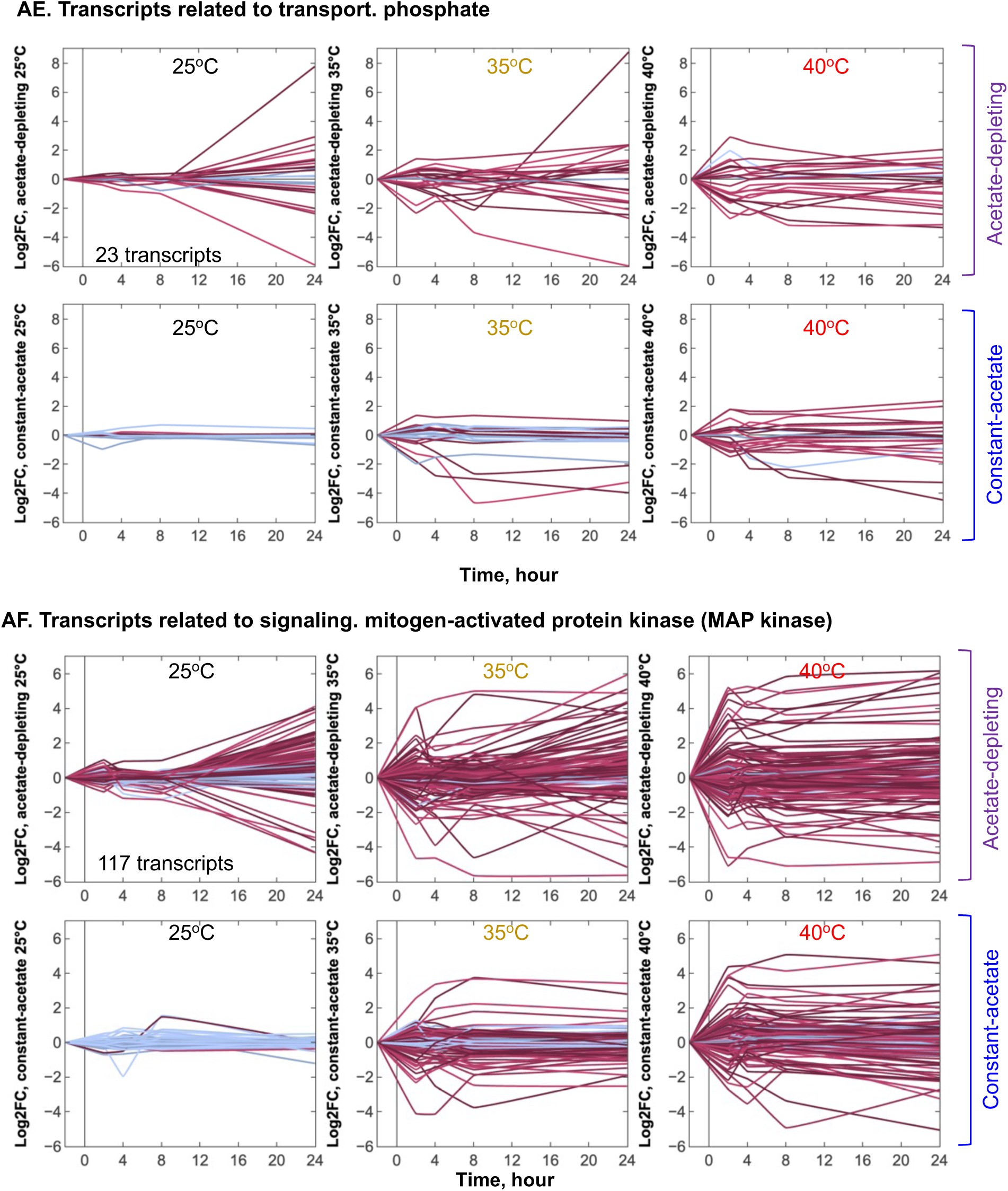
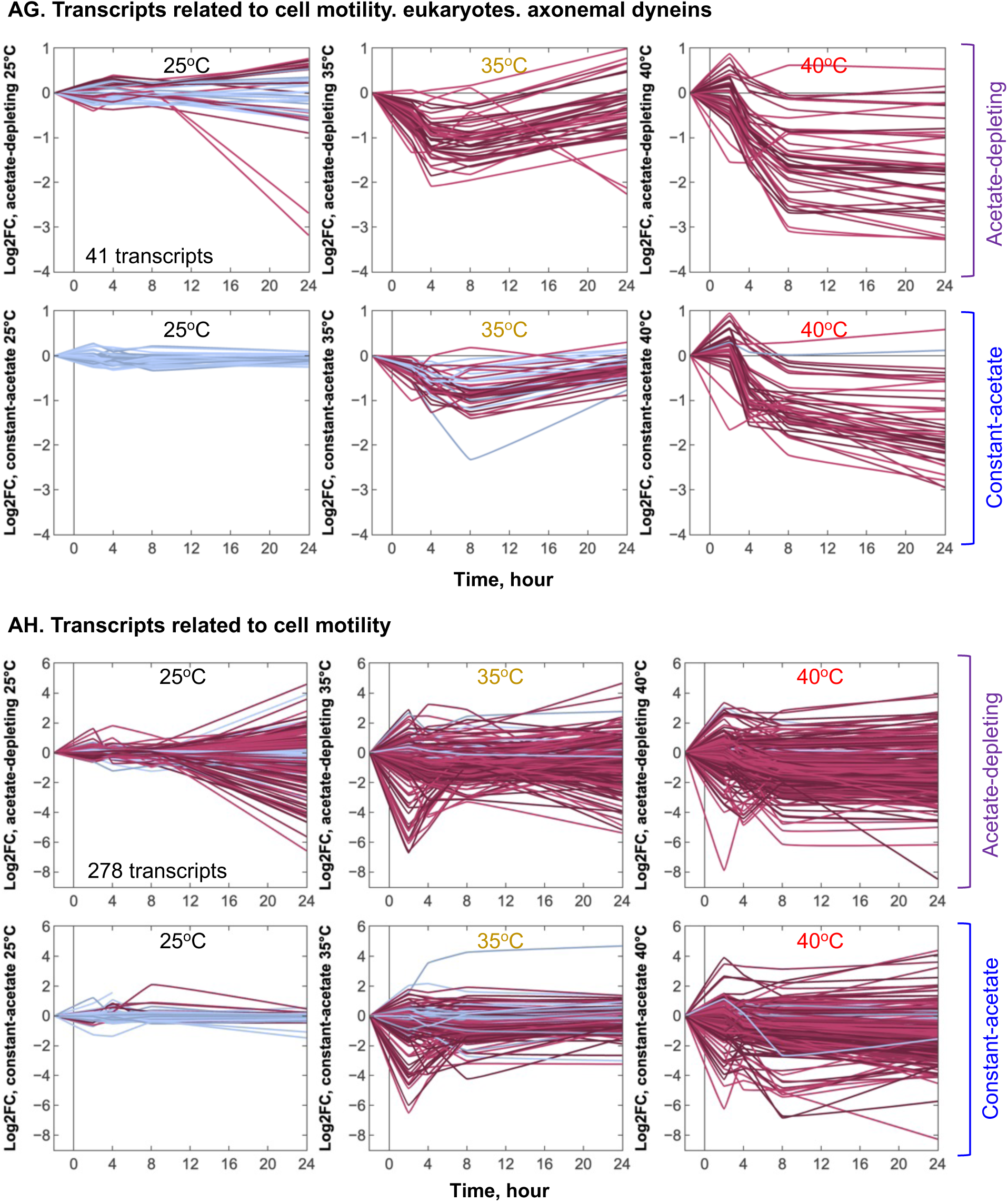
Log_2_(fold change, FC) of mean normalized read counts of groups of transcripts based on MapMan function annotation or manual curation. Plots are Log_2_FC of transcripts related to the indicated function groups under six different treatments, as compared to the pre-heat time points. RNA-seq data with normalized read counts were used for this analysis. Black vertical lines mark the start of heat treatments or the corresponding time point at constant 25°C. Each line represents one unique transcript in the indicated function group. ANOVA test was applied to the time course (five time points) of one transcript for each condition. If a transcript differed significantly during the time course (or underwent a global significant change) of one condition, it is colored purple (ANOVA significant); blue lines had no significant changes during the time course. Different intensities of purple and blue colors were used to distinguish different transcripts. The total number of transcripts plotted in each function group is marked on the top left panel. See interactive figures with gene IDs and annotations in Supplemental Dataset 2

**Supplemental Fig. 5.**
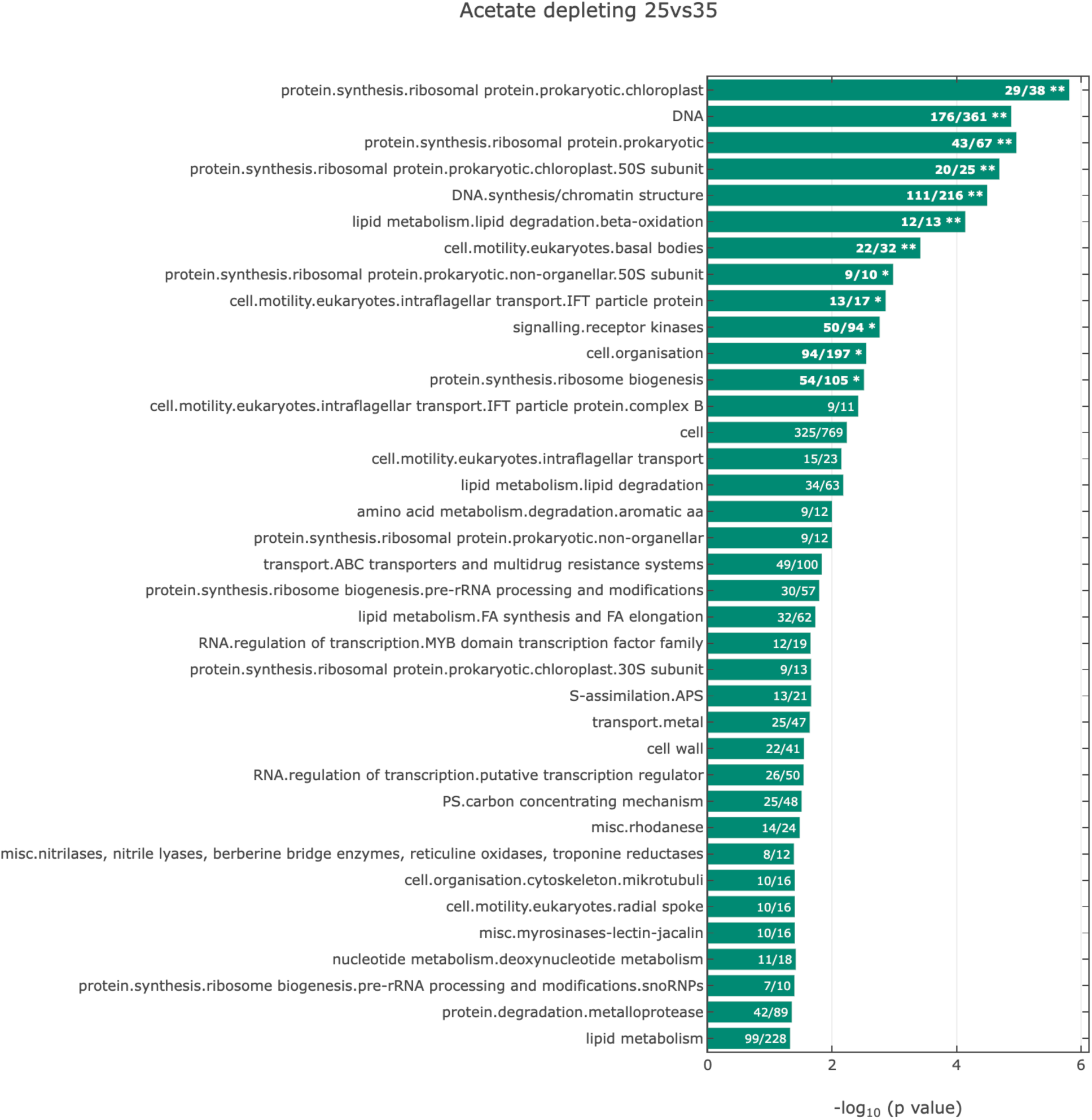
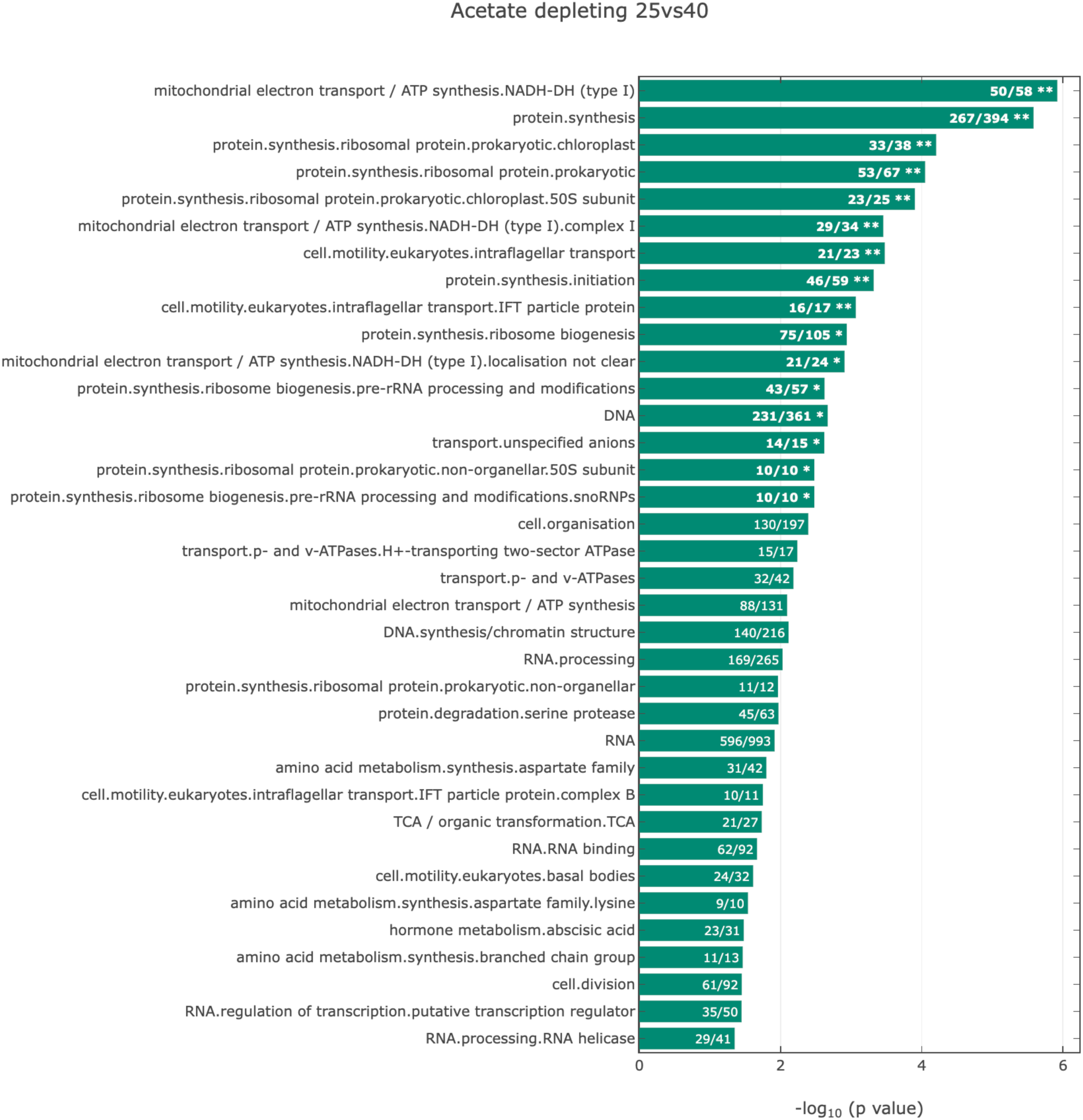
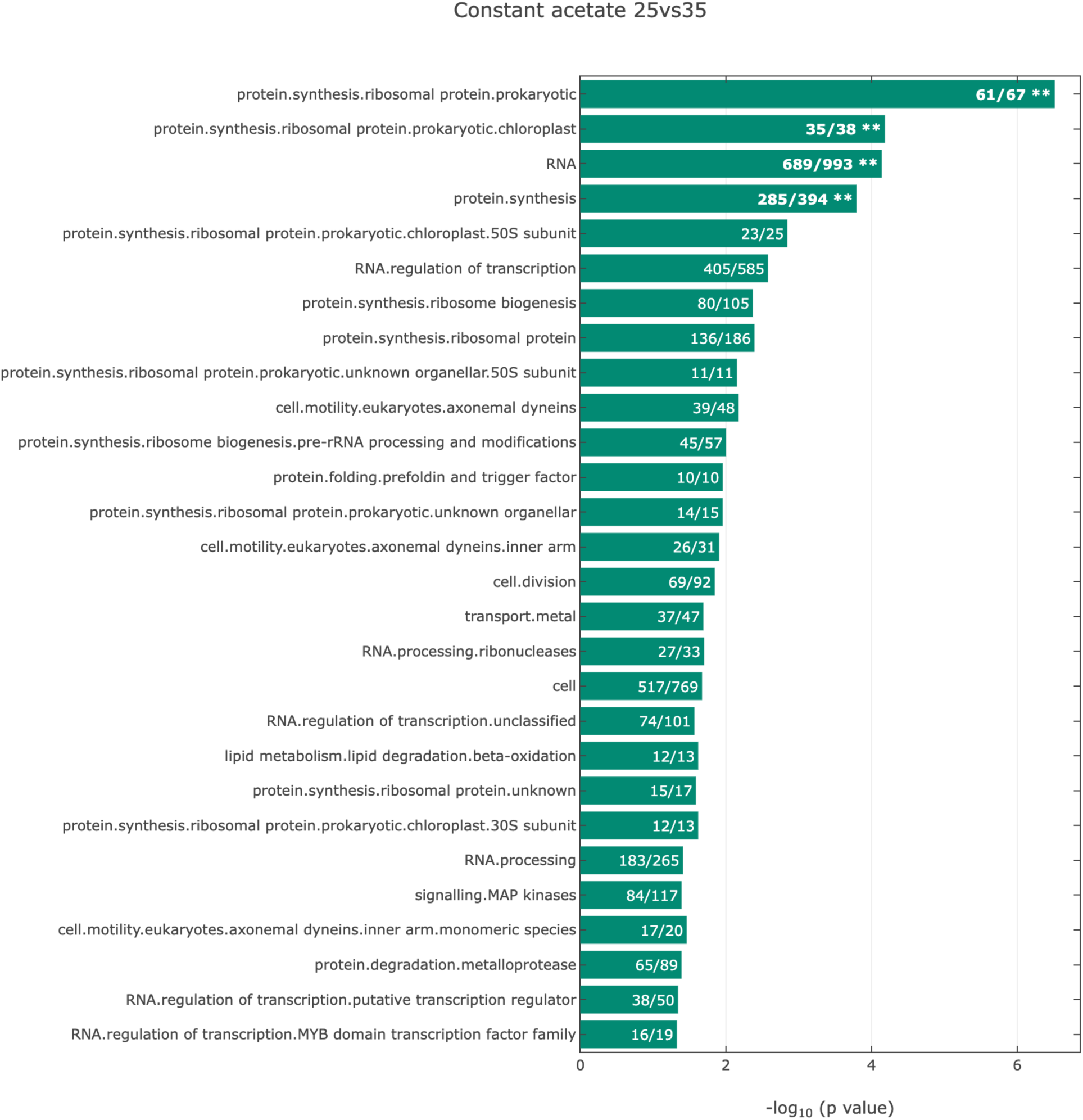
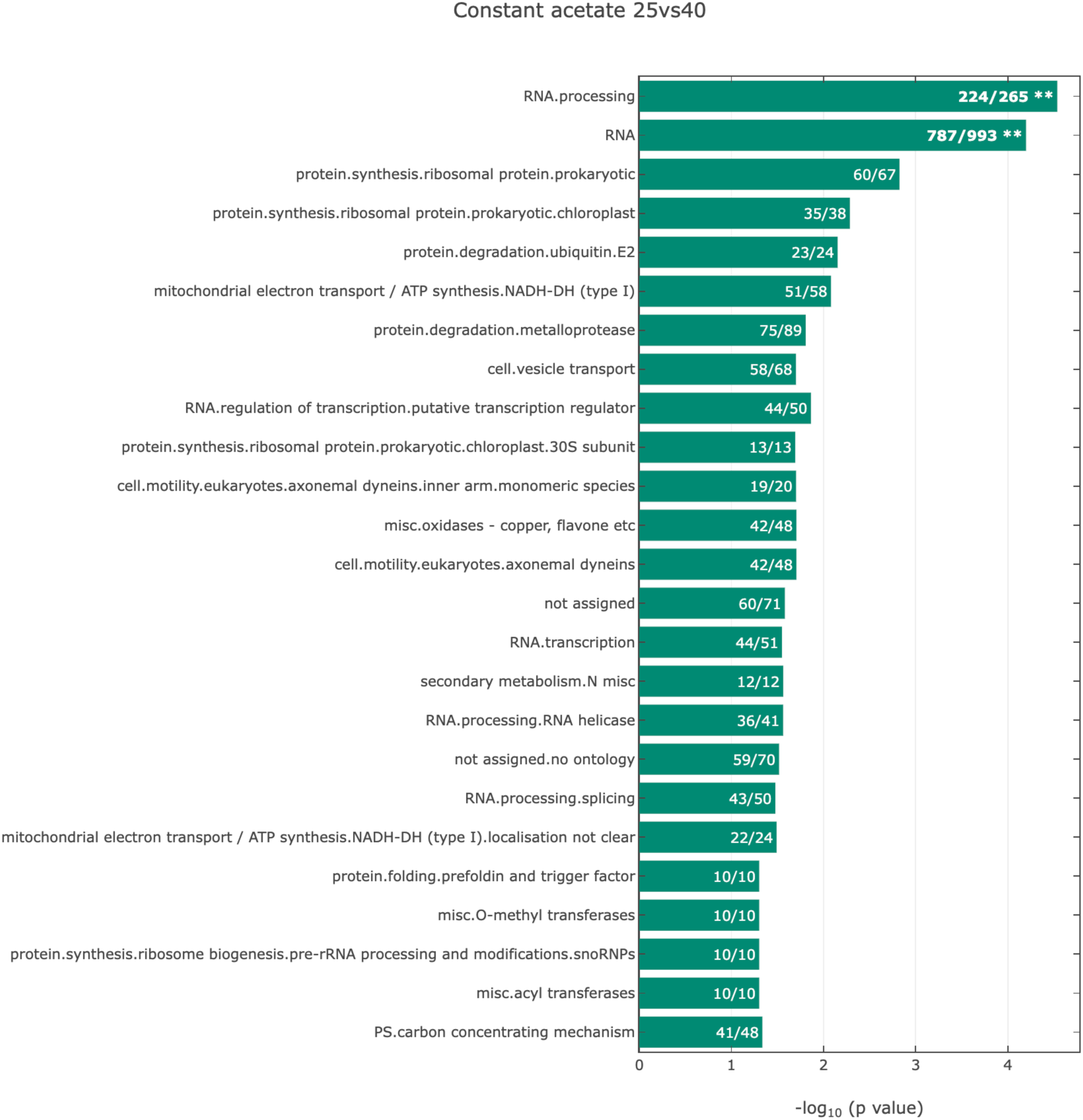
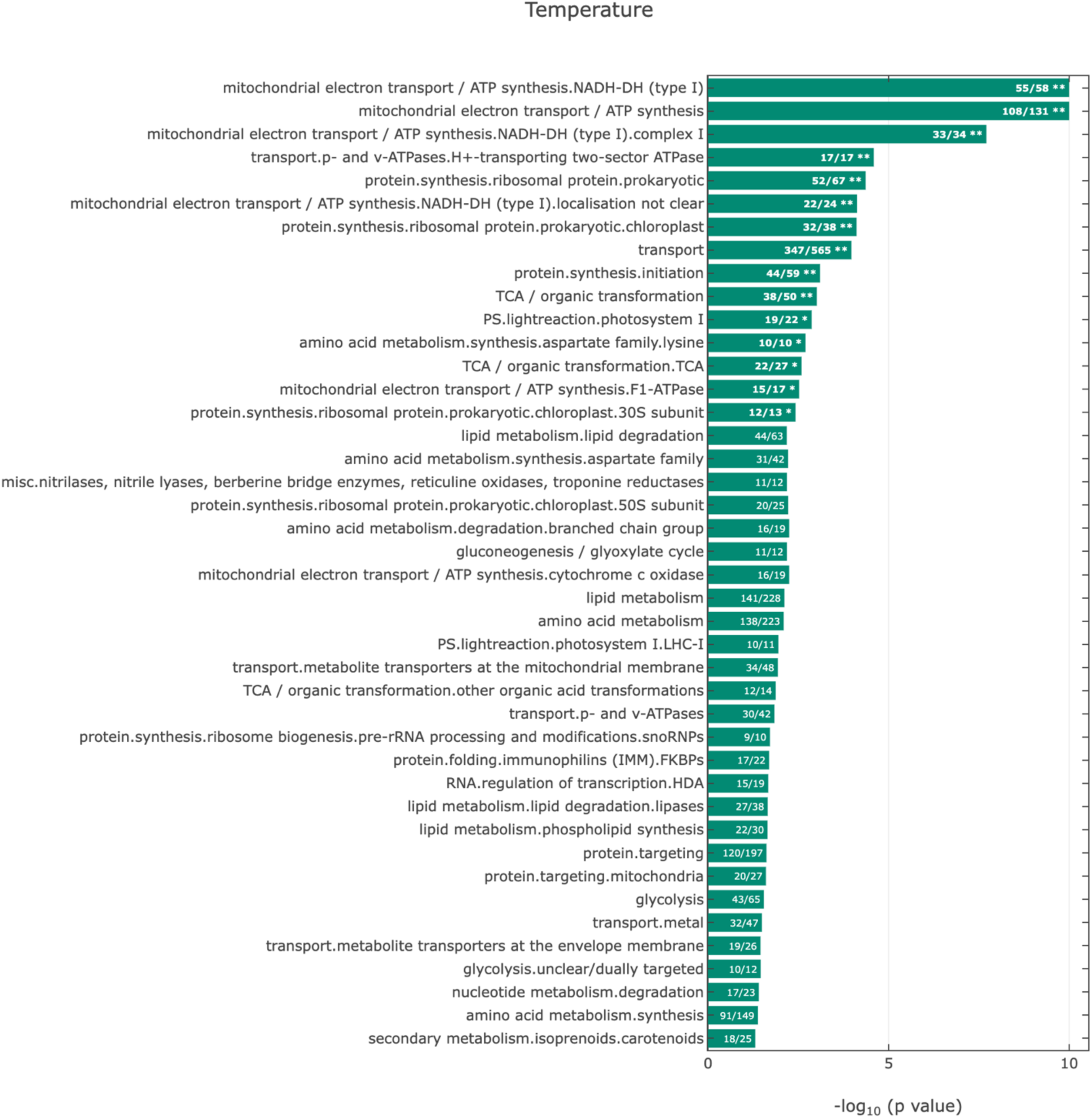
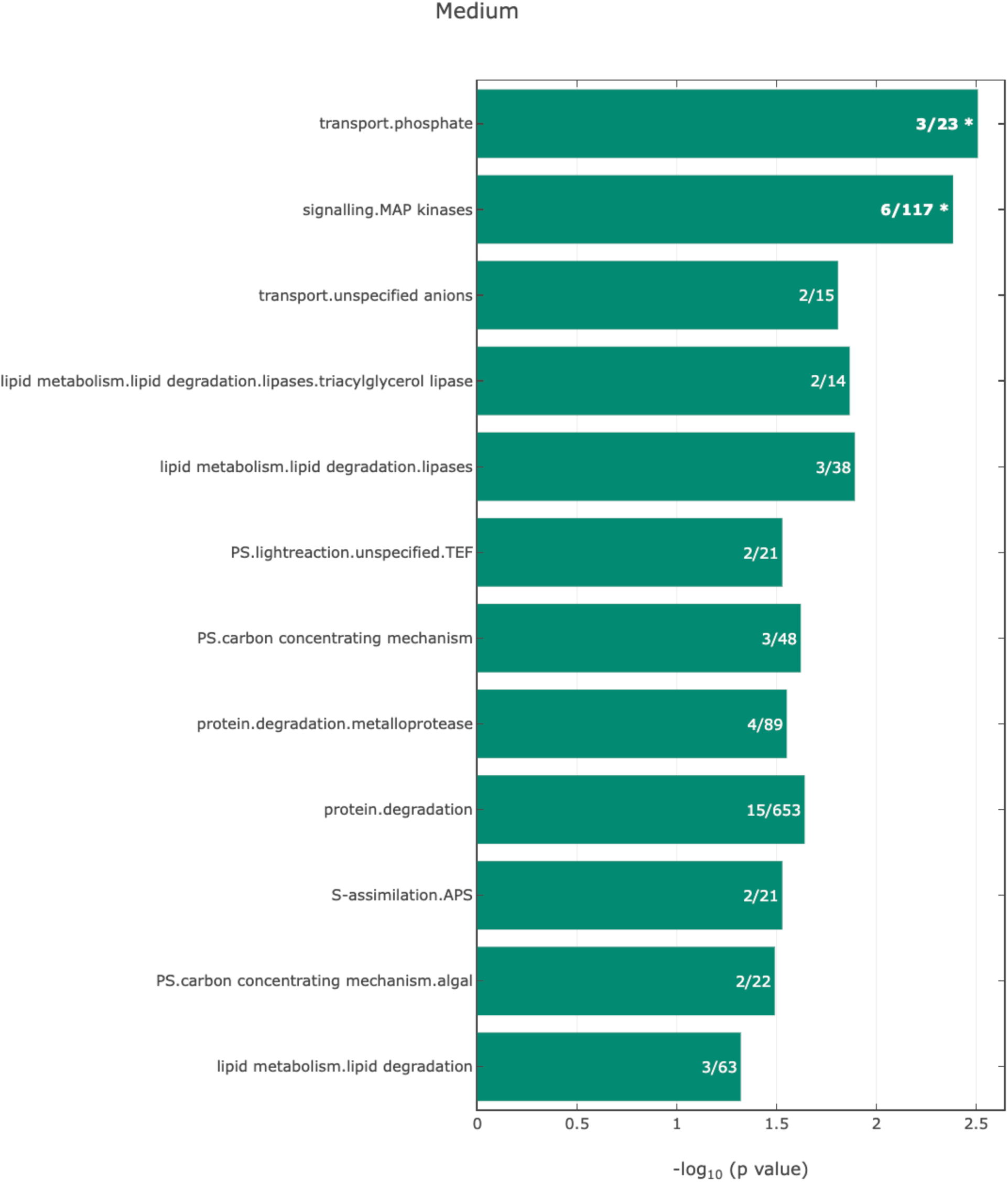
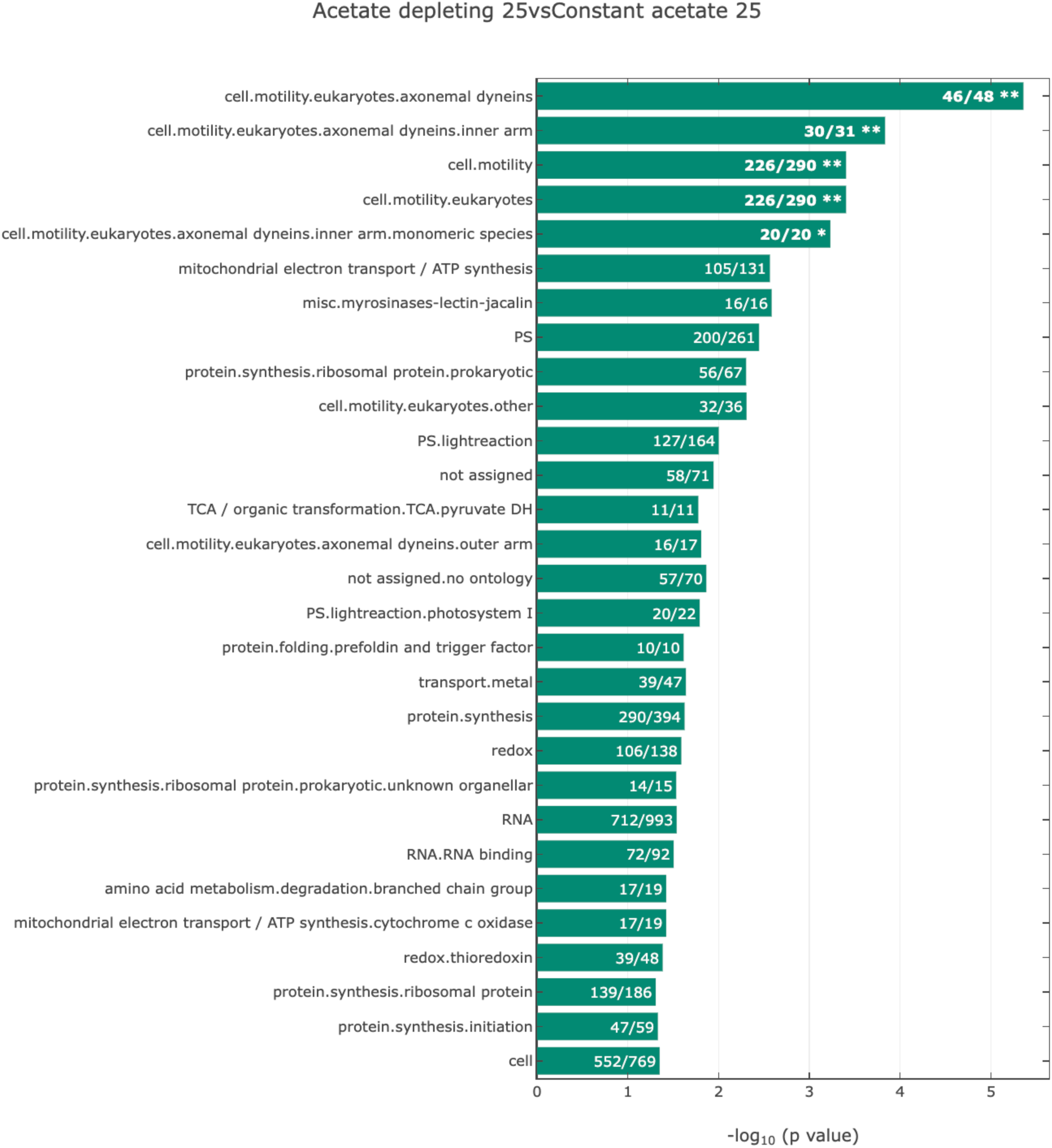
MapMan ontology enrichment revealed differentially regulated biological processes under different conditions. Ontology enrichment was performed for transcripts with significantly differential expression in the indicated comparisons using the extended MapMan ontology. The p values were calculated using a hypergeometric test. All functional sets with a p value less than 0.05 and more than 10 annotated transcripts are shown. The ratio at the end of each bar indicates the number of significant transcripts between the two compared conditions vs detected transcripts in our RNA-seq data contained in the respective function bin. Multiple testing correction was performed by Benjamini & Hochberg FDR, * FDR < 0.1, ** FDR < 0.05. The bars are sorted based on the FDR p values, from top to bottom, from the small to big FDR values. **(A)** acetate-depleting 25°C vs acetate-depleting 35°C; **(B)** acetate-depleting 25°C vs acetate-depleting 40°C; **(C)** constant-acetate 25°C vs constant-acetate 35°C; **(D)** constant-acetate 25°C vs constant-acetate 40°C; **(E)** temperature effect: (constant-acetate 35°C and acetate-depleting 35°C) vs (constant-acetate 40°C and acetate-depleting 40°C); **(F)** medium effect: (constant-acetate 35°C and constant-acetate 40°C) vs (acetate-depleting 35°C and acetate-depleting 40°C); **(G)** acetate-depleting 25°C vs constant-acetate 25°C. See interactive figures and detailed enrichment analysis in Supplemental Dataset 3.

**Supplemental Fig. 6.**
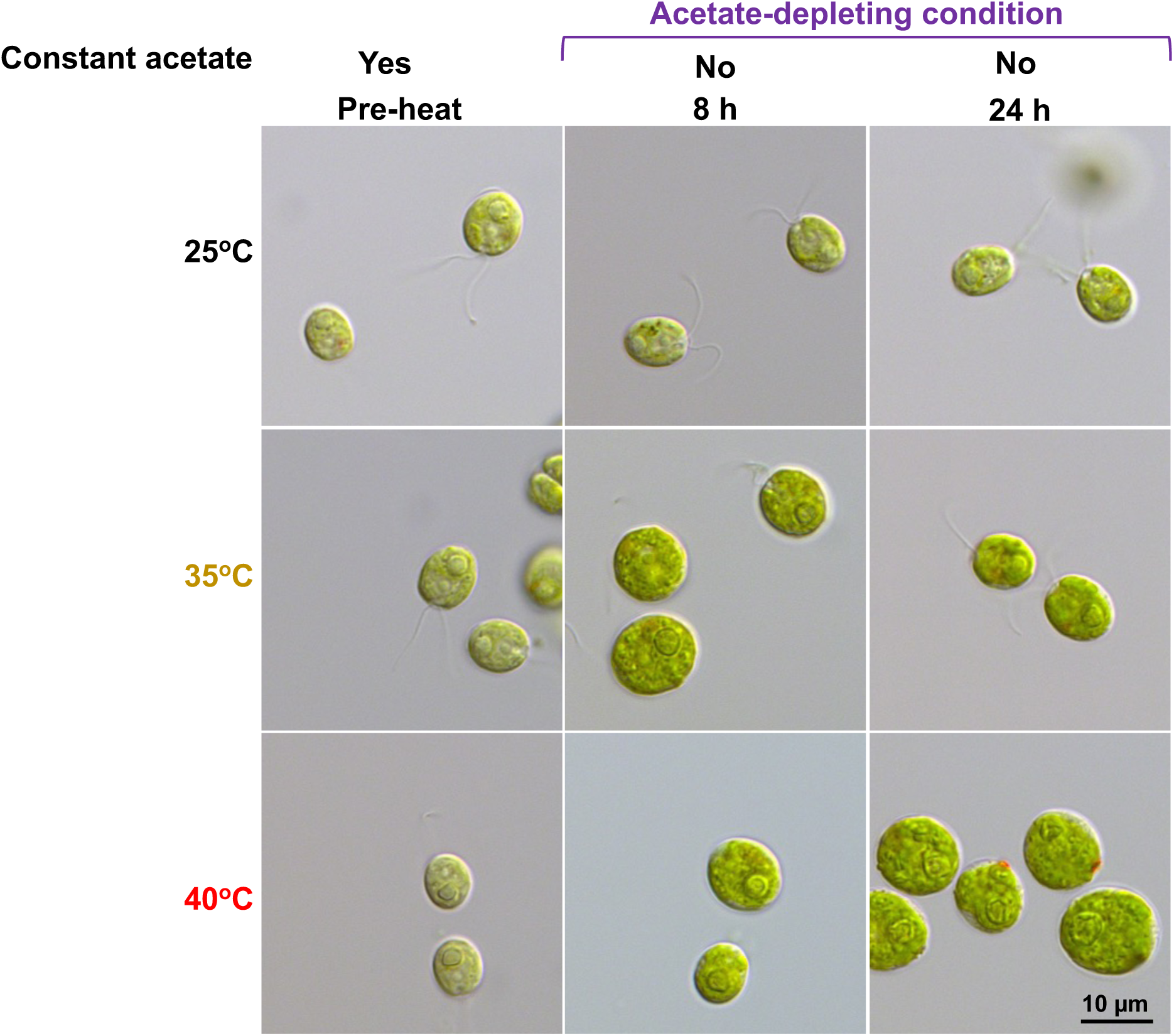
Chlamydomonas cells had different morphologies at different temperatures under the acetate-depleting condition. Light microscopic images of Chlamydomonas cells under the acetate-depleting condition. Images shown are representative results from at least three biological replicates. The algal cultivation and heat treatments were the same as in Fig. 1B.

## Notes

### Competing Interest Statement

The authors have declared no competing interest.

### Summary of Updates

We revised some of the figures, introduction, results, and discussion according to reviewers' suggestions. The main conclusion of the paper stays the same.

