## Supplemental_Dataset_1 for "Moderate High Temperature is Beneficial or Detrimental Depending on Carbon Availability in the Green Alga *Chlamydomonas reinhardtii*": CDD2_Cre12.g509600.html

 Plotly.NET DatavisualizationPlotly.NET DatavisualizationPlotly.NET DatavisualizationPlotly.NET DatavisualizationPlotly.NET DatavisualizationPlotly.NET DatavisualizationPlotly.NET DatavisualizationPlotly.NET DatavisualizationPlotly.NET DatavisualizationPlotly.NET DatavisualizationPlotly.NET DatavisualizationPlotly.NET DatavisualizationPlotly.NET Datavisualization

| Key | Value |  |
| --- | --- | --- |
| CreIdentifier | Cre12.g509600 |  |
| GeneName | CDD2 |  |
| MapMan | nucleotide metabolism.deoxynucleotide metabolism.cytosine deaminase |  |
| Annotation |  |  |
| FDR(acetate-depleting 25vs35) | 0.0189 | \* |
| FDR(acetate-depleting 25vs40) | 0.8691 |  |
| FDR(constant-acetate 25vs35) | 0.6798 |  |
| FDR(constant-acetate 25vs40) | 0.0021 | \*\* |
| FDR(medium) | 0.9807 |  |
| FDR(temperature) | 0.0058 | \*\* |
| FDR(constant-acetate 25 vs acetate-depleting 25) | 0.0169 | \* |

  

**First figure**, mean normalized transcript read counts are visualized with error bars representing standard deviation of triplicates.  
  
**Second figure**, heatmap, the color bar represents the mean normalized read count values. The dashed grey box indicates time points that were imputed from additional samples harvested from the same experiment of the constant-acetate 25°C (0 h, 0.5 h, 26 h, and 48 h, respectively). The cell physiologies were steady under the constant-acetate 25°C. See RNA-seq methods for details.  
  
**Third figure**, the transcriptional change of the transcript under each condition is presented as log2 (fold change, FC) regarding the pre-heat time point. Log2FC were determined on the mean normalized transcript abundance.  
  
Black vertical lines in the first and third figures indicate the start of heat treatments under 35°C or 40°C or the equivalent time point at 25°C. The first data point before the black line is from the pre-heat time point.
